# G protein-coupled receptor diversity and evolution in the closest living relatives of Metazoa

**DOI:** 10.1101/2025.04.15.649019

**Authors:** Alain Garcia De Las Bayonas, Nicole King

## Abstract

G protein-coupled receptors (GPCRs) play a pivotal role in the perception of environmental cues across eukaryotic diversity. Although GPCRs have been relatively well characterized in metazoans, GPCR signaling is poorly understood in their sister group, the choanoflagellates, and in other close relatives of metazoans (CRMs). Here, we examine GPCR diversity and evolution in choanoflagellates by curating a catalog of 918 GPCRs, 141 G proteins, and 367 associated regulators from 23 choanoflagellate genomes and transcriptomes. We found that the repertoire of choanoflagellate GPCRs is larger and more diverse than previously anticipated, with 18 GPCR families found in choanoflagellates, of which 12 families are newly identified in these organisms. Comparative analyses revealed that most choanoflagellate GPCR families are conserved in metazoans and/or other eukaryotic lineages. Adhesion GPCRs and a class of GPCRs fused to kinases (the GPCR-TKL/Ks) are the most abundant GPCRs in choanoflagellates. The identification of GPCR repertoires in CRMs and other non-metazoans refines our understanding of metazoan GPCR evolution and reveals the existence of previously unreported GPCR families in metazoans and at the root of the eukaryotic tree.

## INTRODUCTION

G protein-coupled receptors (GPCRs) constitute one of the largest and oldest families of receptors used to sense extracellular cues in eukaryotes (Nordström et al. 2011; Krishnan et al. 2012; De Mendoza et al. 2014). A conserved seven transmembrane (7TM) domain is a hallmark of GPCRs, while the wide spectrum of extracellular and intracellular domains in some GPCRs reflects the diversification of the gene family and its functions (Schiöth and Lagerström 2008).

For example, the extracellular N-terminus and the three extracellular loops of the 7TM domain respond to a wide range of cues, including odorant molecules, peptides, amines, lipids, nucleotides, and other molecules (Yang et al. 2021). Ligand binding triggers a conformational change in the intracellular loops, thereby activating heterotrimeric GTP binding proteins (G proteins) and additional regulators to shape downstream cellular responses (Latorraca et al. 2017). Thus, GPCRs are essential for the control of metazoan development and tissue homeostasis, and their dysregulation often leads to pathological conditions in humans, making them the most researched drug targets in the pharmaceutical industry (Hauser et al. 2017).

Based on pioneering phylogenetic analyses in vertebrates, most metazoan GPCRs have been classified into five major families: Glutamate, Rhodopsin, Adhesion, Frizzled, and Secretin (GRAFS) (Fredriksson et al. 2003; Schiöth and Fredriksson 2005; Schiöth and Lagerström 2008a). Later studies found that four of these families (Glutamate, Rhodopsin, Adhesion, and Frizzled), alongside cyclic AMP (cAMP) Receptors, GPR-107/108-like GPCRs, GPCR PIPKs, and GPR180, exist in most metazoans and in diverse other eukaryotic lineages (Bakthavatsalam et al. 2006; Kamesh et al. 2008; Nordström et al. 2008; Krishnan et al. 2012; Krishnan et al. 2014; De Mendoza et al. 2014; Van Den Hoogen et al. 2018; Mojib and Kubanek 2020; Hall et al. 2023; Luo et al. 2023). Nonetheless, the premetazoan ancestry of many metazoan GPCRs remains unclear.

With the increasing availability of genomic and transcriptomic datasets from diverse early branching metazoans (Srivastava et al. 2010; Guzman and Conaco 2016; Gold et al. 2019; Nong et al. 2020; Schultz et al. 2021; Francis et al. 2023; Santini et al. 2023; Steffen et al. 2023; Vargas et al. 2024), the closest living relatives of metazoans, the choanoflagellates (King et al. 2008; Fairclough et al. 2013; Richter et al. 2018; Brunet et al. 2019; Hake et al. 2024), and other holozoans, including members of Filasterea, Ichthyosporea, and Corallochytrea (Suga et al. 2013; Torruella et al. 2015; Grau-Bové et al. 2017; Hehenberger et al. 2017; Ocaña-Pallarès et al. 2022; Sarre et al. 2024) (Fig. 1A), we set out to refine our understanding of GPCR evolution.

**Figure 1:**
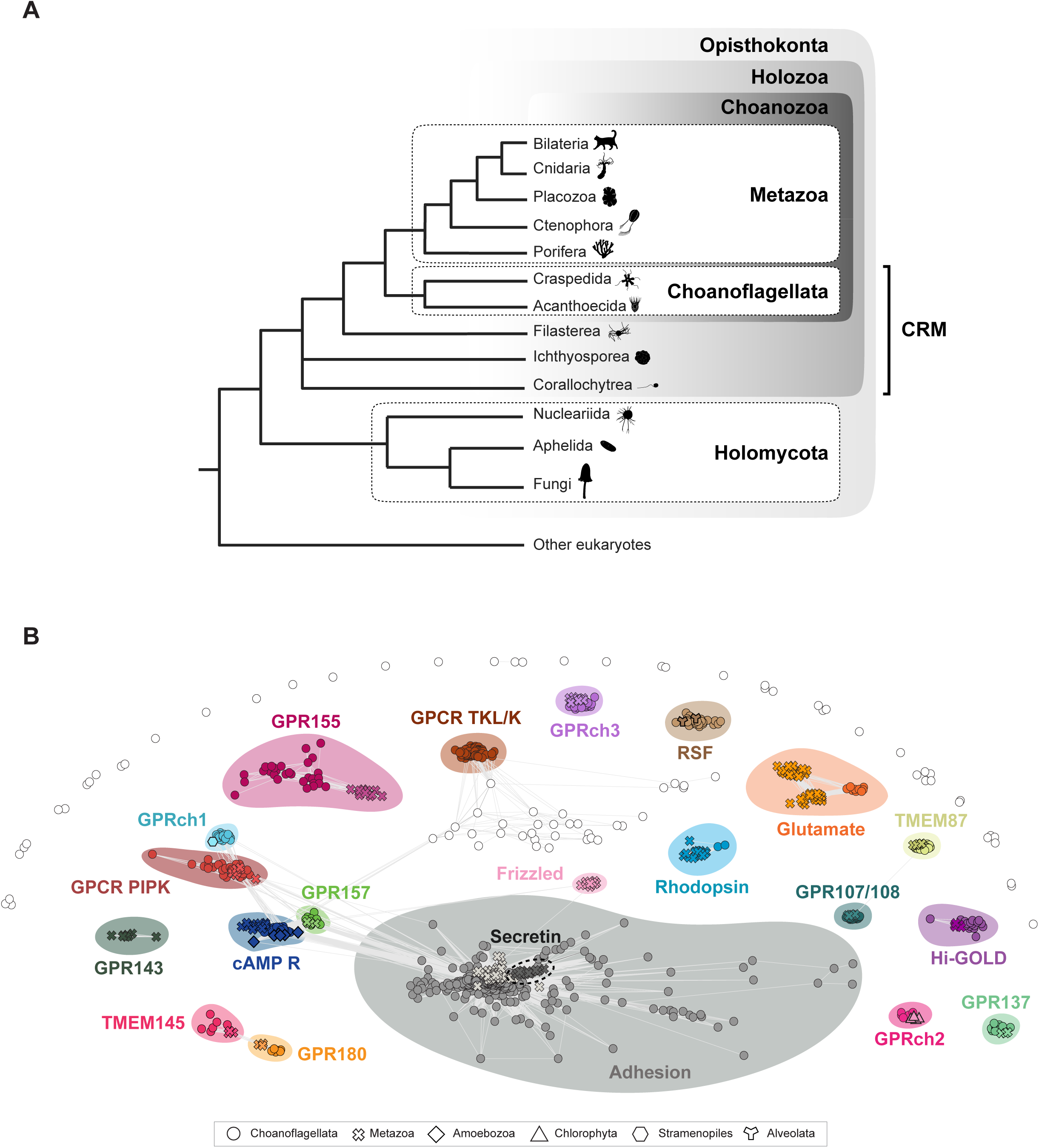
Sequence clustering reveals similarities among the 7TMs of GPCRs from choanoflagellates, metazoans, and other eukaryotes. **(A)** Choanoflagellates are the closest living relatives of metazoans. Shown is a consensus phylogeny of the major lineages analyzed in this study. We use the term “close relatives of metazoans” (CRM) to denote the paraphyletic group of non-metazoan holozoans that includes choanoflagellates, filastereans, ichthyosporeans, and corallochytreans. Organism silhouettes are from PhyloPic (http://phylopic.org/). **(B)** Most choanoflagellate GPCRs cluster with GPCRs from metazoans and other eukaryotes. The 918 choanoflagellate GPCRs (circles) identified in this study were sorted into clusters based on sequence similarity of their 7TM domains and the 7TM domains of metazoan, amoebozoan, chlorophyte, stramenopile, and alveolate GPCRs. Connecting lines (light grey) correspond to pairwise BLAST scores of p-value <1e^-6^. With the exceptions of RSF, GPCR TKL/K, GPRch1, and GPRch2, all choanoflagellate GPCR clusters contained metazoan GPCRs and, in most cases, GPCRs from other eukaryotes. No choanoflagellate GPCRs clustered with metazoan Secretin GPCRs, Frizzled GPCRs, or GPR143 GPCRs. The collection of choanoflagellate GPCRs shown as open circles did not meet the statistical threshold for designation as clusters. All the GPCR sequences used in this analysis are provided in Supplementary File 6.

To this end, we analyzed the GPCR repertoires of 23 choanoflagellate species and diverse other eukaryotes. Because of the structural modularity of GPCRs, the canonical 7TM domain on one side and the associated N-terminal and C-terminal regions on the other can follow distinct evolutionary trajectories. In addition, many receptors grouped under the same GPCR family (e.g. Rhodopsins and aGPCRs) are subdivided into numerous subfamilies. Therefore, we described and compared GPCRomes at three different levels in our analysis: (1) family- and (2) subfamily-level, based on the 7TM sequences of diverse GPCRs, and (3) associated protein domain composition, based on their respective N- and C-terminal regions.

## RESULTS

### Identification of GPCRs and downstream signaling pathway components in the proteomes of 23 diverse choanoflagellates

To catalog the diversity of GPCRs and components of the downstream signaling pathway encoded by choanoflagellates, we analyzed the genome-derived and transcriptome-derived predicted proteomes of 23 choanoflagellate species (Supplementary Table 1; King et al. 2008; Fairclough et al. 2013; Richter et al. 2018; Brunet et al. 2019; Hake et al. 2024). We first surveyed for candidate choanoflagellate GPCRs by searching choanoflagellate proteomes with Hidden Markov Models (HMMs) from the 7TM domains of 54 previously identified GPCR families (https://www.ebi.ac.uk/interpro/set/pfam/CL0192/; Fig. S1.1; Supplementary Table 2). In a complementary approach, we used an HMM that reflects the topology of the 7TM domains of all GPCRs (GPCRHMM; (Wistrand et al. 2006). These two approaches recovered 1095 and 1070 putative choanoflagellate GPCRs, respectively, of which 381 sequences were shared between the two datasets, leaving 1784 unique GPCR candidates. To remove potential false positives, highly fragmented sequences, and isoforms, the GPCR candidates were then subjected to additional filtering through homology- and topology-based approaches; 1113 sequences were removed after these filtering steps (Fig. S1.2).

Because existing 7TM HMMs may be biased toward metazoan sequences, we used the recovered choanoflagellate GPCRs to generate new choanoflagellate-derived 7TM HMMs (Supplementary file 2). To this end, the 671 validated choanoflagellate GPCRs were sorted by sequence similarity, resulting in 18 clusters (Table 1, Methods “Recovering additional choanoflagellate GPCRs using choanoflagellate GPCR BLAST queries and custom choanoflagellate GPCR HMMs” and “Clustering of the 918 validated choanoflagellate GPCRs”); the 76 GPCRs that did not assort into any of these clusters were kept for downstream analyses. We then built 7TM HMMs for each of the 18 choanoflagellate clusters and 76 additional HMMs for the unclustered GPCRs and used these custom HMMs to re-screen the 23 choanoflagellate proteomes (See Methods: “Recovering additional choanoflagellate GPCRs using choanoflagellate GPCR BLAST queries and custom choanoflagellate GPCR HMMs”). To complement this approach, we also selected GPCR sequences from each GPCR cluster and used them as BLAST queries to search for additional GPCRs in the 23 choanoflagellate proteomes (See Methods: “Recovering additional choanoflagellate GPCRs using choanoflagellate GPCR BLAST queries and custom choanoflagellate GPCR HMMs”). After filtering for fragmented sequences, isoforms, and false positives, we recovered 247 additional GPCRs that were not detected during the first round of screening. Together, our bioinformatic analysis identified a total of 918 choanoflagellate GPCRs (Supplementary file 1), with an estimated number of GPCRs per species ranging from 16 in *Barroeca monosierra* to 122 GPCRs in *Acanthoeca spectabilis* (Fig. S2).

**Table 1.** List of 18 GPCR clusters identified in choanoflagellates.

| GPCR family | Additional domain(s) | Biological function(s) | Subcellular localization | Natural ligand(s) | References |
| --- | --- | --- | --- | --- | --- |
| Rhodopsin | LRR, LDLa | light sensing, chemosensing, thermotaxis, energy metabolism, taste (✖) (★) | plasma membrane (outer segment, primary cilia), disc membranes, vacuoles (✖) (★) | light/retinal, hormones (GnRH, oestrogen, melanocortin,...), neurotransmitter (dopamine,acetylcholine,...), lipids (✖) (★) | (Pérez-Cerezales et al, 2015) (Chadha et al, 2021) (Karthikeyan et al, 2023) (Roy et al, 2020) (Cardoso et al, 2012) |
| Adhesion | GPS, EGF, LamG, Calx-beta, HRM, TSP1, EGF_CA, LRR, FN3, Kringle, CUB, MANEC, OLF, SEA, PT, etc | cell adhesion, PCP, cell migration, tissue morphogenesis, neurodevelopment, etc (★) | plasma membrane (adherens junction, FAs, dendrite, primary cilia, stereocilia), spindle, nucleus, ER (✖) | proteins of the extracellular matrix (glycosaminoglycans, collagens, laminin, ...), (✖) other membrane proteins (FLRT, neuexins, teneurins,...) | (Kusuluri et al, 2021) (Sakurai et al, 2022) (Yona et al, 2008) (Langenhan et al, 2013) (Hamann et al, 2015) (Araç and Leon, 2019) |
| Glutamate | ANF, NCD3G, EGF, Cache | synaptic transmission and neuronal excitability, cell proliferation, cell migration, phototaxis, calcium homeostasis, taste (✖) | plasma membrane (dendrite) (✖) | L-Glutamate, GABA, Glycine, sugars, peptide pheromones, ... (✖) | (Elliott and Leys, 2010) (Crupi et al, 2019) (Niswender and Conn, 2010) (Wong et al, 2022) (Sigel and Steinmann, 2012) (Qian et al, 2023) (Dubovski et al, 2022) (Silva and Antunes, 2017) (Laboute et al, 2023) |
| cAMP | none | cell chemotaxis, cell aggregation, cell differentiation, cyst formation (★) | plasma membrane (★) | cAMP (★) | (Kawabe et al, 2009) (Louis et al, 1994) (Saxe et al, 1993) (Klein et al, 1988) |
| GPR137 | none | cell proliferation and apoptosis, neuronal differentiation (✖) | lysosome (✖) | unknown | (Gan et al, 2019) (Men et al, 2018) |
| GPR155 | PIN-like transporter, DEP | nutrient sensing, energy metabolism, tissue patering (✖) | lysosome (✖) | cholesterol (✖) | (Shin et al, 2022) (Bayly-Jones et al, 2023) (Schöneberg, 2024) (Wang et al, 2018) |
| GPR157 | none | neuronal differentiation (✖) | plasma membrane (primary cilia) (✖) | unknown | (Takeo et al, 2016) |
| GPCR-TKL/K | Tyrosine kinase-like / kinase | unknown | unknown | unknown | this study |
| TMEM145 | GOLD | structural integrity of hair cell stereocilia (✖) | plasma membrane (stereocilia) (✖) | unknown | (Roh et al, 2025) |
| TMEM87 | GOLD | protein trafficking, Golgi-Ph maintenance (✖) | Golgi (✖) | unknown | (Hoel et al, 2022) (Hirata et al, 2015) (Shin et al, 2020) |
| GPR107/108 | GOLD | protein trafficking, recycling of receptors, (✖) immunity | Golgi, nucleus, clathrin vesicles (✖) | neurostatin, gambogic acid (✖) | (Yang et al, 2024) (Yosten et al, 2012) (Zhou et al, 2014) (Lyu et al, 2022) (Balazova et al, 2021) |
| GPR180 | GOLD | gametogenesis (★), lipid metabolism (✖) | plasma membrane, vesicles (✖) | L-lactate, CTHRC1 (✖) | (Yoshida et al, 2023) (Wang et al, 2022) (Mosienko et al, 2018) |
| Hi-GOLD | GOLD | unknown | unknown | unknown | this study |
| GPCR PIPK | PIPK | cell density sensing, bacterial defense, phagocytosis, asexual development, chemotaxis (★) | endosomal vesicles, phagosome (★) | unknown | (Van Den Hoogen et al, 2018) (Bakthavatsalam et al, 2006) (Bakthavatsalam et al, 2007) (Hua et al, 2013) (Riyahi et al, 2011) |
| RSF | none | unknown | unknown | unknown | this study |
| GPRch1 | none | unknown | unknown | unknown | this study |
| GPRch2 | none | unknown | unknown | unknown | this study |
| GPRch3 | none | unknown | unknown | unknown | this study |

✖ Metazoa ▲ Embryophyta ★ Protozoa
GRAFS
GOST

To search for downstream signaling components of GPCRs, we aligned the 23 choanoflagellate proteomes with HMMs specific to the three classes of heterotrimeric G proteins (Gα, Gβ, and Gγ) (Neves et al. 2002; Dupré et al. 2009; Wootten et al. 2018), positive regulators (Ric8, Phosducin) (Willardson and Howlett 2007; Srivastava et al. 2019), and negative regulators (GRK, Arrestin, RGS, GoLoco) (Willard et al. 2004; Rajagopal and Shenoy 2018; Bagnato and Rosanò 2019; Gurevich and Gurevich 2019; O’Brien et al. 2019) of G protein signaling (Fig. S3 and Supplementary Table 2). Our analysis recovered 141 heterotrimeric G proteins, 308 positive regulators, and 59 negative regulators of G protein signaling (Supplementary text and Supplementary files 3, 4, and 5). Together with the abundant GPCRs detected in choanoflagellates, the finding of nearly complete G protein signaling pathways suggests that choanoflagellates engage in canonical G protein signaling.

To assess the predictive power of our protein-detection pipeline, we then compared the new GPCR and cytosolic signaling component datasets from two choanoflagellates – *Salpingoeca rosetta* and *Monosiga brevicollis* – with previously published GPCR and downstream GPCR signaling component counts for these two species (Nordström et al. 2009a; Krishnan et al. 2012; De Mendoza et al. 2014; Krishnan et al. 2015; Lokits et al. 2018). All 11 GPCRs previously described in *M. brevicollis* and all 14 GPCRs found in *S. rosetta* were present in our dataset, along with 10 and 9 newly identified GPCRs, respectively. Similarly, all previously identified heterotrimeric G proteins and diverse positive and negative regulators of G protein signaling from these two choanoflagellates were in our dataset, along with one newly identified Gβ subunit, one RGS, one Goloco, and one additional Arrestin in *M. brevicollis* (Fig.S3; Supplementary file 3). Due to the stringency of our filtering approach and the use of transcriptome-derived proteomes for a majority of choanoflagellate species, these numbers still likely underestimate total GPCR and downstream component abundance and diversity in choanoflagellates. Moreover, in the case of transcriptome-derived proteomes, failure to detect homologs could reflect false negatives rather than a real absence of the gene families.

Conversely, sequence contamination and splice isoforms could possibly account for false positives.

### Identification of 18 GPCR families in choanoflagellates

The conservation and alignability of the 7TM domain enable the straightforward categorization of GPCRs into families across eukaryotic diversity (Attwood and Findlay 1993; Fredriksson et al. 2003; Schiöth and Fredriksson 2005). Therefore, to investigate the relationships among choanoflagellate GPCRs, we started by extracting and clustering their 7TM sequences. In addition, we included 7TM sequences from diverse metazoans, amoebozoans, stramenopiles, alveolates, and chlorophytes in our analysis to aid in the identification of the choanoflagellate GPCR families (see Methods “Clustering of the 918 validated choanoflagellate GPCRs”; Supplementary file 6).

Through all-against-all pairwise comparisons of the 7TM domains of the 918 choanoflagellate GPCRs recovered in the previous analysis (Fig. S1.3c) with 7TMs from diverse metazoans, amoebozoans, stramenopiles, alveolates, and chlorophytes, we found that most choanoflagellate GPCRs belong to 18 classes of GPCRs (Fig. 1B). Six of these classes were previously detected in *S. rosetta* and/or *M. brevicollis*: Glutamate, Adhesion, GPR107/108, cAMP, GPR137, and GPCR_PIPK (Figs. 1B, 2, and S4; Table 1; Krishnan et al. 2012; De Mendoza et al. 2014; Van Den Hoogen et al. 2018; Gan et al. 2019)).

**Figure 2.**
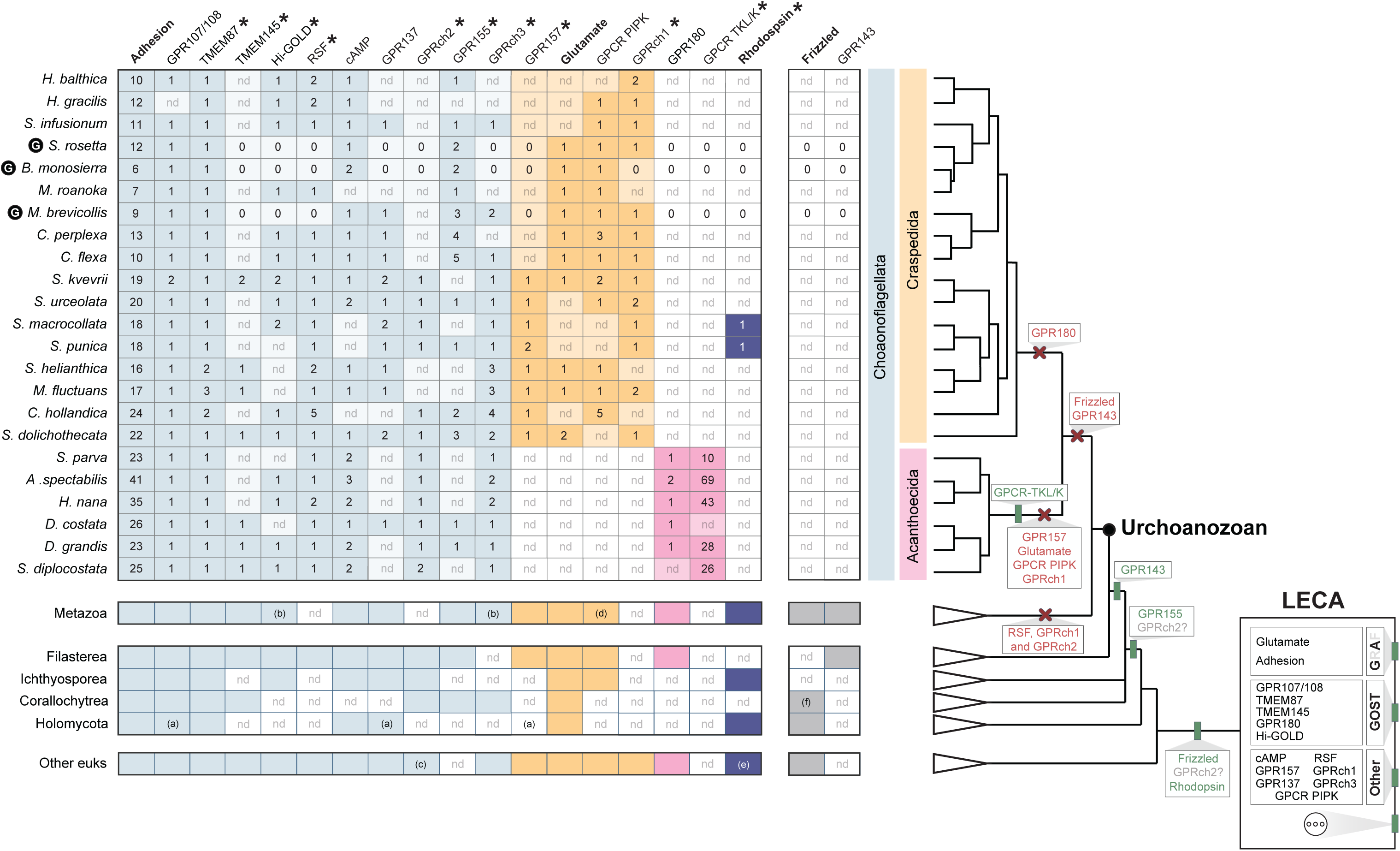
Evolutionary history of GPCR families detected in choanoflagellates, metazoans, and other eukaryotes. **(Left)** Taxonomic distribution of GPCR families. Shown in the table are the numbers of GPCR family members (columns) detected in each lineage (rows). The GPCR families are grouped based on their inferred phylogenetic distributions in choanoflagellates: pan-choanoflagellate GPCRs (light blue), GPCR families found in craspedids but not acanthoecids (orange), GPCR families found in acanthoecids but not craspedids (pink), and GPCRs found only in *S. macrocollata* and *S. punica* (dark blue). Adhesion, Glutamate, Rhodopsin, and Frizzled, four of the five GRAFS GPCR families, are indicated in bold. Asterisks indicate GPCR families that were not known to exist in choanoflagellates prior to this study. For those species in which only transcriptomes are available, gene numbers represent a minimum. nd = not detected in lineages for which only transcriptome data are available. For species with both transcriptome and genome data (*S. rosetta*, *B. monosierra*, and *M. brevicollis*; G enclosed within a black circle) (King et al. 2008; Fairclough et al. 2013; Hake et al. 2024), failure to detect a GPCR subfamily member is indicated with a “0”. (a) only found in nucleariids, (b) lost in vertebrates, (c) only found in chlorophytes, (d) only found in sponges, (e) only found in amoebozoans, (f) only found in *Syssomonas multiformis*. **(Right)** A consensus phylogeny shows the relationships among the 23 choanoflagellates included in this study, metazoans, filastereans, ichthyosporeans, corallochytreans, holomycotans, and diverse other eukaryotes. The inferred origins (vertical green rectangle) and subsequent losses (red cross) of GPCR families are indicated at relevant branches on the consensus phylogeny. GPCR families inferred to have originated in the Last Eukaryotic Common Ancestor (LECA) are represented at the root of the phylogeny (box). The presence of additional GPCR families in LECA, not covered in our study, is depicted by three dots. “GA” indicates two of the GRAFS GPCR families: Glutamate and Adhesion. “GOST” indicates subfamilies of GOST GPCRs. Additional GPCR families are listed under “Other.” Uncertainty about the ancestry of GPRch2 is indicated with a question mark.

This left twelve new GPCR families that had not, to our knowledge, been previously detected in choanoflagellates: Rhodopsin, TMEM145, GPR180, TMEM87, GPR155, GPR157, and six additional GPCR families that appear to fall outside all previously characterized GPCR families in eukaryotes. For reasons that will be discussed further below, we have named these six new GPCR families “Rémi-Sans-Famille” (RSF), “Hidden Gold” (Hi-GOLD), GPCR-TKL/K, GPRch1, GPRch2, and GPRch3. (Fig. 1B; Table 1). An additional 76 choanoflagellate GPCRs did not cluster above threshold with any other GPCRs detected from choanoflagellates or other eukaryotes.

### Overview of choanoflagellate GPCR family evolution

Of the 18 GPCR families identified in choanoflagellates (Table 1), 16 were found, based strictly on the similarity of their signature 7TM domains, in diverse holozoan and non-holozoan eukaryotes and likely evolved in stem eukaryotes (Fig. 2; Supplementary File 7). GPR155 was only detected in metazoans and close relatives of metazoans (CRMs; Fig. 1A), supporting an origin in stem holozoans. The GPCR-TKL/K family was only detected in acanthoecid choanoflagellates, suggesting it evolved in the acanthoecid stem lineage.

Although Adhesion, GPR107/108, TMEM87, TMEM145, Hi-GOLD, RSF, cAMP, GPR137, GPR155, and GPRch3 GPCRs were detected in a wide range of choanoflagellates, other GPCR families appeared to be restricted to one or the other of the two choanoflagellate orders: craspedids (GPR157, Glutamate, GPCR PIPK, and GPRch1) or acanthoecids (GPR180 and GPCR TKL/K; Fig.2). The phylogenetic distributions of these GPCR families are likely the result of secondary losses in choanoflagellates because homologs of these GPCRs are detected in other eukaryotes.

Two additional choanoflagellate GPCR families evolved in stem eukaryotes but were lost from metazoans and some CRMs: RSF GPCRs and GPRch1. Frizzled GPCRs, which possibly originated in the last common ancestor of opisthokonts and amoebozoans (Krishnan et al. 2012; De Mendoza et al. 2014), and GPR143, which emerged in stem holozoans (Fig.2; De Mendoza et al. 2014), were both lost from stem choanoflagellates.

Metazoan-like Rhodopsins (as defined by GPCRs that are significant matches to the 7tm_1(PF0001) HMM (Clarke et al. 2023), possess an eight helix (Krishna et al. 2002; Kock et al. 2009; Knepp et al. 2012; Sensoy and Weinstein 2015), and present characteristic motifs such as aspartate (D) in TM2, E/DRY in TM3, CWxP in TM6, or NPxxY in TM7 (Krishnan et al. 2012; Rinne et al. 2019)) were detected in two choanoflagellate sister species ̶ *Salpingoeca macrocollata* and *Salpingoeca punica* (Richter et al. 2018; López-Escardó et al. 2019; Ginés-Rivas and Carr 2025a). The detection of metazoan-like Rhodopsins in metazoans, choanoflagellates, ichthyosporeans, fungi, and amoebozoans suggests that these receptors predate the split between amoebozoans and opisthokonts (Fig. 2).

### Metazoan GPCR families that originated in stem eukaryotes

#### Glutamate Receptors

The Glutamate Receptor family, which includes important regulators of neuronal excitability, synaptic transmission, and taste recognition in metazoans (Pin et al. 2003; Schiöth and Lagerström 2008a; Chun et al. 2012), first evolved in stem eukaryotes (Fig. 2; (Krishnan et al. 2012; De Mendoza et al. 2014)). In addition to their 7TM domain, Glutamate Receptors have a large, structured extracellular region that varies in different members of the family (Fig. 3A; (Ellaithy et al. 2020a)). For example, mGluR/T1R/CaSR receptors exhibit a bi-lobed ligand-binding domain (ANF) directly N-terminal to a cysteine-rich domain (NCD3G) (Koehl et al. 2019), both of which are essential for receptor activity (Hu et al. 2000; Jiang et al. 2004; Rondard et al. 2006). In GABA_B_ receptors, the ANF domain is conserved while the NCD3G is absent (Evenseth et al. 2020; Papasergi-Scott et al. 2020). Finally, mGlyR/GPR179 receptors present a distinct domain organization, with a ligand-binding Cache domain sitting atop an EGF-like domain (Jeong et al. 2021; Laboute et al. 2023; Rosenkilde and Mathiesen 2023; Yun et al. 2024).

**Figure 3:**
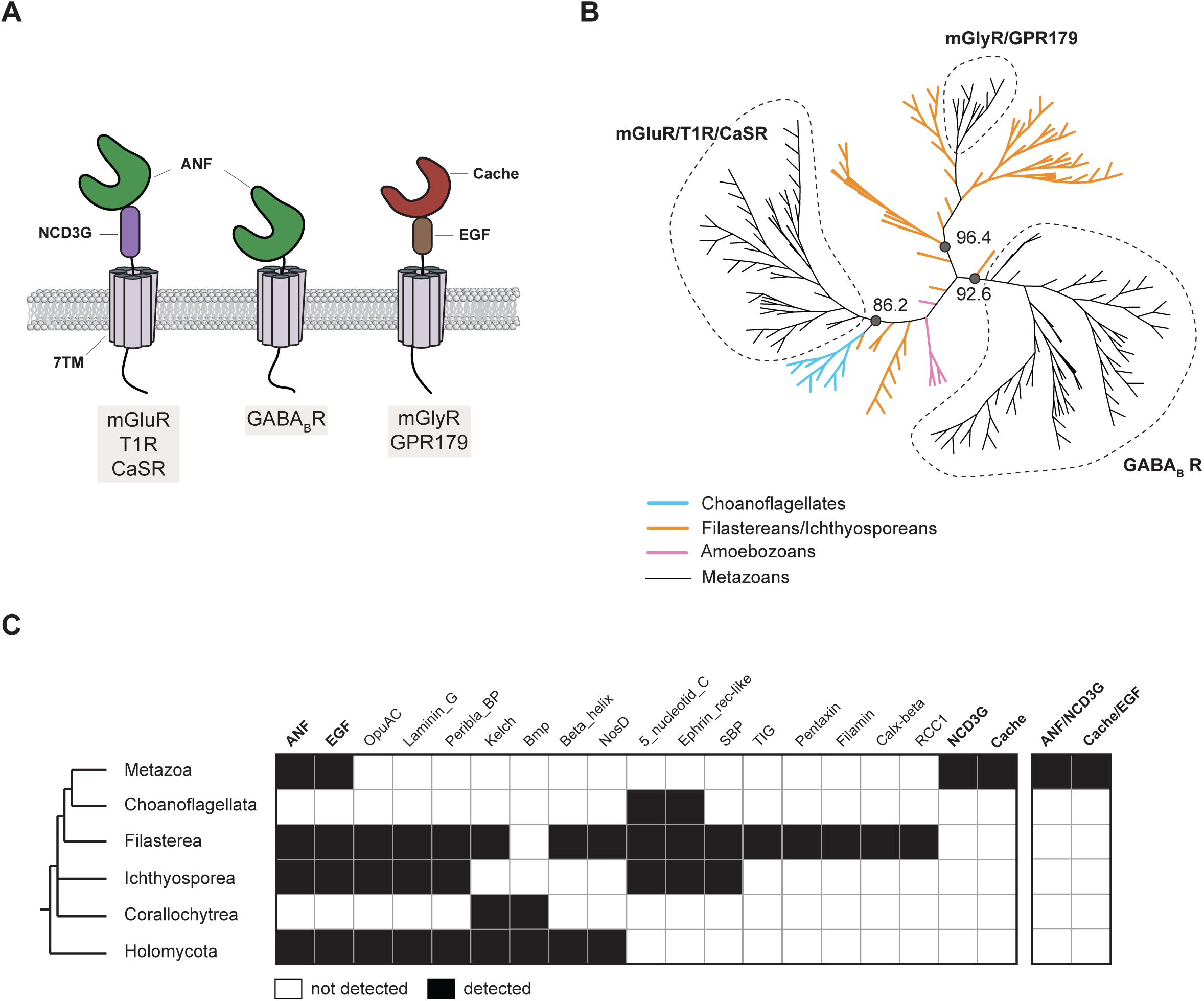
Evolution of Glutamate Receptor protein domain architecture. **(A)** Schematic representation of the main protein domains found in metazoan the Glutamate Receptor family. GluR/T1R/CaSR receptors have a conserved extracellular module containing an ANF ligand-binding domain fused to a cysteine-rich NCD3G domain. GABA_B_R GPCRs contain only the ANF ligand binding domain. mGlyR/GPR179 GPCRs contain a Cache ligand-binding domain fused to an EGF-like domain. **(B)** Phylogenetic analysis of the 7TM domains of metazoan, choanoflagellate, filasterean, ichthyosporean, and amoebozoan Glutamate Receptors yielded three well-supported clades of GPCRs corresponding to the receptors depicted in panel A. Filasteran and ichthyosporean sequences (orange) clustered as the sister group to metazoan mGluR/CaSR/T1R, mGlyR/GPR179, and GABA_B_ GPCRs, or branch separately from the rest of GPCRs used in this analysis. Choanoflagellate sequences (light blue) branch as the sister group to the metazoan mGluR/CaSR/T1R GPCRs. No amoebozoan sequence (pink) clustered with metazoan or CRMs. UFboot support values are indicated for each three major nodes of this unrooted maximum-likelihood phylogenetic tree. All the sequences used to build this phylogeny are listed in Supplementary Files 8 and 9. **(C)** Phylogenetic distribution of ECDs detected in Glutamate Receptors from diverse opisthokonts. While the ANF and EGF protein domains evolved before the diversification of opisthokonts, the presence of the NCD3G and Cache protein domains in GPCRs was not detected in any non-metazoan. Similarly, the ANF/NCD3G and Cache/EGF protein domain modules were only detected in metazoans. Glutamate Receptors from non-metazoans contain diverse extracellular protein domains that are not found in metazoan GPCRs. Protein domains (left) or modules containing pairs of protein domains (right) are indicated in the columns. Taxonomic distribution is indicated in the rows. Protein domains found in metazoan Glutamate Receptors – ANF, EGF, NCD3G, and Cache – are indicated in bold. See Supplementary File 10 for a list of all Glutamate Receptors screened in this analysis.

While members of the Glutamate family and their canonical domain architectures are conserved in a wide range of metazoans, including sponges and ctenophores (Krishnan et al. 2014; De Mendoza et al. 2014; Krishnan and Schiöth 2015), it is presently unclear how CRM Glutamate Receptors identified by clustering of their 7TM domains relate to their metazoan counterparts.

To better understand the evolution of Glutamate Receptors, we conducted a phylogenetic analysis of the 7TM domains from metazoan and CRM Glutamate Receptors (Fig. 3B and Supplementary files 8 and 9). All choanoflagellate Glutamate Receptors identified in our study formed the sister group to metazoan mGluR/T1R/CaSR GPCRs. The Glutamate Receptors of filastereans and ichthyosporeans branched either at the base of metazoan mGluR/T1R/CaSR receptors, mGlyR/GPR179 receptors, or GABA_B_ receptors; or at positions on the tree that were distinct from the metazoan Glutamate Receptor families. The Glutamate Receptors from the amoebozan *Dictyostelium discoideum*, of which at least one, GrlE, binds both GABA and Glutamate presumably through its conserved ANF domain (Anjard and Loomis 2006; Taniura et al. 2006; Wu and Janetopoulos 2013), grouped separately from metazoan and CRM GPCRs in our analysis.

Next, we assessed the protein domain compositions of the extracellular regions of Glutamate Receptors from diverse opisthokonts (Fig. 3C and Supplementary file 10). Although earlier studies reported no extracellular domains in choanoflagellate Glutamate Receptors (Krishnan et al. 2012; De Mendoza et al. 2014), our analysis revealed the presence of 5’-nucleotidase C-terminal domains (5_nucleotid_C) and Ephrin receptor-like domains (Ephrin_rec-like) that are also found in Glutamate Receptors from other CRMs, but not in metazoans.

Two domains present in the N-termini of some metazoan Glutamate Receptors – ANF and EGF – were also found in Glutamate Receptors from other opisthokonts, suggesting an ancient association of these domains with the signature Glutamate 7TM. On the contrary, no non-metazoan GPCRs yet identified possess the canonical metazoan mGluR/T1R/CaSR architecture of an ANF_receptor domain coupled with an NDC3G domain or the mGlyR/GPR179 architecture of a Cache domain coupled with an EGF domain. Together, our data suggest that while the 7TM domains of the three main classes of metazoan Glutamate Receptors ̶ mGluR/T1R/CaSR, mGlyR/GPR179, and GABA_B_ receptors ̶ have a pre-metazoan origin, the stereotypical combination of domains in their N-terminus was a stem metazoan innovation.

#### Rhodopsins

Rhodopsins are the largest GPCR family in most metazoans and include receptors for hormones, neuropeptides, neurotransmitters, nucleotides, and light, among others (Fredriksson et al. 2003; Schiöth and Lagerström 2008; De Mendoza et al. 2014; Pándy-Szekeres et al. 2018). Orthologs of bilaterian Rhodopsins have been found in Cnidaria and Ctenophores but have not been detected in sponges (Feuda et al. 2014; Krishnan et al. 2014; Thiel et al. 2023). While Rhodopsins expanded and diversified in metazoans (Fredriksson and Schiöth 2005; De Mendoza et al. 2014; Thiel et al. 2023), the origin of this family might have predated the emergence of metazoans, as homologs have been detected in Fungi and in other eukaryotes (Fig. 2; Krishnan et al. 2012; De Mendoza et al. 2014).

Upon searching the choanoflagellate genomes and transcriptomes with an HMM profile built solely on the aligned 7TM domains from metazoan Rhodopsin sequences (7TM_1; Supplementary Table 2), we identified two previously undetected choanoflagellate Rhodopsins. Additionally, we detected a metazoan-like Rhodopsin from the ichthyosporean *Pirum gemmata* (Fig. 2 and Supplementary file 11).

All-against-all pairwise comparison of the choanoflagellate and ichthyosporean Rhodopsins with the complete Rhodopsin repertoires of representative metazoans (6149 Rhodopsins in total) revealed that choanoflagellate Rhodopsins are most similar to metazoan opsins (including arthropsin, cephalochordate Go opsin, placozoan placopsin, vertebrate opsin 3, and Teleost multiple tissue opsins) and, to a lesser extent, SOG receptors (Fig. 4A, B and Supplementary files 11 and 12; Fredriksson et al. 2003; Kamesh et al. 2008; Nordström et al. 2008; McVeigh et al. 2018; Yañez-Guerra et al. 2022). No statistically significant metazoan or choanoflagellate hits were detected for the predicted ichthyosporean Rhodopsin (Supplementary file 12).

**Figure 4:**
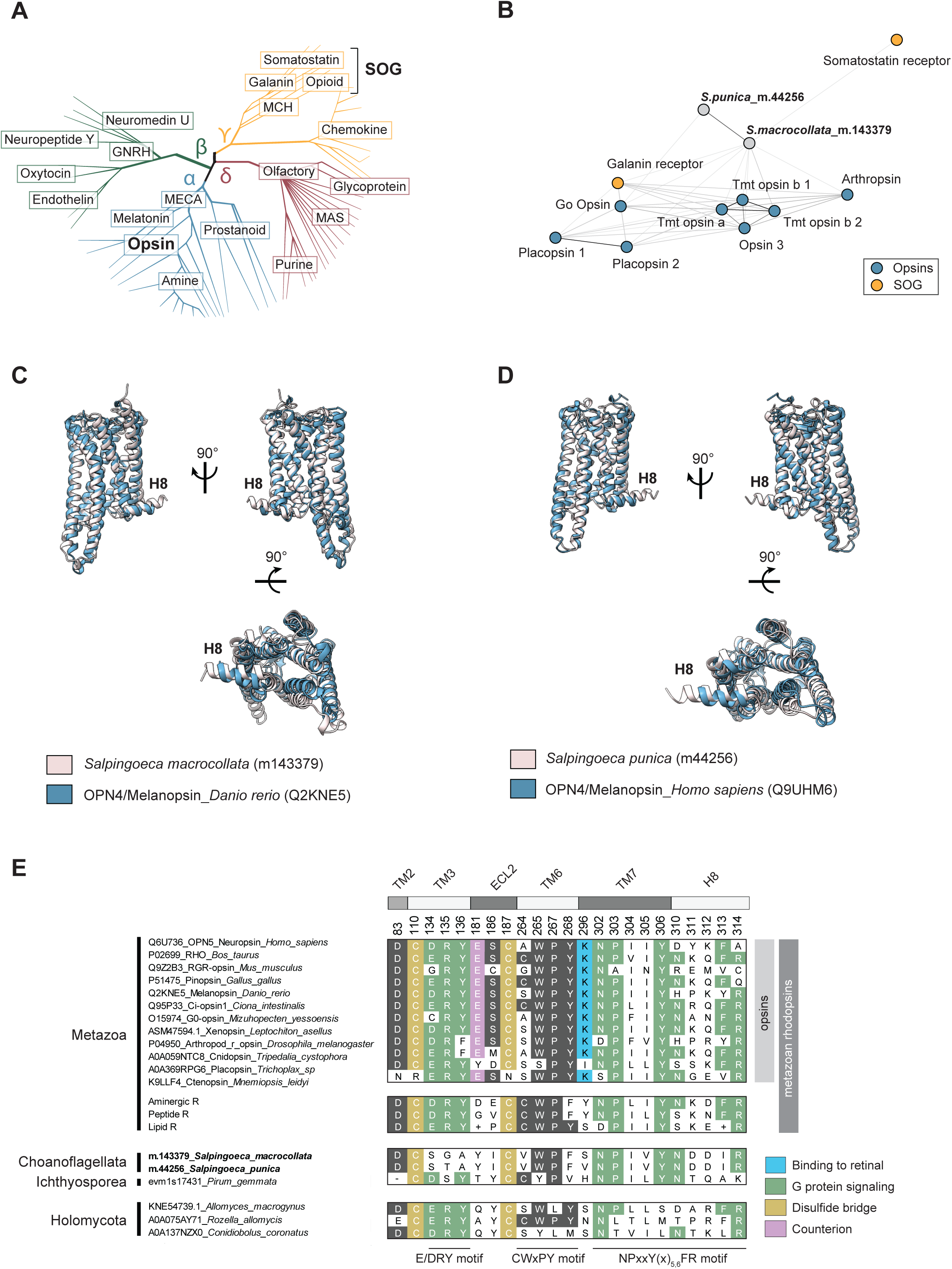
Sequence and structural similarities and differences of choanoflagellate Rhodopsins and metazoan opsins. **(A)** Rhodopsin diversity in metazoans; adapted from (Fredriksson et al. 2003; Cardoso et al. 2012; Lv et al. 2016). Rhodopsins cluster into four main groups – α (light blue), β (green), γ (yellow), and δ (maroon). The Somatostatin/Opiod/Galanin group of Rhodopsins is indicated as SOG. Opsins and SOG receptors are indicated in bold for reference. **(B)** Choanoflagellate Rhodopsins cluster with opsins and SOG receptors from metazoans. All-against-all pairwise comparison of the choanoflagellate Rhodopsins with the complete Rhodopsin repertoire of representative metazoans revealed that choanoflagellate Rhodopsins are most similar to opsins and SOG. Shown is the local sequence similarity network, with nodes indicating Rhodopsin sequences and lines indicating BLAST connections of p-value < 1e^-12^. Choanoflagellate Rhodopsins are shown in grey (*S.macrocollata*_m.143379 and *S.punica*_m.44256), metazoan opsins in blue ((Placopsin 1_*T.adherens* (A0A369S8C3), Placopsin 2_*Trichoplax* sp (XP_002114592.1), Tmt opsin a_*D.rerio* (A0A2R8Q4C0), Tmt opsin b 1_*M.mola* (ENSMMOP00000004804.1), Tmt opsin b 2 _*D.rerio* (A0A2R8Q4C0) Opsin_3 *A.platyrhynchos* (XP_012958902.3), Arthropsin 8_*D.pulex* (EFX84032.1), Go Opsin_*B.floridae* (DAC74052.1)), and metazoan SOG receptors in orange (Somatostatin receptor_*B.floridae* (XP_032829507.1), Galanin 1_*H.sapiens* (P47211)).The complete dataset of 6149 Rhodopsins used in this analysis is provided in FASTA format in Supplementary File 11. See Supplementary File 12 for the full analysis. **(C and D)** Protein structure predictions link choanoflagellate Rhodopsins to metazoan opsins. **(C)** The predicted structure of *S. macrocollata* Rhodopsin most closely matches that of opn4a/Melanopsin-A from *Danio rerio*. Shown is the predicted structural similarity of *S. macrocollata* Rhodopsin (m.143379; pink) with Foldseek top hit (E-value: 6.26e^-13^) opn4a/Melanopsin-A from *Danio rerio* (AF-Q2KNE5-F1-model_v4; blue). Low confidence regions (>70 pLDDT) were removed for clarity. Shown are views of the superimposed models from the plane of the membrane (top) and from the extracellular perspective (bottom). **(D)** The predicted structure of *S. punica* Rhodopsin most closely matches that of opn4/Melanopsin from humans. Predicted structural similarity of *S. punica* Rhodopsin (m.44256; pink) with Foldseek top hit (E-value: 8.34e^-13^) opn4/Melanopsin from humans (AF-Q9UHM6-F1-model_v4; blue). Low confidence regions (>70 pLDDT) were removed for clarity. Shown are views of the superimposed models from the plane of the membrane (top) and from the extracellular perspective (bottom). **(E)** Alignment showing the conservation of functionally important motifs in metazoan, choanoflagellate, ichthyosporean, and holomycotan Rhodopsins. The alignment includes diverse metazoan opsins, three consensus sequences of human non-opsin Rhodopsins (Aminergic, Peptide, and Lipid Rhodopsins; Supplementary File 14), the two choanoflagellate Rhodopsins identified in this study (highlighted in bold), one ichthyosporean Rhodopsin, and three representative Rhodopsins from holomycotans. Residues identified as being critical for Rhodospin protein structure and function are shown. These include: a conserved Aspartic acid (D) at position 83 in the transmembrane helix 2 (TM2); two conserved Cysteines (C; orange) at positions 110 and 187 that are involved in disulfide bond formation; and the conserved E/DRY, CWxPY, and NpxxY(x)_5,6_FR motifs (green), located in TM3, TM6, and TM7/H8, respectively (Davies et al. 2010; Nagata and Inoue 2021). These three motifs are essential for G protein interaction and to control the activity of the Rhodopsins. In addition, residues that are specific to opsins are also depicted: Glutamic acid (E)181, which acts as a counterion to the protonated Schiff base (Davies et al. 2010; Hankins et al. 2014; Nagata and Inoue 2021), Serine (S)186 in extra-cellular loop 2 (EL2), and the highly conserved Lysine (K) at position 296 (blue) in TM7, that is almost universally found across all metazoan opsins (Gühmann et al. 2022; McCulloch et al. 2023). Lys(K)296 is required for covalent binding to the 11-cis retinal chromophore (Devine et al. 2013). Notably, the two choanoflagellate Rhodopsins show a Lys296Ser (*Salpingoeca_macrocollata*_m.143379) and a Lys296Val (*Salpingoeca_punica*_m.44256) substitutions, suggesting that these Rhodopsins may not have light-responsive functions. Canonically conserved functional residues and positions follow bovine Rhodopsin numbering (Nathans and Hogness 1983). The consensus sequences of the three non-opsin subfamilies of human Rhodopsins (Aminergic R, Peptide R, and Lipid R) were downloaded from GPCRdb (https://gpcrdb.org/). (+) symbol is used when consensus sequences cannot be resolved at a given position (ambiguity). The 36 human Aminergic receptors, 76 human Peptide receptors, and 36 human Lipid receptors aligned to build these consensus sequences are provided in Supplementary File 14.

To complement our sequence-based clustering approach, we then predicted the structures of the two choanoflagellate Rhodopsins and the ichthyosporean Rhodopsin using Alphafold 3 (Abramson et al. 2024). We searched for structural matches within three Alphafold databases: AFDB-PROTEOME, AFDB-SWISSPROT, and AFDB50 (van Kempen et al. 2024). In these searches, metazoan opsins (a subclass of Rhodopsins; Fig. 4A) were recovered as the closest structural matches to choanoflagellate Rhodopsins, with e-values ranging from e^-12^ to e^-14^, with Melanopsin/OPN4, an opsin with non-visual and visual functions in metazoans (Hankins et al. 2008; Koyanagi et al. 2013; Karthikeyan et al. 2023), being the top hit for the two choanoflagellate Rhodopsins (Fig. 4C, D). In contrast, aminergic or peptide Rhodopsin receptors more closely matched the predicted structure of the ichthyosporean Rhodopsin candidate (Fig. S5). Notably, the predicted structures of choanoflagellate and ichthyosporean Rhodopsins exhibit a cytoplasmic helix eight (H8) located immediately after the end of the seventh transmembrane domain, a common feature of metazoan Rhodopsins that is involved in trafficking of the receptor to the cell membrane and in binding to G protein and β-Arrestin (Fig. 4C, D; Krishna et al. 2002; Kock et al. 2009; Knepp et al. 2012; Sensoy and Weinstein 2015; Dijkman et al. 2020).

In metazoan Rhodopsins, conserved sets of amino acids in transmembrane passes 2, 3, 6, and 7 (TM2, TM3, TM6, and TM7), along with extracellular loop 2 (ECL2), serve important structural roles and as determinants of signal transduction (Fig. 4E, Supplementary files 13 and 14; Davies et al. 2010; Hankins et al. 2014; Nagata and Inoue 2021b). We found that two cysteines (C110 and C187) whose disulfide bridges often connect ECL2 and TM3 in metazoan Rhodopsins (Rader et al. 2004), were conserved in both the choanoflagellate and ichthyosporean receptors. Similarly, asparagine D83 was conserved in TM2 of choanoflagellate and holomycotan Rhodopsins. Among the sites shown to be important for controlling G protein signaling, the CWxPY and NpxxY(x)_5,6_FR motifs were partially conserved in choanoflagellate and ichthyosporean sequences. In contrast, the E/DRY motif, which might regulate the activation of metazoan Rhodopsins (Rovati et al. 2007; Sandoval et al. 2016), was lost from TM3 in the choanoflagellate Rhodopsins but partially conserved in the ichthyosporean Rhodopsin. Finally, none of the residues mediating light sensing in opsins – namely the highly conserved Lysine K269, which mediates the binding to the chromophore retinal and the glutamic acid E181 that serves as a counterion to the protonated Schiff base (Davies et al. 2010; Hankins et al. 2014; Nagata and Inoue 2021) – were detected in CRM or holomycotan Rhodopsins, suggesting that these residues (and their functions in light sensing) evolved in stem metazoans.

#### Adhesion GPCRs

In metazoans, Adhesion GPCRs (aGPCRs) are generally the second most abundant GPCR family after the Rhodopsin family (Kamesh et al. 2008; Nordström et al. 2008; Krishnan et al. 2014; De Mendoza et al. 2014). They are critical for the multicellular biology of metazoans, in which they regulate epithelial morphogenesis, neuronal development, and immunity and are implicated in the progression of various cancers (Hamann et al. 2015a; Langenhan et al. 2016; Liebscher et al. 2022). The signature 7TM domain of aGPCRs has been detected in CRMs, holomycotans, and diverse other lineages (Fig. 2; (Krishnan et al. 2012; De Mendoza et al. 2014)).

Phylogenetic analysis of the 7TM domains of choanoflagellates uncovered at least 19 subfamilies of aGPCRs (subfamilies α-τ; Figs. S6A and S6B; Supplementary files 15 and 16), of which only one, class θ, is orthologous to a metazoan aGPCR subfamily (Bootstrap support 83%; Fig. 5A and Supplementary files 17 and 18), the ADGRV family (Weston et al. 2004; Hamann et al. 2015a; Scholz et al. 2019; Kusuluri et al. 2021). The other choanoflagellate aGPCR families presumably evolved on the choanoflagellate stem lineage or within choanoflagellates, as they are not detected in other lineages. In addition, a set of uncharacterized cnidarian and cephalochordate GPCRs grouped with a subset of filasterean aGPCRs (Bootstrap support 98%, Fig. 5A).

**Figure 5:**
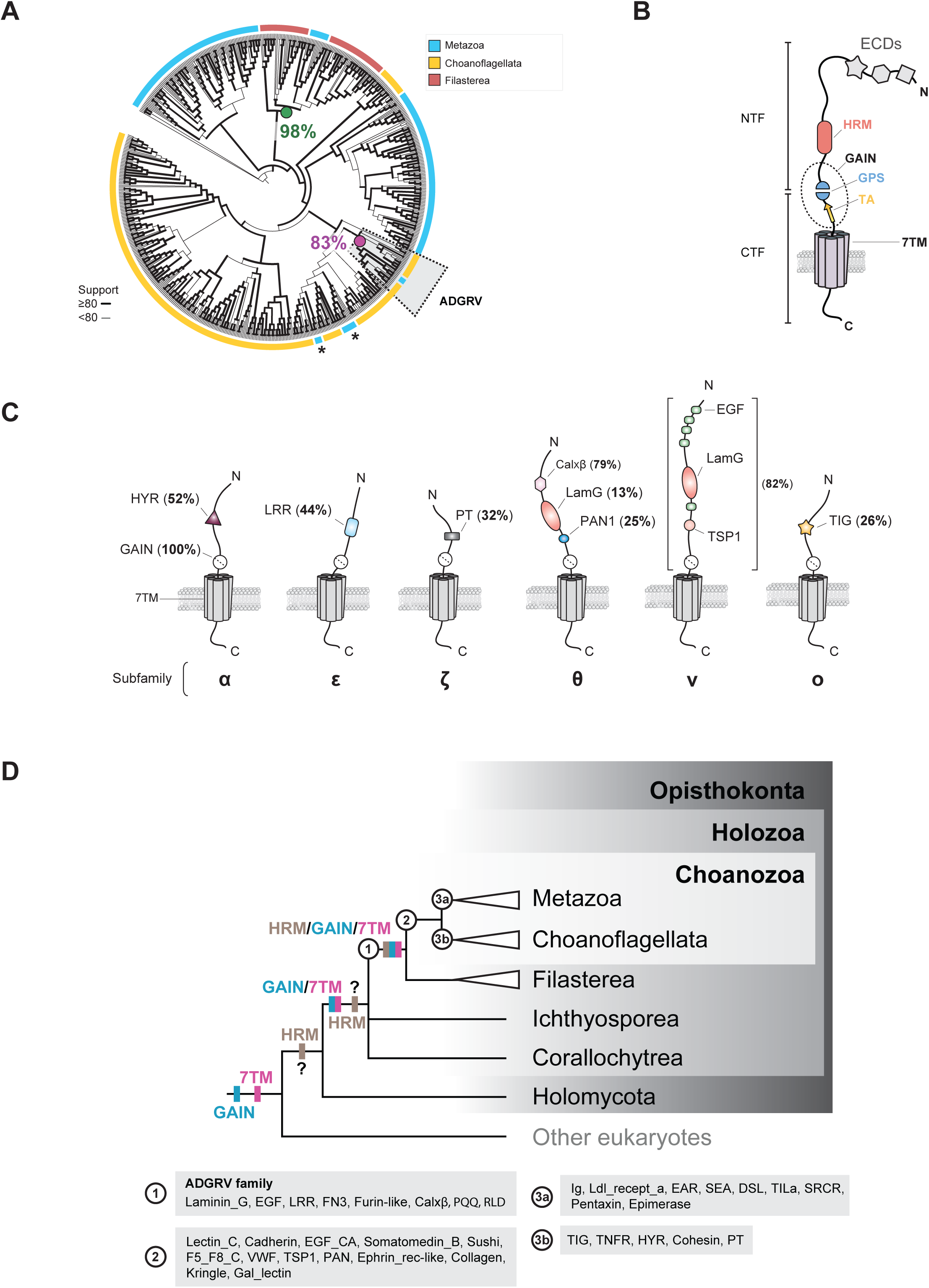
Evolution of aGPCR protein domain architecture. **(A)** Although aGPCRs were present in LECA (Fig. 2), phylogenetic analysis of holozoan aGPCRs revealed that they diversified independently in metazoans and CRMs. Most metazoan (blue), choanoflagellate (orange), and filasterean (red) aGPCRs formed distinct clades in our analysis, suggesting an absence of orthologous relationships between the 7TM region of the aGPCRs from these three clades. A notable exception was the metazoan **AD**hesion **G** protein-coupled **R**eceptor **V** (ADGRV) GPCRs that grouped with choanoflagellate 7TM sequences (dotted grey box; 83% bootstrap support for ancestral node, magenta circle), suggesting they are orthologous. In addition, we also observed that members of the metazoan ADhesion G protein-coupled Receptor A (ADGRA) subfamily (asterisks) tended to group with choanoflagellate aGPCRs but either lacked reliable confidence value support or were not systematically recovered in all the inferred phylogenies. A subset of filasterean aGPCRs clustered with a set of uncharacterized cnidarian and cephalochordate receptors (bootstrap support 98%, green circle). This maximum-likelihood phylogenetic tree infers the evolutionary history of the 7TM domain of 329 choanoflagellate aGPCRs identified in this study, along with 76 filasterean aGPCRs and 253 metazoan aGPCRs. The 7TM sequence of a ciliate aGPCR (Stentor_coeruleus_OMJ80129.1_19670) was also included in the analysis and used as an outgroup to root the tree. The width of branches scales with UFboot support for the ancestral node. Branch lengths do not scale with evolutionary distance in this rendering. All the 7TM sequences used in this phylogenetic reconstruction are found in Supplementary File 17, and the fully annotated version of this phylogenetic tree, including bootstrap values, branch lengths, and all species names, is found in Supplementary File 18. **(B)** Protein domain architecture of metazoan aGPCRs. Like all GPCRs, aGPCRs contain a 7TM domain (represented by seven barrels) that anchors the protein in a lipid bilayer. What separates aGPCRs from other GPCRs is their possession of an autoproteolytic GAIN domain (dotted oval) containing a proteolysis site (GPS, blue) and a hydrophobic tethered agonist element (TA, orange). The cleavage site in the GPS represents the boundary between the N-terminal fragment (NTF) and the C-terminal fragment (CTF). Many, but not all, aGPCRs also contain an HRM domain (salmon) within a few hundred amino acids of the GPS (**Fig. S10C**) (Prömel et al. 2012b; Araç D, Sträter N 2016)In metazoans, the NTF often contains a diversity of additional extracellular protein domains (ECDs, represented by grey star, hexagon, and square) that likely contribute to the diversity of ligands bound by aGPCRs (Araç and Leon 2019; Knierim et al. 2019). (**C**) Diversity and conservation of protein domains in the NTFs of different choanoflagellate aGPCR families. Like metazoan aGPCRs, nearly all choanoflagellate aGPCRs contain a GAIN domain with conserved GPS and TA (**Fig. S9**). Many aGPCR subfamilies identified through their 7TM were also distinguishable through their characteristic combinations of N-terminal protein domains (**Fig. S7A**). Shown here are six subfamilies – α, ε, ζ, θ, ν, and ο – whose members frequently share one or more protein domains. While subfamilies α, ε, θ, ν, and ο are seemingly unique to choanoflagellates, subfamily θ is notable because the full-length proteins (both the NTFs and CTFs) are homologous to those of the ADGRV subfamily in metazoans (Fig. 5A and **Fig. S8**). Percentages indicate the number of members within a given subfamily possessing a conserved protein domain (subfamilies α, ε, ζ, θ, and ξ) or a conserved combination of protein domains (subfamily ν). All the choanoflagellate aGPCR NTF sequences used in these analyses are found in Supplementary File 19. **(D)** Hypothesized evolution of the HRM/GAIN/7TM module and additional ECDs in aGPCRs. For four nodes on the eukaryotic tree of life – (1) the last common ancestor of filastereans and choanozoans, (2) stem choanozoans, (3a) stem metazoans, and (3b) stem choanoflagellates – we reconstructed the phylogenetic distribution of diverse protein domains in the NTFs of aGPCRs. Most aGPCRs in metazoans, choanoflagellates, and filastereans contain a protein domain module composed of a GAIN domain and a 7TM. Additionally, the HRM/GAIN/7TM module, which is less common than the GAIN/7TM module in aGPCRs, is nonetheless conserved across most metazoan, choanoflagellate, and filasterean lineages. The linkage of the GAIN and 7TM domains likely occurred in stem holozoans, as today the two domains are restricted to the aGPCRs of extant holozoans, although the two domains are encoded in other genes in diverse non-holozoans (Araç et al. 2012; Krishnan et al. 2012; De Mendoza et al. 2014), suggesting that they originated in earlier branches of the eukaryotic tree. The linkage of the HRM domain with the GAIN/7TM module likely occurred in the last common ancestor of filastereans and choanozoans. The HRM domain is largely restricted to holozoans, although we found six non-aGPCR proteins containing HRM domains in the proteome of the nucleariid *Fonticula alba* (Fig. S10D). Therefore, HRMs either evolved in stem holozoans and were incorporated into the nucleariids by horizontal gene transfer, they are homologous in the two lineages and were lost from most holomycotans, or the similarities of HRMs in *F. alba* and holozoans are the result of convergent evolution. The repertoires of ECDs inferred for each node – (1), (2), (3a), and (3b) – are shown in grey boxes labeled accordingly. Finally, ADGVR likely evolved in the last common ancestor of filastereans and choanozoans.

Unlike other GPCRs, metazoan aGPCRs have a large N-terminal extracellular region with diverse protein domains that are linked to the 7TM domain by a conserved GPCR-autoproteolysis-inducing (GAIN) domain (Fig. 5B; (Araç et al. 2012; Langenhan et al. 2013; Hamann et al. 2015a)). Ligand binding can cause the N-terminal fragment (NTF) to be released from the rest of the protein at the GPCR proteolytic site (GPS), exposing a tethered agonist element (TA) that activates downstream G protein signaling (Barros-Álvarez et al. 2022; Kleinau et al. 2022; Ping et al. 2022; Xiao et al. 2022; Seufert et al. 2023). The structural complexity of the metazoan NTF, including the GAIN domain and additional extracellular domains (ECDs), appears restricted to holozoans, as holomycotans and other eukaryotes have shorter, unstructured N-termini (Krishnan et al. 2012). Therefore, a detailed analysis of aGPCR subfamilies and their protein domains in CRMs promised to illuminate the evolution of aGPCRs after the holozoan-holomycotan divergence.

Although the phylogeny of aGPCRs in choanoflagellates was inferred solely based on the comparison of their 7TM domains, the resulting groups tended to cluster receptors that also shared sequence similarity and protein domain architecture in their NTFs (Fig. 5C and Fig. S7A). The connection between the 7TM phylogeny and the N-terminal protein domain architectures of different aGPCRs, which was previously reported for metazoan aGPCR families (Bjarnadóttir et al. 2007; Hamann et al. 2015a), is exemplified by the choanoflagellate aGPCR groups α, ε, ζ, θ, ν, and σ, whose NTFs generally contain one or multiple conserved protein domains that are shared among members (Fig. 5C). Comparison of metazoan and CRM NTFs revealed that only ADGRV receptors share robust N-terminal sequence similarity among metazoan, choanoflagellate, and filasterean aGPCRs (Fig. S8 and Supplementary file 19).

While the GAIN and aGPCR 7TM domains evolved before the origin of opisthokonts (Araç et al. 2012; Krishnan et al. 2012; De Mendoza et al. 2014), we detected the fusion of these two domains into a single module (GAIN/7TM) in most, but not all, holozoan aGPCRs (Fig. 5D, Fig. S7B and S9A; Supplementary file 20; Prömel et al, 2013; Krishnan et al. 2014). Therefore, the GAIN/7TM module likely evolved in stem holozoans. Indeed, motifs necessary for self-proteolysis in metazoan GAIN domains (Araç et al. 2012; Stoveken et al. 2015; Kleinau et al. 2022; Ping et al. 2022; Xiao et al. 2022; Seufert et al. 2023) are also conserved in CRM aGPCRs, suggesting that these receptors may similarly undergo cleavage and activation (Fig. S9B and S9C; Supplementary file 21).

The presence of other aGPCR ECDs (Fig. 5B) likely evolved in the last common ancestor of filastereans and choanozoans. This is supported by the absence of ECDs from ichthyosporean, corallochytrean, or non-holozoan aGPCRs (Fig. S7B), coupled with the conservation of Laminin_G, EGF, LRR, FN3, Furin-like, Calxβ, PQQ, and RLD domains in the aGPCRs of filastereans with those of either choanoflagellates or metazoans (Fig. 5D, node “1”). We infer that the aGPCR ECD repertoire diversified in the choanozoan stem lineage (Fig. 5D, node “2), followed by lineage-specific acquisitions of additional ECDs in metazoans and choanoflagellates (Fig. 5D, node “3a” and “3b”; Fig. S7B). Interestingly, the HRM domain — previously identified only in the metazoan Secretin and aGPCR families (Fig. S10A and Schiöth and Lagerström 2008; Nordström et al., 2009b; Krishnan et al., 2012; Krishnan et al., 2014; Araç D, Sträter N 2016) — was also found in 22 choanoflagellate and 6 filasterean aGPCRs in our analysis (Fig. S10B). Like its positioning in metazoan aGPCRs (Araç et al. 2012; Prömel et al. 2012a), the HRM domain is located directly N-terminal to the GAIN domain in choanoflagellate and filasterean aGPCRs, with a comparable separation of ∼280-330 amino acids between the C-terminus of the HRM domain and the N-terminus of the 7TM domain (Fig. S10B and S10C).

These findings suggest that the HRM domain and its integration into an HRM-GAIN-7TM module likely originated in the last common ancestor of Filasterea and Choanozoa (Fig. 5D). Supporting this view, the HRM domain was absent from all non-holozoan proteomes analyzed, with one exception: the nucleariid *Fonticula alba* (Fig. S10D), a close relative of Fungi (Liu et al. 2009; Torruella et al. 2015). *Fonticula alba* encodes five non-GPCR transmembrane proteins that contain HRM domains (Supplementary file 22).

These findings clarify the evolution of key features of holozoan aGPCRs. While the autoproteolytic GAIN domain (Araç et al. 2012), the aGPCR 7TM domain (Krishnan et al. 2012; De Mendoza et al. 2014), and possibly the HRM domain (this study) existed before the emergence of Holozoa, our analysis indicates that they were first combined into a single module in holozoans. The GAIN-7TM module originated in stem holozoans, followed by the positioning of the HRM domain N-terminal to the GAIN domain in the last common ancestor of Filasterea and Choanozoa. This was followed by the acquisition of additional extracellular protein domains in choanozoans and filastereans (Fig. 5D).

#### cAMP Receptors

Twenty-eight choanoflagellate GPCRs grouped with cAMP Receptors from diverse other eukaryotes (Fig. 1B). These receptors possess short N- and C-termini and a unique 7TM signature that distinguishes them from other GPCRs (Saxe et al. 1993; Krishnan et al. 2012; Greenhalgh et al. 2020a).

The cAMP Receptors were present in stem eukaryotes and are inferred to have given rise to GRAFS GPCRs (Nordström et al. 2011; Krishnan et al. 2012). cAMP Receptors were first described in the amoebozoan *Dictyostelium discoideum*, where they control chemotaxis, aggregation, morphogenetic movements, cell growth, and many developmentally regulated genes (Klein et al. 1988; Johnson et al. 1993; Saxe et al. 1993; Louis et al. 1994). While cAMP Receptors bind to cAMP in *D. discoideum* (Juliani and Klein 1981; Caterina et al. 1995; Milne et al. 1997), the precise cAMP binding site(s) have yet to be experimentally identified (Greenhalgh et al. 2020b). Since then, cAMP Receptors have been identified in holozoans, holomycotans, alveolates, plants, and other lineages (Fig. 2; (Krishnan et al. 2012; De Mendoza et al. 2014)). Notably, although present in diverse metazoans, cAMP Receptors have been lost from vertebrates. Previous studies found that the choanoflagellates *S. rosetta* and *M. brevicollis* each encode a single cAMP Receptor (Krishnan et al. 2012; De Mendoza et al. 2014). This study found cAMP Receptors in 17 additional choanoflagellate species, including craspedids and acanthoecids (Fig. 2).

#### GPR137

GPR137 is a lysosomal receptor that controls mTORC1 translocation to the lysosomes and modulates the Hippo pathway (Gan et al. 2019). Moreover, GPR137 regulates autophagy, cell growth, and cell proliferation and is implicated in various cancers (Men et al. 2018; Iwasa et al. 2023; Li et al. 2024). GPR137 does not possess other protein domains outside the 7TM domain. The receptor is broadly conserved in metazoans, and homologs have been found in the choanoflagellate *Monosiga brevicollis* and the amoebozoan *D. discoideum* (Fig. 2 and (Gan et al. 2019)).

We detected GPR137 in a wide range of choanoflagellate species, as well as in filastereans, ichthyosporeans, and non-holozoans, including nucleariids and other eukaryotic lineages (Fig. 2). Thus, GPR137 probably evolved at the stem of the eukaryotic tree. Whether GPR137 controls cell growth and the cell cycle in non-metazoans remains to be tested.

#### GPR157

GPR157 Receptors promote neuronal differentiation of radial glial progenitors in mice, where they localize to the primary cilia and signal via a G(q)-protein that activates a phosphatidylinositol-calcium second messenger (Takeo et al. 2016). No additional domains outside the 7TM have been detected in GPR157.

GPR157 had previously been identified only in mammals (Takeo et al. 2016). In this study, we detected GPR157 homologs in most metazoans, including sponges, as well as in choanoflagellates, filastereans, nucleariids, amoebozoans, rhizarians, and various other eukaryotes, suggesting that GPR157 likely evolved at the root of the eukaryotic tree (Figs. 2 and S11A; Supplementary File 23). Choanoflagellate and metazoan GPR157 share a level of sequence similarity with the 7TM domains of cAMP, aGPCRs, and GPCR PIPKs (Fig. 1B). In addition, we noticed a cytoplasmic helix 8 (H8), better known from Rhodopsins, directly C-terminal to TM7 in choanoflagellate, metazoan, and other eukaryote GPR157 protein structure predictions (Fig. S11B). Striking conservation of motifs that have not previously been characterized was noted in TM2 (“LLxxLSL/V/IxD”), ECL2 (“WCWI/V/L”), and TM7 (“QGxxNxIxF”) of the GPR157 TM domain (Fig. S11C).

#### GPR107/108, TMEM145, GPR180, and TMEM87 (GOST GPCRs)

We identified 63 choanoflagellate GPCRs that clustered with a group of metazoan GPCRs named “GOST” for Golgi-dynamics domain seven-transmembrane helix proteins (Fig. 1 and Table 1; (Hoel et al., 2022). All GOST members exhibit a Golgi dynamics (GOLD) domain in their extracellular region, which is possibly involved in ligand recognition, followed directly by the 7TM domain (Anantharaman and Aravind 2002; Hoel et al. 2022). GOST GPCRs include GPR107/108, TMEM145, GPR180, and TMEM87 – all found in choanoflagellates – along with GPR181 and Wntless, neither of which were detected in choanoflagellates.

The structural similarity between choanoflagellate and metazoan GOST receptors includes both the 7TM domain and the N-terminal GOLD domain (Fig. S12). Nonetheless, it is unclear whether the GOST GPCRs are monophyletic, as GPR107/108/TMEM87 and GPR180/TMEM145 grouped separately in our analyses of the 7TM domains (Fig. 1B). Despite this uncertainty, all GOST members have homologs in holozoans and various other eukaryotes (Fig. 2), suggesting that they were present in stem eukaryotes.

GOST GPCRs are understudied, but the characterization of several members showed that they localize at the Golgi and endocytic vesicles, suggesting a common function in trafficking membrane-associated cargo (Zhou et al. 2014; Hirata et al. 2015; Shin et al. 2020; Hoel et al. 2022; Mitrovic et al. 2024). In addition, a recent study found that TMEM145 also contributes to the structural integrity of hair cell stereocilia (Roh et al. 2025), expanding the functions associated with GOST GPCRs. Although ligands have been proposed for some GOST proteins ̶ neurostatin for GPR107 (Yosten et al. 2012; Yang et al. 2024), gambogic acid for GPR108 (Lyu et al. 2022), L-lactate (Mosienko et al. 2018), and CTHRC1 for GPR180 (Balazova et al. 2021) ̶ it is presently unclear whether these receptors are competent for G protein signaling since no molecular evidence has been reported for any member of this family (Balazova et al. 2021; Hoel et al. 2022).

#### Hi-GOLD

Seventeen choanoflagellate GPCRs clustered independently from other known GOST GPCRs but exhibit a similar topology with a conserved GOLD domain directly N-terminal to their 7TM (Fig. 1B and Fig. S12). We named this group of GPCRs “Hidden GOLD” (Hi-GOLD). Hi-GOLD GPCRs are conserved in both craspedids and acanthoecids (Fig. 2). Additionally, they have homologs in metazoans, including sponges, ctenophores, placozoans, cnidarians, holothuroideans, and cephalochordates, but are absent from vertebrates (Supplementary File 24). Hi-GOLD GPCRs were also detected in other CRMs (filastereans and ichthyosporeans) and non-holozoans, such as amoebozoans. Thus, we have uncovered a new group of GPCRs bearing structural similarity with GOST GPCRs and having ancient origins in eukaryotes.

#### GPCR PIPK

Twenty-seven choanoflagellate GPCRs were identified as GPCR PIPKs based on their domain composition. These receptors comprise an N-terminal 7TM combined with a C-terminal phosphatidylinositol phosphate kinase (PIPK) catalytic domain (Bakthavatsalam et al. 2006; Van Den Hoogen et al. 2018; van den Hoogen and Govers 2018) (Fig. S13A and S13B). A diagnostic “LR(x)_9_GI” motif, previously found to be present in the linker region between the 7TM domain and the PIPK domain of these receptors (Van Den Hoogen et al. 2018) was also found in the choanoflagellate GPCR PIPKs (Fig. S13C).

Initially discovered in oomycetes, where the family greatly expanded, GPCR PIPKs have now been detected in diverse eukaryotes and sponges (Bakthavatsalam et al. 2006; Van Den Hoogen et al. 2018). Sponges appear to be the only metazoans encoding GPCR PIPKs (Fig. 2 and (Van Den Hoogen et al. 2018)). Thus, genes encoding GPCR PIPKs were likely present in the last common eukaryotic ancestor and secondarily lost in diverse eukaryotes, including a loss after the divergence of sponges from the rest of metazoans.

A previous study found that one GPCR PIPK each was encoded in *S. rosetta* and *M. brevicollis* species (Van Den Hoogen et al. 2018); our analysis revealed that GPCR PIPKs are also expressed by most craspedid choanoflagellates (Fig. 2). However, no acanthoecid choanoflagellates in our study appeared to possess these receptors (Fig. 2).

While GPCR PIPKs have previously been grouped into the same family because of their domain architecture ̶ the 7TM domain combined with a PIPK domain ̶ our analysis suggests that this family is also supported by the sequence homology of their 7TM domain. Indeed, both choanoflagellate and sponge GPCR PIPKs analyzed in this study clustered together, forming a class distinct from other GPCR families (Fig. 1B). In addition, blasting the 7TM domains of choanoflagellate GPCR PIPKs against the full EUKPROT database (Richter et al. 2022) recovered GPCR PIPKS from various other eukaryotes (Filasterea, Discoba, Collodictyonida, Telonemida, Rhodelphida, Ancyromonadida, Centroplasthelida) suggesting that the 7TM domain is enough to find homologs, independently of the C-terminal PIPK domain. Thus, GPCR PIPKs likely constitute a family of GPCRs on their own, with a signature 7TM domain that might have evolved in stem eukaryotes.

Experimental data regarding the function of GPCR PIPKs is limited. Pioneer studies in oomycetes implicated the receptor in the regulation of sexual reproduction and virulence (Hua et al. 2013). In contrast, studies of *D. discoideum* GPCR PIPKs suggested a role in phagocytosis, cell density sensing, and bacterial defense (Bakthavatsalam et al. 2007; Riyahi et al. 2011).

Although the catalytic domain of GPCR PIPKs is predicted to have a role in phospholipid signaling, experimental evidence for this hypothesis is still missing. Moreover, it is not known whether G protein signaling is involved downstream of GPCR-PIPKs (van den Hoogen and Govers 2018). Similarly, no ligands for GPCR PIKs have been identified.

#### GPRch3

The GPRch3 cluster contained 28 choanoflagellate GPCRs with short and unstructured N-terminal and C-terminal regions (Fig. S1A); they did not show any similarity to any other previously described GPCR families, either by sequence or structure (Fig. 1B). Nonetheless, these GPCRs appear to have homologs in diverse metazoans (sponges, ctenophores, cnidarians, protostomes and deuterostomes), corallochytreans, and non-holozoans (e.g. amoebozoans and alveolates; Fig. 2). Thus, the GPRch3 GPCRs likely represent a new family of GPCRs with ancient origin in eukaryotes that is retained in metazoans.

### Holozoan GPCRs

#### GPR155

GPR155, also known as lysosomal cholesterol sensing (LYCHOS) protein, is an atypical GPCR that binds to lysosomal cholesterol and regulates mTORC1 activity (Shin et al. 2022; Bayly-Jones et al. 2024; Schöneberg 2024). It contains an unusual 17-transmembrane helix structure composed of an N-terminal transporter-like domain (10 transmembrane helices) fused to a 7TM, with Dishevelled, EGL-10 and Pleckstrin (DEP) domains located at the C terminus. The ability of GPR155 to transduce signals via G proteins is presently unclear. While it has been suggested that the 7TM of GPR155 shares similarities with the aGPCR family (Bayly-Jones et al. 2024), it appears to form a cluster of its own in our analysis (Fig. 1B).

GPR155 has been detected in diverse bilaterians (Vassilatis et al. 2003; Umeda et al. 2017; Wang et al. 2018), and its transporter domain has an ancient origin in eukaryotes (Dabravolski and Isayenkov 2022; Bayly-Jones et al., 2024). The finding that GPR155 homologs are encoded in CRMs, but are absent from non-holozoan lineages, suggests that GPR155 originated in the ancestor of holozoans (Fig. 2). Moreover, the conservation of GPR155 in CRMs extends beyond the 7TM as both the N-terminal transporter-like domain and the C-terminal DEP domain are detected in GPR155 homologs from CRMs (Fig. S15).

While choanoflagellates and close relatives can produce a variety of sterols (Kodner et al. 2008; Gold et al. 2016; Najle et al. 2016), they are not known to synthesize cholesterol, a vital lipid for metazoans (Zhang et al. 2019; Shamsuzzama et al., 2020). The ligand(s) of GPR155 in CRMs and whether it mediates sterol-induced signaling await further investigation.

#### GPR143

GPR143 homologs were detected in diverse metazoans, including sponges (Krishnan et al. 2014), and the filasterean *Capsaspora owczarzaki* (Fig. 2 and (De Mendoza et al. 2014)), but not in choanoflagellates. Therefore, GPR143 likely originated in holozoans and was lost in stem choanoflagellates (Fig. 2).

While little is known about GPR143, the receptor is expressed in pigment-producing cells in the skin and eyes of metazoans, where it controls melanosome biogenesis, organization, and transport (Bueschbell et al. 2022). The downstream signaling pathway is mostly uncharacterized, but GPR143 associates with several Gα and Gβ subunits, along with β-Arrestin (Schiaffino et al. 1999; Schiaffino and Tacchetti 2005; De Filippo et al. 2017). L-3,4-dihydroxyphenylalanine (L-DOPA) and dopamine have been suggested as ligands for GPR143 (Lopez et al. 2008; Goshima et al. 2019).

### Stem eukaryote GPCRs that were lost from metazoans

#### Rémi-sans-famille (RSF)

28 choanoflagellate receptors clustered together and did not appear to share sequence similarities with other GPCR families previously described in eukaryotes. These GPCR candidates contained a 7TM domain but no additional domains (Fig. S16A). Their predicted structures revealed an additional short helix between TM6 and TM7 with a “NxLQxxMNxL” conserved motif (Fig. S16A and S16B). Structural similarity search suggested a weakly supported connection to the THH1/TOM1/TOM3 GPCR family (Yamanaka et al. 2000; Lu et al. 2018) (Fig. S16C).

While these GPCRs are absent from metazoans, we detected homologs in filastereans, as well as in other non-holozoans, including provorans (Tikhonenkov et al. 2022), rhodophytes, alveolates, amoebozoans, and ancyromonads (Fig. 2). Thus, it is likely that this family of GPCRs evolved early during the evolution of eukaryotes and was secondarily lost in various lineages, including metazoans. We named this new class of GPCRs “Rémi-sans-famille” (RSF), inspired by a fictional orphan (Malot 1878).

#### GPRch1

Clustering on the 7TM domains revealed a group of 16 craspedid choanoflagellate GPCRs, which we have named GPRch1 GPCRs. Although these proteins lack a PIPK domain, their 7TM domains resembled those of GPCR PIPKs from choanoflagellates and sponges (Figs. 1B and S14B). The GPRch1 7TMs also clustered with 7TMs from amoebozoans, stramenopiles, alveolate, and apusomonadids; interestingly, the non-choanoflagellate GPCRs in this cluster all contained a PIPK domain in the C-terminus. Therefore, we infer that there was a duplication and divergence of GPCR PIPKs in stem eukaryotes, with one of the paralogs giving rise to GPRch1 GPCRs. Subsequently, the choanoflagellate GPRch1 GPCRs lost the PIPK domain. The absence of GPRch1 GPCRs from acanthoecid choanoflagellates, other CRMs, and metazoans suggests that they were lost from these lineages (Fig. 2).

### Choanoflagellate-specific GPCRs

#### GPCR-TKL/K and GPCR-TKs

The GPCR-TKL/K family was identified by examination of a cluster of 176 acanthoecid choanoflagellate GPCRs, 92 of which contain a kinase domain in their C-terminus (Fig. 1 and Fig. 2; Fig. S17A, S17B, and S17C). Further analysis showed that the kinase domains of these GPCRs relate to Tyrosine Kinase-like (TKL) domains and other non-tyrosine kinase domains (Fig. S17B and Supplementary files 25, 26, and 27); we therefore named these receptors GPCR-Tyrosine Kinase-Like/Kinase (GPCR-TKL/K).

The fusion between a 7TM and a kinase domain has recently been reported in oomycetes and amoebozoans but was not found in other eukaryotes (Judelson and Ah-Fong 2010; Van Den Hoogen et al. 2018). Thus, the acanthoecid GPCR-TKL/K represents the first observation, to our knowledge, of a GPCR-kinase fusion in holozoans. While the overall protein domain architecture of GPCR-TK/Ks is conserved between choanoflagellates, oomycetes, and amoebozoans, the 7TM domains of acanthoecid GPCR-TKL/K receptors did not cluster with any other GPCRs in eukaryotes (Fig. 2), suggesting that this family probably originated through convergent evolution in stem acanthoecids (Fig. 2 and Fig. S17D).

Additionally, we detected 11 choanoflagellate GPCRs that contain a predicted tyrosine kinase domain in their C-terminal region, but that did not cluster with the GPCR-TKL/Ks mentioned above (Fig. S17B and Fig. S17C; Supplementary files 25, 26, 27, 28, and 29). These GPCRs are found in both acanthoecids and craspedids but failed to form a unified family based on their 7TMs (Fig. S17C and S17D). Unlike their 7TM domains, the kinase domains of these GPCRs clustered together and separately from the GPCR TKL/K kinase domains; all recovered strong blast hits among curated metazoan tyrosine kinases (Fig. S17C and Supplementary file 26). Therefore, we identified these GPCRs as GPCR Tyrosine Kinases (GPCR-TKs).

While no metazoan homologs were found when using the 7TM domain of choanoflagellate GPCR-TKs as queries, using the conserved tyrosine kinase domains as queries recovered GPCR-TKs in sponges but not in other metazoan lineages or other holozoans (Fig. S17E). To test whether GPCR-TKs in sponges and choanoflagellates are homologous, we performed phylogenetic analyses of their TK and 7TM domains (Fig. S17F and G; Supplementary files 30, 31, 32, and 33). While the TK domains of GPCR-TKs from sponges and choanoflagellates formed a well-supported clade, their 7TM domains did not. These results point to a heterogeneous evolutionary history that may include domain swapping (i.e. ancestral GPCR-TKs in which the 7TM domain was replaced in either the sponge or choanoflagellate lineages) or convergent evolution, in which homologous 7TM domains fused with unrelated 7TM domains in the sponge and choanoflagellate lineages.

Together, our data suggest that the fusion of a C-terminal kinase domain to a 7TM domain likely evolved repeatedly and independently among eukaryotes and within the choanoflagellate phylogeny. The evolution of GPCR-TKs seems to be restricted to choanoflagellates and early-branching metazoans. The evolutionary and molecular implications underlying the fusion of a 7TM domain with an intracellular kinase domain are unknown, and, to our knowledge, no GPCR-TK, GPCR-TKL, or GPCR-Ks have been functionally characterized.

### Additional GPCRs

#### GPRch2

A cluster of twelve choanoflagellate GPCRs showed no similarity to previously identified GPCR families (Fig. 1). Although these GPCRs do not possess additional protein domains outside their signature 7TM domain (Fig. S18A), the GPRch2 GPCRs exhibit an unusual large intracellular loop 3 (ICL3) linking the TM5 to the TM6 (Fig. S18A and S18B).

While homologs of these GPCRs were also found in filastereans, ichthyosporeans, and corallochytreans, they are absent from metazoans and were only detected in green algae outside of holozoans (Fig. 2). Thus, it is likely that the GPRch2s evolved in stem holozoans either *de novo*, or via a horizontal transfer from green algae, and were subsequently lost in stem metazoans.

#### Frizzled

The Frizzled family, which encompasses Frizzled GPCRs and Smoothened, controls cell proliferation, cell fate, tissue polarity, and cell polarity during metazoan development (Taipale and Beachy 2001; Logan and Nusse 2004; Schulte 2024). These receptors have a highly conserved cysteine-rich domain called the Frizzled (Fz) domain, located in their extracellular N-terminus (Schiöth and Lagerström 2008a; Zheng and Sheng 2024). This domain is connected to the seven-transmembrane (7TM) region by a linker segment. The Fz domain serves as the ligand-binding site for most Frizzled receptors. Several ligands have been found to activate members of the Frizzled family, of which the secreted glycoprotein Wingless/Int-1 (Wnt) has been the most studied (Schulte 2024).

Although initially characterized in *Drosophila*, Frizzled receptors have been detected in most metazoans, including sponges (Fig. 2; (Krishnan et al. 2014; Holzem et al. 2024)). While Frizzled receptors have not been previously detected in CRMs, homologs were found in fungi and amoebozoans, suggesting a probable origin of the family in the last common ancestor of opisthokonts and amoebozoans (Krishnan et al. 2012; De Mendoza et al. 2014). Notably, fungal and amoebozoan Frizzled homologs also possess a Fz domain in their N-termini, supporting an overall conserved receptor topology.

While we could not detect members of the Frizzled family in any of the 23 choanoflagellate species used in our study (Fig. 2), we found four Frizzled/Smoothened receptors in the transcriptome of the corallochytrean *Syssomonas multiformis* (Fig. 2; Fig. S19A and S19B; Supplementary File 32). However, contaminating sequences have been reported in this data set, and therefore, this observation should be treated with caution (Hehenberger et al. 2017).

## DISCUSSION

Our transcriptomic and genomic survey demonstrates that choanoflagellate GPCRomes are richer than previously understood, both in the number and diversity of GPCRs represented. We identified 18 distinct GPCR families in choanoflagellates, of which 12 ̶ Rhodopsin, GPR155, GPR157, GPCR TKL/K, TMEM145, TMEM87, GPR180, Hi-GOLD, RSF, GPRch1, GPRch2, and GPRch3 ̶ were newly described in these organisms (Table 1, Fig. 2). Among these families, we observed that aGPCRs generally constitute the largest class of GPCRs in choanoflagellates, apart from the GPCR TKL/K family in acanthoecids. Most choanoflagellate GPCR families are conserved in metazoans or other eukaryotes. This supports the view that GPCRs are ancient gene families found across eukaryotic diversity (Krishnan et al. 2012; De Mendoza et al. 2014; Mojib and Kubanek 2020).

In addition, by assessing the conservation of choanoflagellate GPCRs in other eukaryotes, we uncovered GPCR families in metazoans that, to our knowledge, had not been reported in this clade before. These include Hi-GOLD, GPRch3, and GPCR TKL/Ks. In addition, we found that five GPCR families ̶ RSF, Hi-GOLD, GPRch3, GPR157, and GPRch1 ̶ are conserved across diverse eukaryotes, thereby expanding the repertoire of GPCRs inferred at the root of the eukaryotic tree (De Mendoza et al. 2014). Future increases in the sequencing of non-metazoans will likely expand further the inferred diversity of GPCRs present in the progenitors of metazoans and other eukaryotes.

The identification of Glutamate Receptors, Rhodopsins, and aGPCRs in choanoflagellates, three GPCR families with relatively well-characterized subfamilies in metazoans (Fredriksson et al. 2003; Pin et al. 2003; Bjarnadóttir et al. 2004; Chun et al. 2012; Scholz et al. 2019; Ellaithy et al. 2020b; Wittlake et al. 2021), led us to investigate whether these subfamilies are conserved in choanoflagellates and close relatives. Based on analyses of the 7TM domains, we found relatives of the main subfamilies of metazoan Glutamate Receptors in CRMs (Fig. 3B). In contrast, most aGPCR subfamilies reported in metazoans have no obvious orthologs in CRMs, suggesting that metazoan and CRM aGPCRs diversified independently (Fig. 5). One exception was the ADGRV family (Weston et al. 2004; Hamann et al. 2015a; Kusuluri et al. 2021), for which clear orthologs are found in choanoflagellates and filastereans (this study and (Krishnan et al. 2012; Peña et al. 2016)).

Whereas Rhodopsins are the most abundant GPCRs in metazoans, we identified only three metazoan-like Rhodopsins in CRMs: two in choanoflagellates and one in an ichthyosporean. In addition, we uncovered Rhodopsin-like GPCRs from diverse fungi and amoebozoans (Supplementary File 7), corroborating the findings of prior studies (Krishnan et al. 2012; De Mendoza et al. 2014). Sequence comparisons and structural predictions suggest that the two choanoflagellate Rhodopsins most closely resemble metazoan opsins, a Rhodopsin subfamily (Fredriksson et al. 2003; Mickael et al. 2016). Opsins, light-detecting receptors supporting vision, have only been described in metazoans (Feuda et al. 2012; Fleming et al. 2020; Wong et al. 2022; Hagen et al. 2023; Aleotti et al. 2025). Unlike metazoan opsins, the two choanoflagellate Rhodopsins lack canonical residues essential for light-sensing, including the highly conserved Lysine K296, which mediates binding to the chromophore retinal (Davies et al. 2010; Hankins et al. 2014; Nagata and Inoue 2021). Thus, while it is formally possible that Rhodopsins existed in stem choanoflagellates and were lost in most modern choanoflagellate lineages, either horizontal gene transfer or convergent evolution in the shared ancestor of *S. macrocollata* and *S. punica* are similarly plausible explanations for their presence in these species. Differentiating between these alternative evolutionary scenarios is challenging because of rapid rate of sequence evolution within the family and the resultant loss of phylogenetic signal. Our own preliminary investigations of Rhodopsin evolution in non-metazoans were inconclusive. Therefore, ambiguities about the provenance and function of CRM Rhodopsins currently obscure the ancestry of metazoan Rhodopsins and opsins.

The N-terminal and C-terminal protein domains of choanoflagellate, other CRM, metazoan, and additional eukaryotic GPCRs revealed uneven evolutionary conservation between GPCR families. For example, we found that the C-terminal PIPK domain of the GPCR PIPK family, the N-terminal transporter domain and C-terminal DEP domain of GPR155, and the N-terminal GOLD domain of GOST GPCRs are generally conserved in CRMs, metazoans, and other eukaryotes. In contrast, the stereotypical association of N-terminal protein domains from metazoan Glutamate Receptors is absent from non-metazoan Glutamate Receptors identified by 7TM clustering. Moreover, we observed that the N-termini of Glutamate Receptors from filastereans and holomycotans appear to possess a larger diversity of protein domains than metazoan Glutamate Receptors, suggesting functional constraint exerted on the architecture of metazoan Glutamate Receptors. This likely has implications for both the ligands that non-metazoan Glutamate Receptors may recognize and their underlying mechanism of activation (Hu et al. 2000; Jiang et al. 2004; Rondard et al. 2006; Levitz et al. 2016; Ellaithy et al. 2020b; Laboute et al. 2023).

The aGPCR family shows the greatest level of protein domain composition diversity in metazoans (Schiöth and Lagerström 2008b; Krishnan et al. 2012; Hamann et al. 2015a). Similarly, we uncovered a large diversity of N-terminal protein domain combinations in CRM aGPCRs, with a clear diversification of the repertoire in choanoflagellates and, to a lesser extent, in filastereans (Figs. 1B, 5, S4, and S7). The aGPCRs in CRMs displayed all the core structural hallmarks of metazoan aGPCRs, including a long N-terminus containing a GAIN domain and many additional ECDs. Moreover, we found that the HRM/GAIN/7TM module, previously thought to be restricted to metazoans, is also found in CRM aGPCRs. Our data support an evolutionary scenario wherein the linkage between the GAIN and 7TM domains evolved first in stem holozoans, followed by the incorporation of the HRM into the HRM/GAIN/7TM module in the progenitors of Filasterea and Choanozoa.

Despite its conservation, the function of the HRM-GAIN-7TM module is presently unclear (Shima et al. 2004; Kimura et al. 2006; Araç et al. 2012; Prömel et al. 2012b; Araç and Leon 2019). Interestingly, the HRM domain binds peptide hormones and neuropeptides in the Secretin GPCR family (Schiöth and Lagerström 2008a; Pal et al. 2012; Zhao et al. 2023). The presence of HRM-containing aGPCRs in CRMs could suggest a pre-metazoan origin of peptide-based endocrine signaling. This is reinforced by the recent finding that sponges and choanoflagellates express sequences that resemble metazoan neuropeptides and peptide hormones (Steffen et al. 2021; Yañez-Guerra et al. 2022; Zhenjian Lin, Vinayak Agarwal, Ying Cong 2024).

Finally, the diversity of protein domain architectures in the aGPCRs of choanoflagellates and metazoans suggests that aGPCR evolution was shaped by extensive domain shuffling (Gilbert 1978; Patthy 1999; Smithers et al. 2019; Patthy 2021). While some discrete protein domain associations (GAIN/7TM and HRM/GAIN/TM) or, in rare cases, most of the receptor sequence (ADGRV/subfamily VIII) survived, presumably under selection, new protein domain architectures arose, perhaps through a combination of gene duplication followed by domain shuffling. The diversification of NTF domains might have facilitated the recognition of a diversity of ligands (e.g., proteoglycans and proteins).

Of course, the aGPCRs were named for the presence of adhesion-related protein domains in the NTFs of metazoan aGPCRs, some of which support cell-cell and cell-matrix adhesion (Langenhan et al. 2013; Hamann et al. 2015b). The relevance of adhesion domains in CRM aGPCRs is intriguing and requires further investigation. Interestingly, diverse CRMs form facultative multicellular structures (Fairclough et al. 2010; Marshall and Berbee 2011; Glockling et al. 2013; Sebé-Pedrós et al. 2013; Carr et al. 2017a; Brunet et al. 2019; Dudin et al. 2019; Tikhonenkov et al. 2020; Ros-Rocher et al. 2021). Therefore, the diversification of adhesion-related protein domains in the NTFs of aGPCRs may have contributed to the evolution of multicellularity in holozoans. In addition, a recent study in sponges suggested that aGPCRs could be part of the immune response following exposure to microbial-associated molecular patterns (Pita et al. 2018).

A better appreciation of the GPCRs expressed in choanoflagellates and other CRMs is a first step to understanding cue sensing in these organisms. While environmental stimuli are known to play an essential role in controlling life history transitions in choanoflagellates, the receptors involved have yet to be characterized (Dayel et al. 2011; Alegado et al. 2012; Levin and King 2013; Woznica et al. 2017; Ros-Rocher and Brunet 2023). We anticipate that the identification of GPCR repertoires in the sister group of metazoans will open the way to the functional characterization of this superfamily of receptors in choanoflagellates, which could illuminate the origin and ancestral function(s) of key signaling pathways in metazoans.

## MATERIAL AND METHODS

### Taxon sampling

The taxa surveyed in this study are listed in Table 2.

**Table 2:**
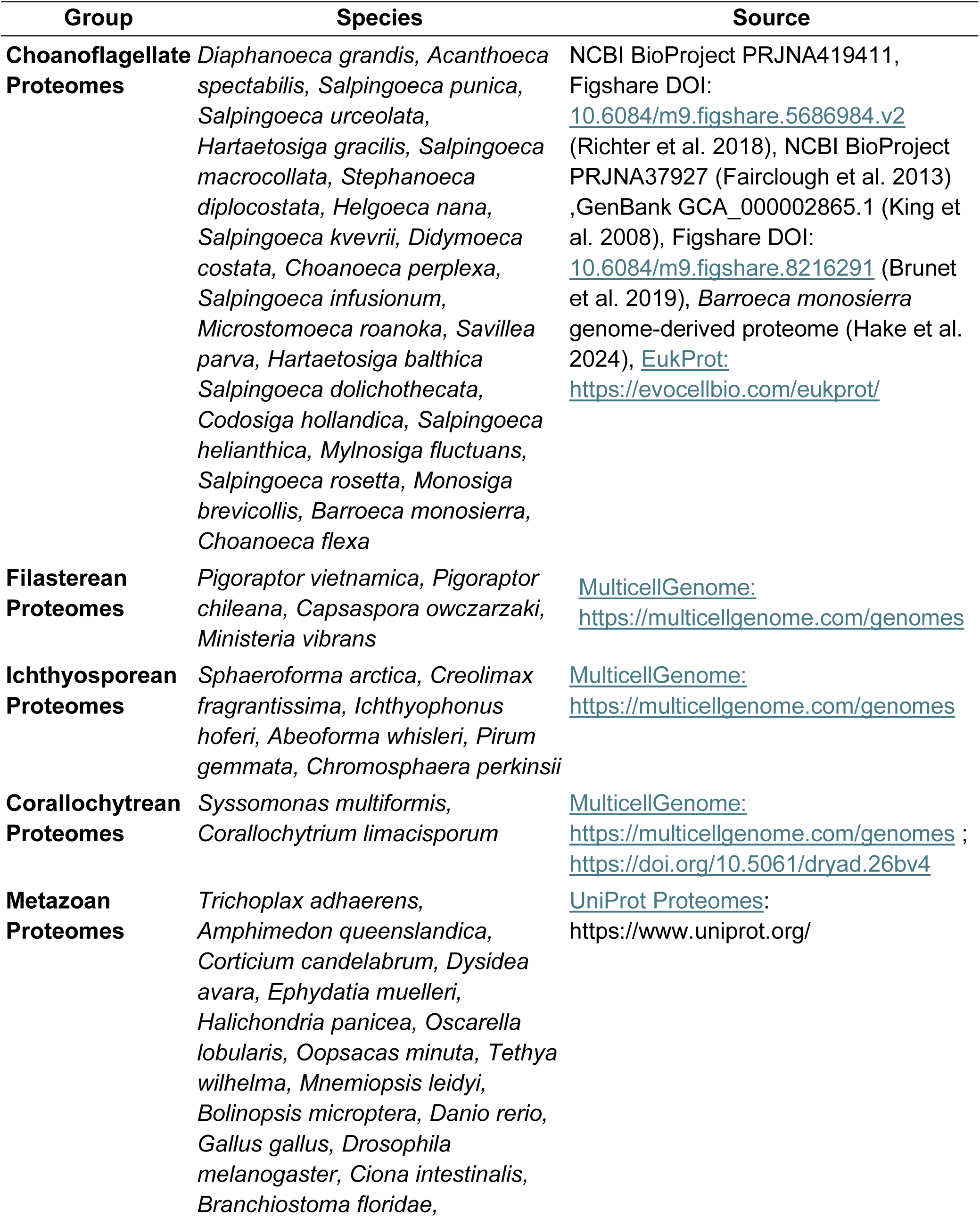
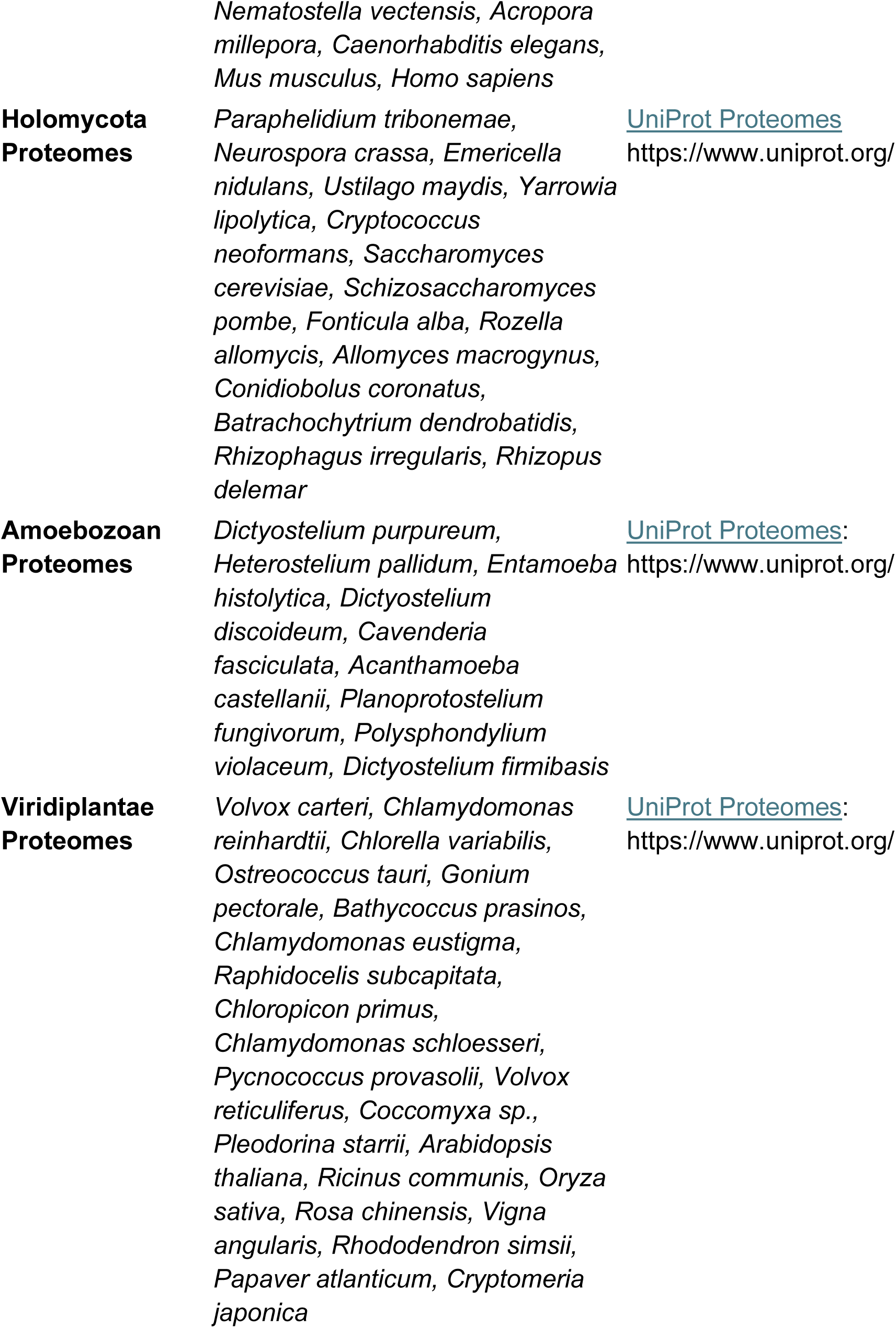
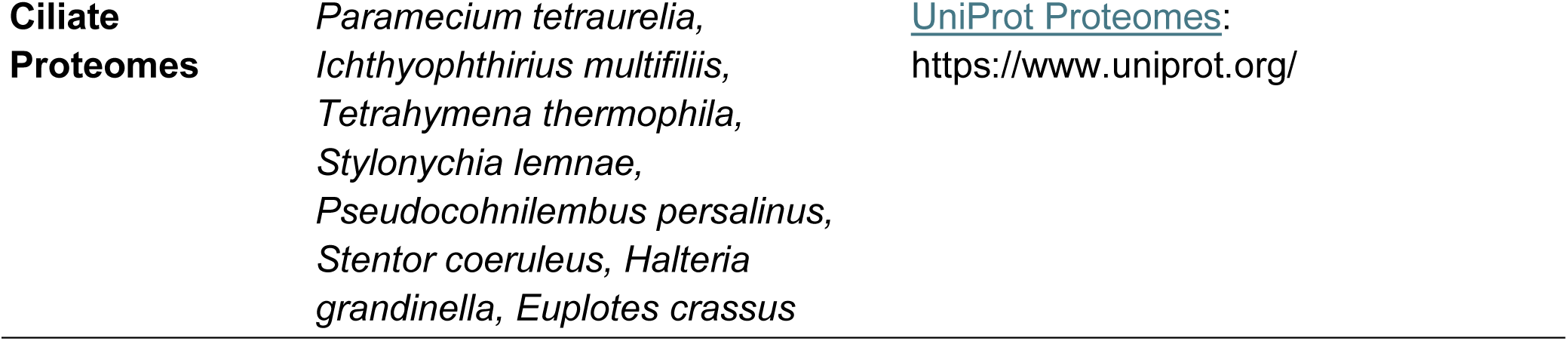
Taxon sampling and data sources for genomes and transcriptomes analyzed in this study.

### Identification of choanoflagellate GPCRs

#### GPCR mining through sequence homology-based approaches

To identify putative choanoflagellate GPCR sequences in the 23 choanoflagellate species (Table 2 and Supplementary Table 1), the choanoflagellate predicted proteomes were searched with Hidden Markov models (HMMs) unique to each of the 54 GPCR families belonging to the GPCR_A Pfam clan (CL0192; Supplementary Table 2) (Mistry et al. 2021) using the HMMER3.3 software package (cutoff threshold value: 0.01) (Eddy 2011). We recovered 1070 candidate GPCRs. In a complementary approach, we used a global HMM profile (GPCRHMM, local score) that mimics the common topology of GPCRs (Wistrand et al. 2006) and recovered 1095 candidates in our analysis. The comparison of the two sets of GPCRs revealed 381 duplicate candidates, which were merged, leaving a final set of 1784 candidates.

#### Validation of candidate GPCRs through domain composition and topology filtering

To reduce false positives and remove highly truncated sequences and possible isoforms, we submitted the candidate GPCRs to a round of filtering that combined complementary approaches. All the candidates were used as queries to perform BLASTP searches against both the NCBI and GPCRdb databases (Altschul et al. 1990; Pándy-Szekeres et al. 2023) using an e-value cut-off of 1x 10^-5^. The sequences that recovered a GPCR within the 10 top hits were kept. In addition, GPCR candidates were evaluated on the basis of their protein domain composition by combining the search tools InterProScan, CDvist, and the transmembrane domain predictor TMHMM 2.0 (Krogh et al. 2001a; Adebali et al. 2015; Blum et al. 2025). The prediction of a 7TM domain, along with additional signature protein domains, was assessed for each candidate. Protein sequences derived from genomes and transcriptomes were retained if containing 6 to 8 transmembrane helices to account for the detection of possible N-terminal signal peptides (Rutz et al. 2015) and TMHMM prediction errors. Sequences that possessed a 7TM domain but that did recover GPCRs when used as BLAST queries were subjected to a comparative structural analysis by predicting their structure using Alphafold 3 (Abramson et al. 2024) and searching for structural homologs within the Alphafold databases (van Kempen et al. 2024)see “Protein structure prediction and search” below for more details on the analysis).

Candidates with a predicted barrel shape conformation of their 7 alpha helices and that recovered GPCR structural hits (E-value < 1e^-9^) were considered GPCRs and were kept in our dataset. Finally, we removed putative splice isoforms using CD-HIT with a 90% identity threshold (Li and Godzik 2006). A total of 1113 false positives, isoforms, and highly truncated sequences were identified and removed from our dataset, leaving 671 validated choanoflagellate GPCRs in total.

#### Recovering additional choanoflagellate GPCRs using choanoflagellate GPCR BLAST queries and custom choanoflagellate GPCR HMMs

Because the HMMs used to search for choanoflagellate GPCRs in our analysis were primarily based on seed alignments of metazoan sequences, we looked for additional choanoflagellate GPCRs by using the 671 validated choanoflagellate GPCRs as queries. To this end, we simplified the choanoflagellate GPCRs dataset by clustering the 671 choanoflagellate GPCRs based on all-against-all pairwise sequence similarities (see “Clustering of the 918 validated choanoflagellate GPCRs” below to find more details about the protocol). The 671 choanoflagellate GPCRs were sorted into 18 clusters, with only 76 GPCRs that did not group with other GPCRs.

Next, we built cluster-specific GPCR HMMs for each of the 18 choanoflagellate GPCR clusters previously identified. We also built individual HMMs for the 76 GPCRs that did not fall into the 18 clusters. To this end, the 7TM domains of all the members of each of the 18 GPCR clusters were aligned independently using MAFFT (Version 7.4) with the E-INS-i algorithm (Katoh et al. 2018). The resulting 18 multisequence alignments were then used to build 18 corresponding choanoflagellate cluster-specific GPCR HMMs using the hmmbuild module of HMMER v3.3. In the case of GPCRs that did not belong to any clusters, HMMs were built based on single 7TM domains. These HMMs were used to search the choanoflagellate proteomes again. In parallel to this approach, we also selected 3 choanoflagellate GPCR sequences per GPCR cluster or a single GPCR sequence in the case of isolated GPCRs, to use as BLASTP queries against the 23 choanoflagellate proteomes (E value: 1.0e^-5^). The resulting GPCR candidates were validated through the filtering approach previously described, leaving 247 new GPCRs that were not predicted with the metazoan-biased HMMs. Added to the original 671 this yielded a total of 918 GPCRs from the 23 choanoflagellate species. The complete list of validated choanoflagellate sequences in FASTA format and the choanoflagellate-specific GPCR HMMs are available in Supplementary Files 1 and 2.

### Identification of GPCR signaling pathway components

Heterotrimeric G proteins (Gα, Gβ, and Gγ), positive regulators (Phosducin and Ric8), and negative regulators (Arrestin, GRK, RGS, and GoLoco) of G protein signaling were identified by searching choanoflagellate proteomes with Pfam HMM profiles using the HMMER3.3 software package (Eddy 2011) (cutoff threshold value: 1e^-5^) with the hmmsearch function. The following Pfam HMMs were used: PF00503 (G protein alpha subunit), PF00400 (WD domain, G-beta repeat), PF00631 (GGL domain), PF02752 (Arrestin (or S-antigen), C-terminal domain), PF00339 (Arrestin (or S-antigen), N-terminal domain), PF02188 (GoLoco motif), PF02114 (Phosducin), PF00615 (Regulator of G protein signaling domain), and PF10165 (Ric8) (Supplementary Table 2). The complete protein domain architecture of the candidates was assessed with CDvist (Adebali et al. 2015) and choanoflagellate sequences from each category were then used as BLAST queries against the 23 choanoflagellate proteomes on the EukProt database (E value: 1.0e^-5^) (Richter et al. 2022) to look for additional sequences that might not have been detected in the previous round of screening. All the sequences identified in this analysis are listed in Supplementary File 3.

### Identification of HRM-containing proteins

To identify HRM-containing proteins across eukaryotes, we searched the proteomes of metazoans, choanoflagellates, filastereans, ichthyosporeans, corallochytreans, holomycotans, amoebozoans, viridiplantae, and ciliates with the Pfam HMM profile HRM (PF02793) using the HMMER3.3 software package (cutoff threshold value: 1e^-5^) with the hmmsearch function (Eddy 2011). The complete protein domain architecture of the candidates was then further assessed using Cdvis (Adebali et al. 2015). The five HRM-containing proteins identified in the nucleariid *F.alba* are provided in Supplementary File 21.

### Identification of GPCR families in other eukaryotes

To assess the presence of Glutamate, Rhodopsin, Adhesion, and Frizzled GPCR families in other eukaryotes, we searched the relevant genomes and transcriptomes (Table 2) with their signature 7TM HMMs (Supplementary Table 2) using the HMMER3.3 software package (cutoff threshold value: 0.01) with the hmmsearch function. The candidates were then validated through the previously described domain composition and topology filtering pipeline (see “Validation of candidate GPCRs through domain composition and topology filtering”) (Supplementary File 7).

To assess the presence of other GPCR families, we searched the entire Eukprot v3 dataset (993 species) with previously published metazoan and/or non-metazoan GPCR family members as queries (threshold E value: 1.0e^-5^). Both the full-length GPCR and the extracted 7TM domains were used as search queries in this analysis. We defined hits as those with at least 70% query coverage and at least 30% sequence identity. When no hits were recovered, an additional search using their signature 7TM HMMs (Supplementary Table 2) was performed on the genomes/transcriptomes dataset (Table 2). All the candidates were then validated through a similar domain composition and topology filtering approach.

### Clustering analyses

#### Clustering of the 918 validated choanoflagellate GPCRs

To sort the choanoflagellate GPCRs into families, we created a dataset composed of all the 7TM regions extracted from the 918 choanoflagellate sequences identified in our study, along with the 7TM sequences of various metazoan, amoebozoan, stramenopile, and chlorophyte GPCRs to aid with the identification of the clusters (Supplementary File 6). The non-choanoflagellate sequences added to the dataset were either top blast hits recovered after searching the entire Eukprot v3 dataset (993 species) with choanoflagellate GPCRs as queries, or previously published and well-documented GPCR sequences from metazoans. To isolate the 7TM region, we aligned all the GPCR sequences on Geneious Prime v2024.07 using MAFTT with E-INS-I algorithm (Katoh et al. 2018), and predicted their transmembrane helices with TMHMM. All the sequences starting from the N-terminal end of the transmembrane helix 1 and finishing at the C-terminal end of the transmembrane helix 7 were kept.

Next, due to the inherent difficulties of analyzing a large set of proteins with mild sequence conservation without decreasing the accuracy of the alignment used for phylogenetic inference, we opted for an all-against-all pairwise similarity-based clustering approach to assess the diversity of GPCR families in choanoflagellates using the clustering tool CLANS 2.2.2 (Frickey and Lupas 2004a; Gabler et al. 2020). Unlike phylogenetic reconstruction, this approach becomes more accurate with an increasing number of sequences (Frickey and Lupas 2004b)(Frickey and Lupas 2004b). First, we used the CLANS web utility in the MPI Bioinformatic toolkit (https://toolkit.tuebingen.mpg.de/tools/clans) to perform an all-against-all BLAST search (with scoring matrix BLOSUM62) to obtain a matrix of pairwise sequence similarities with a cut-off value of 1e^-6^ for BLAST E-values (High Scoring Pairs with E-value higher than 1e^-6^ were not extracted). We then opened the resulting matrix in the CLANS graphical interface, and CLANS was allowed to cycle 20,000 times to optimize the graph.

Clusters were automatically detected using the convex clusters search of four or more proteins with the attraction value limit set at 0.5 standard deviation. A total of 18 choanoflagellate GPCR clusters were recovered. The clustering maps were further edited in Adobe Illustrator 2024 to modify the colors and symbols used.

#### Clustering of holozoan aGPCR N-termini

The extracellular region of the metazoan, choanoflagellate, and filasterean aGPCRs was extracted by combining the transmembrane domain prediction tool TMHMM-2.0 (Krogh et al. 2001b)(Krogh et al. 2001b) with the alignment tool MAFFT (E-INS-I algorithm) on Geneious Prime. All the regions starting at the N-terminal end of the GPCRs and finishing at the start of the first transmembrane helix were retained for downstream analysis (Supplementary File 19). The clustering analysis performed was comparable to the analysis described previously for GPCRs, with a 1e^-20^ cut-off value being used for the matrix pairwise sequence similarities.

#### Clustering of Rhodopsins

To compare the newly identified choanoflagellate Rhodopsins with the Rhodopsins found in a diversity of metazoans and other CRMs, we searched the proteomes of metazoans and CRMs (Table 2) with the Pfam HMM profile 7TM_1/Rhodopsin (PF00001) using the HMMER3.3 software package (cutoff threshold value: 1e-5) with the hmmsearch function. Candidates were then validated by assessing their InterProScan protein domain signature and BLASTing the sequences against the NCBI database. Additionally, we used the two choanoflagellate Rhodopsins – *S.macrocollata*_m.143379 and *Salpingoeca_punica*_m.44256 – to query the EukProt database (993 species), and the top blast hits with E values < 1x10^-10^ were included in our dataset. A total of 6149 validated Rhodopsins were then submitted for all-against-all pairwise comparison with a 1e^-12^ cut-off value being used for the matrix pairwise sequence similarities. All the validated Rhodopsins used in the analysis and the output of the analysis are found in Supplementary Files 11 and 12.

#### Clustering of GPCR Kinase and Kinase domains

To assess the diversity of choanoflagellate GPCR Kinase along with the diversity of their kinase domains, we extracted their 7TM domain and their kinase domain separately. To this end, we aligned all the choanoflagellate GPCR Kinase sequences on Geneious Prime using the alignment tool MAFFT (E-INS-I algorithm). The 7TM domains and the kinase domains were then predicted and extracted by running locally TMHMM-2.0 and Interproscan (PfamA) (Supplementary Files 25, 26, 28and 29). The clustering analyses were similar to the ones described previously, with cut-off values of 1e^-6^ and 1e^-20^ for the 7TM and kinase domains, respectively. In parallel, each kinase domain was also blasted against KinBase (Manning et al. 2002; Bradham et al. 2006; Goldberg et al. 2006)), the curated protein kinase dataset from www.kinase.com, for identification (Supplementary File 26).

#### Sequence alignment and phylogenetic analyses

Phylogenetic analyses of aGPCRs, Glutamate Receptors, Gα subunits, and the 7TM and Kinase domains from GPCR TK/TKL/Ks were performed in this study. To construct the holozoan aGPCR and Glutamate Receptor phylogenies, we first extracted the aGPCR and Glutamate Receptor 7TM domains by combining the transmembrane domain prediction tool TMHMM-2.0 with the alignment tool MAFFT (E-INS-I algorithm) on Geneious Prime v2024.07; only the protein sequences spanning the 7TM region were retained for downstream analysis (Supplementary Files 8, 15 and 17). To reduce the size of the large GPCR datasets while preserving their diversity, we systematically clustered similar sequences using CD-HIT (threshold 0.8; word size = 5). A total of 257 Glutamate Receptor 7TM sequences and 659 aGPCR 7TM sequences were kept for downstream analyses. To construct the phylogenies of the Kinase domain and 7TM domain from the GPCR TK/TKL/Ks, we first built a dataset including all the GPCR TK/TKL/Ks sequences identified in choanoflagellates and in sponges, as well as the GPCR TKL/Ks previously published in oomycetes and amoebozoans (Van Den Hoogen et al. 2018). We extracted the 7TM domain and Kinase domain from each sequence by combining the transmembrane domain prediction tool TMHMM-2.0 and the protein domain prediction tool InterProScan with the alignment tool MAFFT (E-INS-I algorithm) on Geneious Prime v2024.07 (Supplementary Files 30 and 32). We then aligned the aGPCR, Glutamateand GPCR TK/TKL/K Receptor 7TMs, the GPCR TK/TKL/Ks Kinase domain, or the full-length Gα sequences using MAFFT with the E-INS-I algorithm. The resulting alignments were then used for Maximum-likelihood and/or Bayesian inference of phylogenies (Fig. 3B, Fig. 5A, Fig. S3D, Fig. S6A, and Fig. S17F and G; Supplementary Files 5, 9, 16,18, 31, and 33). We built Maximum-likelihood phylogenies with IQ-TREE web server (http://iqtree.cibiv.univie.ac.at/; (Trifinopoulos et al. 2016)) using ModelFinder (Kalyaanamoorthy et al. 2017) and 2000 Ultrafast Bootstraps (UF-boot) (Minh et al. 2013) or 2000 iterations of SH-aLRT. Bayesian inference was performed using MrBayes v3.2.7a (Huelsenbeck and Ronquist 2001; Ronquist et al. 2012) on CIPRES Science Gateway (Miller et al. 2010). The posterior probability of the trees was estimated using Markov Chain Monte Carlo (MCMC) analysis and a fixed LG amino acid substitution model (Aamodelpr = Fixed(LG)). The gamma-shaped model was used to estimate the variation of evolutionary rates across sites (set rates = Gamma). MCMC analysis was set to run for 6,000,000 generations and every 120^th^ tree was sampled. Diagnostics were calculated for every 1000 generations (diagnfreq = 1000) to analyze the convergence of the two independent runs starting from different random trees. To terminate the MCMC generations, a stop rule was applied, and the convergence was analyzed until the average standard deviation of split frequencies dropped below 0.01. To ensure that the parameter estimates were only made from data drawn from distributions derived after the MCMCs had converged, we discarded the first 25% of the sampled trees in the burnin phase using relative burnin setting (relburnin = yes and burninfrac = 0.25). A consensus tree was built from the remaining 75% of the sampled trees with the MrBayes sumt command using the 50% majority rule method. Trees were visualized using iTOL (https://itol.embl.de/) (Letunic and Bork 2024) and further edited in Adobe Illustrator 2024.

### Protein Domain Search

To identify and compare the protein domains found in the extracellular region of aGPCRs and Glutamate Receptors across a range of eukaryotes, we first assessed the aGPCR and Glutamate Receptor repertoires from various metazoans, filastereans, ichthyosporeans, corallochytreans, holomycotans, amoebozoans, viridiplantae, and ciliates (see Table 2). Briefly, the proteomes were searched for aGPCR and Glutamate Receptors with the Pfam HMM profiles – 7TM_2/Adhesion (PF00002) and 7TM_3/Glutamate (PF00003) – respectively, using the HMMER3.3 software package (cutoff threshold value: 1e^-5^) with the hmmsearch function.

Candidates were further verified by assessing their InterProScan protein domain signature and BLASTing the sequences against the NCBI database. The validated aGPCR and Glutamate Receptors from each clade (choanoflagellates, metazoans, filastereans, ichthyosporeans, corallochytreans, holomycotans, amoebozoas, viridiplantae, and ciliates), were then analyzed in batch using CDvist with default parameters to assess their protein domain composition (Fig. 3C and Fig. S7; Supplementary Files 10 and 19).

### Protein structure prediction and search

The protein structures of full-length GPCRs and/or extracted 7TM domains were predicted using the AlphaFold 3 server with default parameters (https://alphafoldserver.com/; (Abramson et al. 2024)). Five models were generated per input, of which only the top-ranked prediction (based on the ranking_score metric) was selected for downstream analyses. The quality of the predicted structure was then assessed based on the pLDDT confidence score. Only models with pLDDT scores above 70 were retained. Structures were analyzed and figures were prepared with ChimeraX (Meng et al. 2023).

To find structural homologs, we searched the Alphafold-predicted structures against the AFDB-PROTEOME, AFDB-SWISSPROT, and AFDB50 databases using the search program Foldseek (https://search.foldseek.com/search; (van Kempen et al. 2024)) with default parameters.

Structural top hits with a pLDDT confidence score above 70 in the aligned region and an associated E-value < 1e^-9^ were considered putative structural homologs.

### Logo analysis

To build GAIN domain sequence logos (Fig. S9B), we isolated the C-terminal region of the GAIN domain from 301 choanoflagellate, 99 filasterean, 1 corallochytrean, and 30 murine aGPCRs (Supplementary file 21). To do so, we aligned separately the aGPCR sequences from the four datasets using MUSCLE v5 (Edgar 2022) on Geneious Prime, and the region of the C-terminal GAIN domain starting with the first N-terminal conserved Cysteine and finishing with the TA consensus “TxFAVLM” was extracted and kept for downstream analysis. We then trimmed the aligned sequences with ClipKIT (Steenwyk et al. 2020) using the GAPPY mode (with the default value:0.9). The resulting processed alignments were analyzed with WebLogo3 (https://weblogo.threeplusone.com/) (Crooks et al. 2004).

## ACKNOWLEDGEMENTS

We are grateful to Demet Araç, Thibaut Brunet, Daniel Richter, and Iñaki Ruiz-Trillo for feedback on the manuscript. We thank the whole King lab for valuable feedback throughout the project and Flora Rutaganira for advice about categorizing kinases. This work was supported by a Human Frontier Science Program long-term fellowship (LT 000919/2020-L) to A.G.D.L.B and an Investigator Award from the Howard Hughes Medical Institute (N.K.).

## DATA AVAILABILITY

**Supplementary Table 1**: Choanoflagellate genome and transcriptome accession numbers.

**Supplementary Table 2**: List of identifiers of all Pfam HMMs used in this study.

The Supplementary Files below are all available on Figshare: https://doi.org/10.6084/m9.figshare.28801289.v1

**Supplementary File 1**: 918 choanoflagellate GPCR sequences identified in this study.

**Supplementary File 2**: Choanoflagellate cluster-specific GPCR HMMs.

**Supplementary File 3**: Choanoflagellate heterotrimeric G proteins, positive and negative regulators of G protein signaling identified in this study.

**Supplementary File 4**: List of all Gα sequences used for phylogenetic reconstruction in Figure S3D.

**Supplementary File 5**: Maximum-likelihood phylogeny of choanoflagellate and metazoan Gα sequences. Related to Figure S3D.

**Supplementary File 6**: Complete set of 7TM sequences used for the clustering analysis related to Figure 1B.

**Supplementary File 7**: Glutamate, Rhodopsin, Adhesion, and Frizzled receptors identified in opisthokonts and other eukaryotes; related to Figure 2.

**Supplementary File 8**: Glutamate Receptor 7TM sequences used in the phylogenetic analysis related to Figure 3B.

**Supplementary File 9**: Fully annotated versions of the Glutamate Receptor phylogeny depicted in Figure 3B.

**Supplementary File 10**: List of opisthokont Glutamate Receptor sequences used for the analysis related to Figure 3C.

**Supplementary File 11**: Set of 6152 Rhodopsin sequences used in the clustering analysis related to Figure 4B.

**Supplementary File 12**: Raw cluster analysis CLANS file; related to Figure 4B.

**Supplementary File 13**: Seed alignment of the Rhodopsin sequences related to Figure 4E.

**Supplementary File 14**: Seed alignments downloaded from GPCRdb used to build human consensus sequences for the Aminergic, Peptide, and Lipid Rhodopsin subfamilies. Related to Figure 4E.

**Supplementary File 15**: List of all the aGPCR 7TM sequences used to build the two phylogenies depicted in Figure S6A and B.

**Supplementary File 16**: Fully annotated versions of the two phylogenies depicted in Figure S6A and B.

**Supplementary File 17**: List of all the aGPCR 7TM sequences used to build the phylogeny depicted in Figure 5A.

**Supplementary File 18**: Fully annotated versions of the aGPCR phylogeny depicted in Figure 5A.

**Supplementary File 19**: List of all the aGPCR NTF sequences used in Figure S7 and in the clustering analysis depicted in Figure S8.

**Supplementary File 20**: List of choanoflagellate aGPCRs without detectable GAIN domain related to Figure S9A.

**Supplementary File 21**: Extracted C-term GAIN sequences and seed alignments used to create the logos related to Figure S9C.

**Supplementary File 22**: Sequences of the five HRM-containing proteins identified in *Fonticula alba*; related to Figure S10D.

**Supplementary File 23**: GPR157 homologs identified in diverse eukaryotes; related to Figure S11A.

**Supplementary File 24**: Hi-GOLD homologs identified in diverse eukaryotes.

**Supplementary File 25**: Extracted kinase domain sequences from choanoflagellate GPCR Kinase used for the clustering analysis in Figure S17B.

**Supplementary File 26**: Raw cluster analysis CLANS file; related to Figure S17B.

**Supplementary File 27**: Top Blast hits of the choanoflagellate kinase domains analyzed in Fig. S17B searched against KinBase.

**Supplementary File 28**: Extracted 7TM domain sequences from choanoflagellate GPCR TKs, TKLs, and Ks used in the clustering analysis in Figure S17C.

**Supplementary File 29**: Raw cluster analysis CLANS file; related to Figure S17C.

**Supplementary File 30**: Extracted Kinase domains from choanoflagellate, sponge, oomycete and amoebozoan GPCR TK/TKL/Ks used to build the phylogeny depicted in Figure S17F.

**Supplementary File 31**: Raw Newick format of the tree depicted in Figure S17F.

**Supplementary File 32**: Extracted 7TM domains from choanoflagellate, sponge, oomycete and amoebozoan GPCR TK/TKL/Ks used to build the phylogeny depicted in Figure S17G.

**Supplementary File 33**: Raw Newick format of the tree depicted in Figure S17G.

**Supplementary File 34**: Frizzled/Smoothened sequences identified in the corallochytrean *S. multiformis*; related to Figure S19.

## Supplemental Text

### Conservation of the GPCR signaling pathway in choanoflagellates

In Metazoa, the binding of a ligand to the N-terminus or extracellular loops of a GPCR triggers a conformational change that initiates the rest of the GPCR signaling pathway (Hilger et al. 2018; Wootten et al. 2018; Hauser et al. 2021). Ligand-bound GPCRs act as guanine nucleotide exchange factors (GEFs) for the GDP-bound Gα subunit, catalyzing the exchange of GDP for GTP and, thereby, promoting the dissociation of activated Gα-GTP from Gβγ dimers (Fig. S3A, (Hilger et al. 2018; Weis and Kobilka 2018; Wootten et al. 2018)). Both activated Gα-GTP and free Gβγ dimers can transduce the signal independently to downstream effectors (Marinissen and Gutkind 2001; Neves et al. 2002; Dupré et al. 2009). Ric8 proteins and Phosducin facilitate the folding and targeting at the membrane of Gα subunit and Gβγ dimers, respectively (Willardson and Howlett 2007; Srivastava et al. 2019). Overstimulation of GPCR signaling is detrimental to the cell, and various mechanisms quickly turn off GPCR activity in Metazoa.

Signal termination is controlled in part by the hydrolysis of Gα-GTP to Gα-GDP, which reconstitutes the inactive Gαβγ heterotrimer. The rate of Gα-GTP hydrolysis is a consequence of the intrinsic GTPase activity of Gα and can be enhanced by the action of regulators of G protein signaling (RGS), proteins that act to accelerate the GTPase activity (O’Brien et al. 2019). In addition, GoLoco motif proteins are negative regulators of Gα-GTP signaling, as they slow the spontaneous release of GDP from the Gα subunit (Willard et al. 2004; Sato et al. 2006). Another major axis to terminate GPCR signaling involves the rapid phosphorylation of the ligand-bound receptor by GPCR-Kinases (GRKs) and the subsequent binding of Arrestin proteins that selectively recognize active phosphorylated receptors (Rajagopal and Shenoy 2018; Gurevich and Gurevich 2019). Arrestins not only abolish G protein-mediated signaling but also act as a scaffold to facilitate multiple downstream signaling pathways and trigger the endocytosis of the receptor (Fig. S3A, (Bagnato and Rosanò 2019)).

We found that the heterotrimeric G proteins Gα, Gβ, and Gγ, along with positive regulators (Ric8, Phosducin) and negative regulators (GRK, Arrestin, RGS, GoLoco) of GPCR signaling, were detected in most choanoflagellates, with some exceptions in a few species (Fig. S3B, Supplementary File 3). Notably, we failed to detect Gγ and GRK in nearly half of the choanoflagellates, which could either suggest real losses, false negatives, or a divergence beyond recognition of these genes in the species concerned.

We decided to assess further the diversity of Gα subunits, the best-characterized transducers of metazoan GPCR signaling, in choanoflagellates. In metazoans, Gα subunits are classified into five main families (Gαs, Gαi/o, Gαq/11, Gα12/13, and Gαv), and members of each family interact with different effectors to produce distinct cellular responses (Fig. S3C, (Marinissen and Gutkind 2001; Neves et al. 2002; Oka et al. 2009; Oka and Korsching 2009; Doktorgrades and Obaid 2022; Zhang et al. 2024)). Although all major G protein Gα classes (Gα_s_, Gα_q/11_, Gα_12/13_, Gα_v_, and Gα_i/o_) likely originated prior to the diversification of metazoans from the rest of holozoans (De Mendoza et al. 2014; Krishnan 2015; Lokits et al. 2018), our phylogenetic analysis recovered only three Gα classes in choanoflagellates (Gα_s_, Gα_q/11_, and Gα_v_) (Fig. S3D and Supplementary Files 4 and 5). Interestingly, Gα_s_ subunits, previously thought to be lost in choanoflagellates, were detected in four closely related species (*S. kvevrii*, *S. urceolata*, *S. macrocollata*, *S. punica*). Based on their functions in metazoans, the activation of these Gα subunits could modulate protein phosphorylation (Gαs and Gαq/11), calcium signaling (Gαq/11), and ion homeostasis (Gαv) downstream of the GPCRs in choanoflagellates (Fig. S3C and S3D) (Marinissen and Gutkind 2001; Neves et al. 2002; Abu Obaid et al. 2024).

Overall, our results suggest that the GPCR signaling pathway is conserved in most choanoflagellate species, providing additional evidence that metazoan-like G protein signaling is part of the repertoire of signaling activities in choanoflagellates (De Mendoza et al. 2014; Krishnan et al. 2015; Lokits et al. 2018).

**Figure S1:**
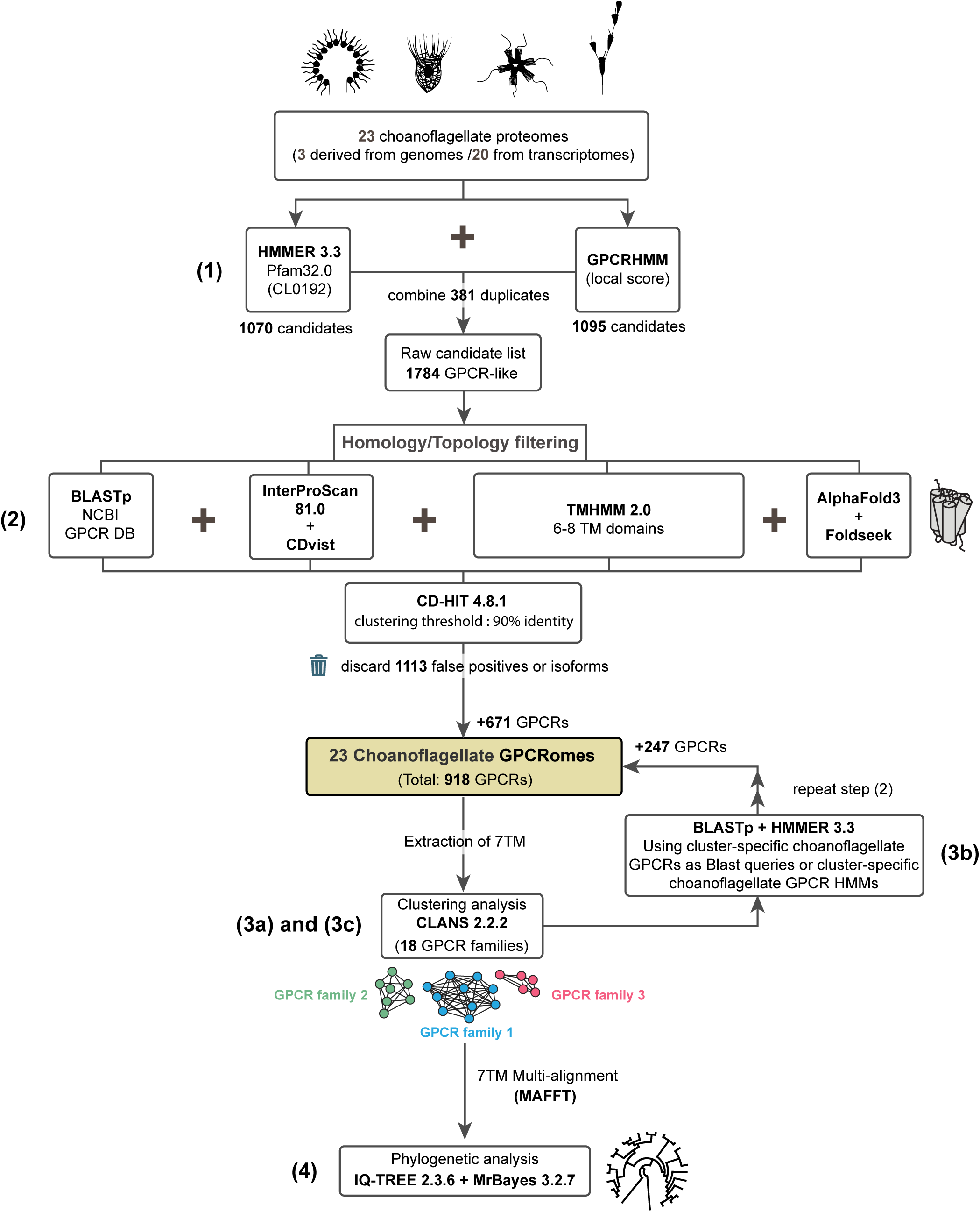
Pipeline for identifying GPCRs in choanoflagellate genomes and transcriptomes. **(1)** GPCR candidates were searched in the genome- and transcriptome-derived proteomes of 23 choanoflagellate species using Hidden Markov model(s) (HMMs) unique to 54 GPCR families in eukaryotes (GPCR_A Pfam clan CL0192) or a global HMM profile that matches the common topology of GPCRs irrespective or their families (GPCRHMM). The comparison of the two sets of candidate GPCRs obtained through these two independent approaches revealed 381 duplicate candidates, which were merged, leaving a final set of 1784 candidates. **(2)** The 1784 candidate choanoflagellate GPCRs were then subjected to a validation step where false negatives and highly truncated sequences were filtered out using the search tools BLASTp, Interproscan, CDvist, and the transmembrane domain predictor TMHMM (see Methods). Alphafold-predicted structures of the GPCR candidates and subsequent structure homology search complemented the analysis. Finally, we used CD-HIT to discard isoforms. A total of 1113 sequences were removed from our dataset, leaving 671 validated choanoflagellate GPCRs. **(3a, 3c)** To assess the diversity of choanoflagellate GPCRs, we then clustered the 671 GPCRs based on all-against-all pairwise sequence similarities using the clustering tool CLANS. Most GPCRs (629 receptors) were sorted into 18 clusters, with a minority of them (76 receptors) sharing no sequence similarities with any other GPCRs in the dataset. The same protocol (3a) was re-iterated in (3c), after refining the choanoflagellate GPCR dataset (3b) and including diverse metazoan, amoebozoan, stramenopile, alveolate, and chlorophyte GPCRs to the analysis. **(3b)** Because the HMMs used in our first round of screening were mostly based on seed alignments of metazoan sequences, we built choanoflagellate cluster-specific GPCR HMMs for each of the 18 choanoflagellate GPCR clusters previously identified (3a) to detect possible additional choanoflagellate GPCRs (Supplementary File 2). We also built individual HMMs for the GPCRs that did not fall into the 18 clusters. These HMMs were used to search the 23 choanoflagellate proteomes again. In parallel, we also selected choanoflagellate GPCR sequences from each GPCR cluster or single GPCR sequences in the case of GPCRs that did not group with other GPCRs, to use as BLAST queries against the 23 choanoflagellate proteomes. The resulting candidates were run through the filtration process in step (2), leaving 247 new GPCRs that were not predicted with the metazoan-biased HMMs. Added to the original 671, this gave a total of 918 GPCRs from the 23 choanoflagellate species (Supplementary File 1). **(4)** Phylogenetic trees were inferred for various choanoflagellate GPCR families, including aGPCRs and Glutamate Receptors, using IQ-TREE and MrBayes.

**Figure S2:**
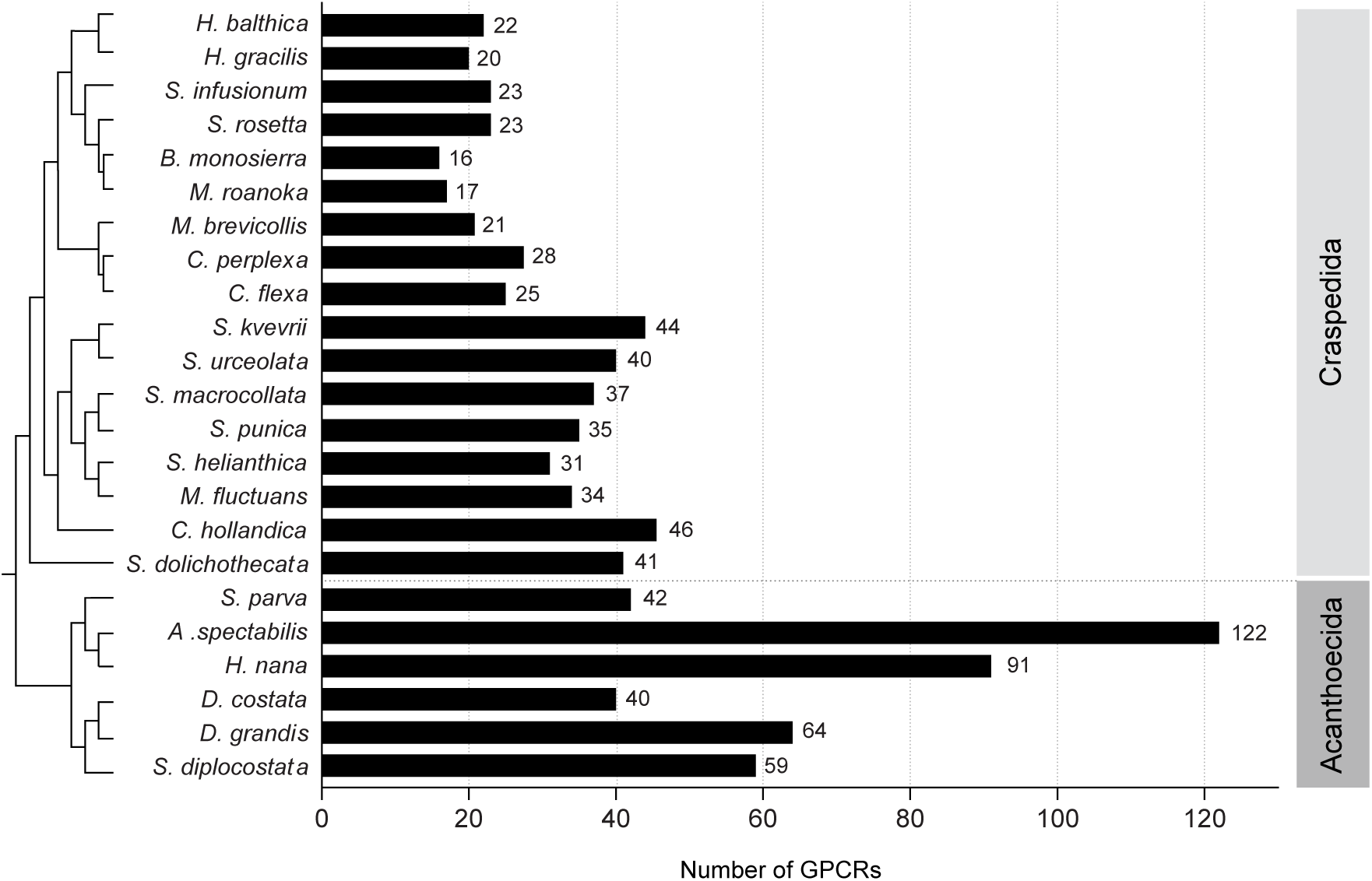
Total number of GPCRs across choanoflagellates varies by species and seems to correlate with phylogenetic affiliation. GPCR numbers tend to be higher in acanthoecids than in craspedids. Shown are GPCR numbers per species, mapped to a consensus phylogeny adapted from (Carr et al. 2017b; Ginés-Rivas and Carr 2025b). Choanoflagellate GPCR numbers range from 16 in *B. monosierra* to 122 in *A. spectabilis.* These are approximate numbers, as the transcriptomes may be incomplete and/or the predicted proteomes may include splice variants.

**Figure S3:**
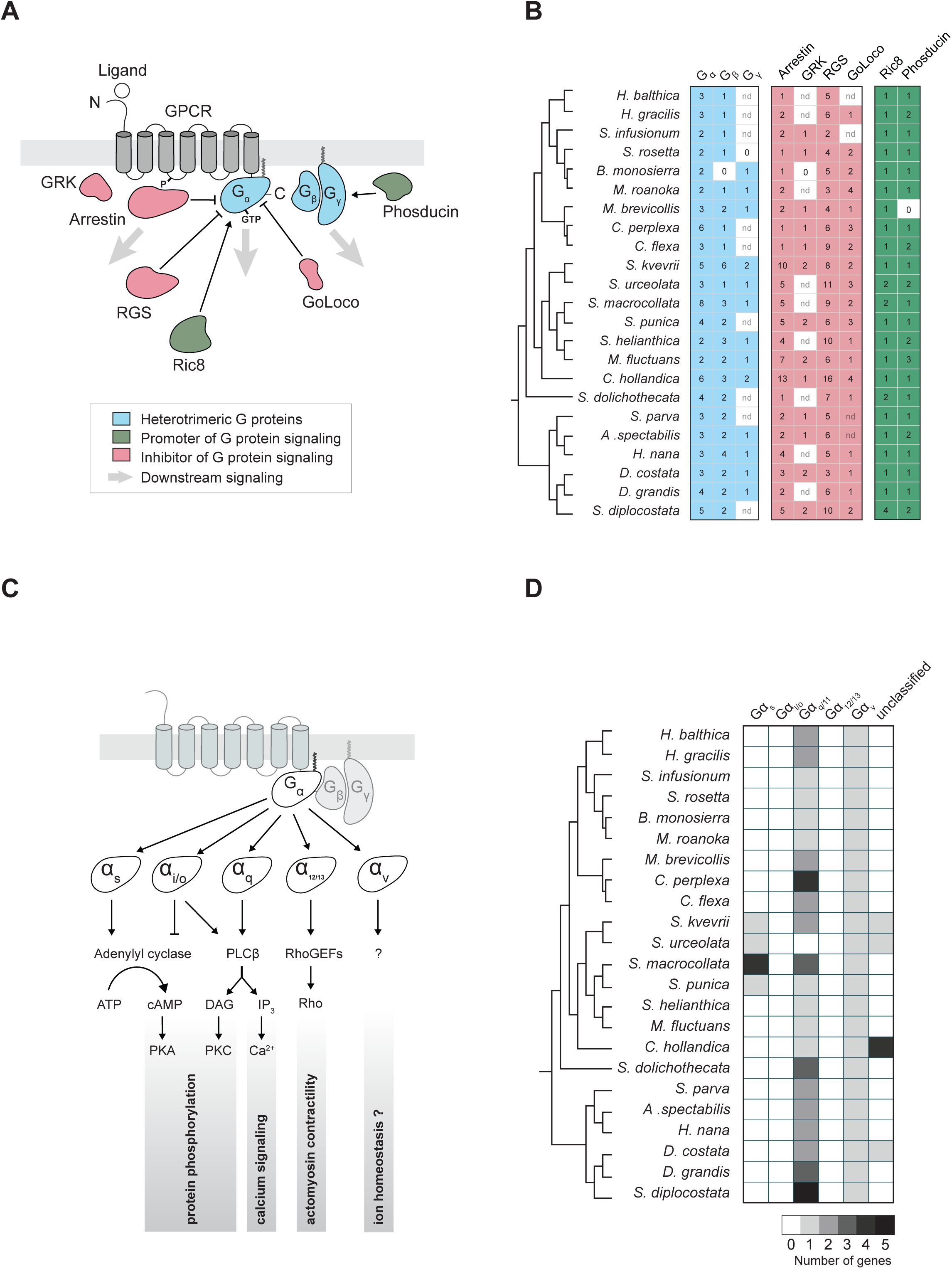
Conservation of metazoan GPCR transducers in choanoflagellates. **(A)** Schematic representation of the core GPCR signaling pathway and associated regulators. Upon binding to a ligand, GPCRs activate downstream heterotrimeric G proteins (blue) that dissociate, giving free Gα and Gβγ subunits that transduce the signal independently from each other (grey arrows). Additional regulators ensure the fine-tuning of GPCR signaling by either promoting G protein signaling (green) or inhibiting it (red) (see Supplementary text for more information). While heterotrimeric G proteins are the canonical signal transducers downstream of GPCR activation, Arrestins act as important signaling hubs (Gurevich and Gurevich 2019). **(B)** Phylogenetic distribution of heterotrimeric G proteins (Gα, Gβ, and Gγ; blue), negative regulators of G protein signaling (Arrestin, GRK, RGS, and GoLoco; red) and positive regulators of G protein signaling (Ric8 and Phosducin; green) within choanoflagellates. The number of proteins recovered in our analysis that correspond to each of these different categories is specified in the matrix for each choanoflagellate species. The GPCR signaling pathway is conserved in most choanoflagellate species. nd = not detected in lineages for which only transcriptome data are available. For species with both transcriptome and genome data (*S. rosetta*, *B. monosierra*, and *M. brevicollis*) (King et al. 2008; Fairclough et al. 2013; Hake et al. 2024), failure to detect a GPCR subfamily member is indicated with a “0”. All the choanoflagellate sequences identified in each of these categories are provided in Supplementary File 3. **(C)** Schematic representation of major pathways activated by different Gα subunits. Gα proteins can be subdivided into five main families (Gα_s_, Gα_i/o_, Gα_q/11_, Gα_12/13_, and Gα_v_) with different signaling properties (Marinissen and Gutkind 2001; Oka and Korsching 2009; Feng et al. 2022; Liu et al. 2024). Gα_s_ proteins activate Adenylyl cyclase, resulting in an accumulation of intracellular cAMP and activation of protein kinase A (PKA). In contrast, the activated Gα_i/o_ inhibits Adenylyl cyclase. Gα_i/o_ proteins also stimulate phospholipase C-β (PLCβ) that generates diacylglycerol (DAG) and inositol trisphosphate (IP_3_), eventually regulating Ca2+ signal and protein kinase C (PKC) activity; a function shared with Gα_q/11_ subunits. Gα_12/13_ controls actomyosin contractility through the activation of the Rho/Rho kinase signaling pathway. Little is known about signal transduction downstream of Gα_v_ which could control ion homeostasis (Abu Obaid et al. 2024). **(D)** Phylogenetic distribution of the five Gα subunit families within choanoflagellates. While Gα_q/11_ and Gα_v_ are shared by all 23 choanoflagellate species, Gα_i/o_ and Gα_12/13_ were not detected in our dataset. We identified Gα_s_ subunits in a monophyletic clade encompassing *S. kvevrii*, *S. urceolata*, *S. macrocollata*, and *S. punica*. Failure to detect Gα_s_ in *S. helianthica* and *M. fluctuans* could be due to false negatives or a secondary loss in the ancestor or these two sister species. Gα proteins that were not assigned to any of these five families were counted as unclassified. The assignment of choanoflagellate Gα proteins to one of the five Gα subunit families was based on the maximum likelihood phylogenetic tree inferred by the G protein alpha subunit from both choanoflagellates and known Gα metazoan subunits (Supplementary Files 4 and 5).

**Figure S4:**
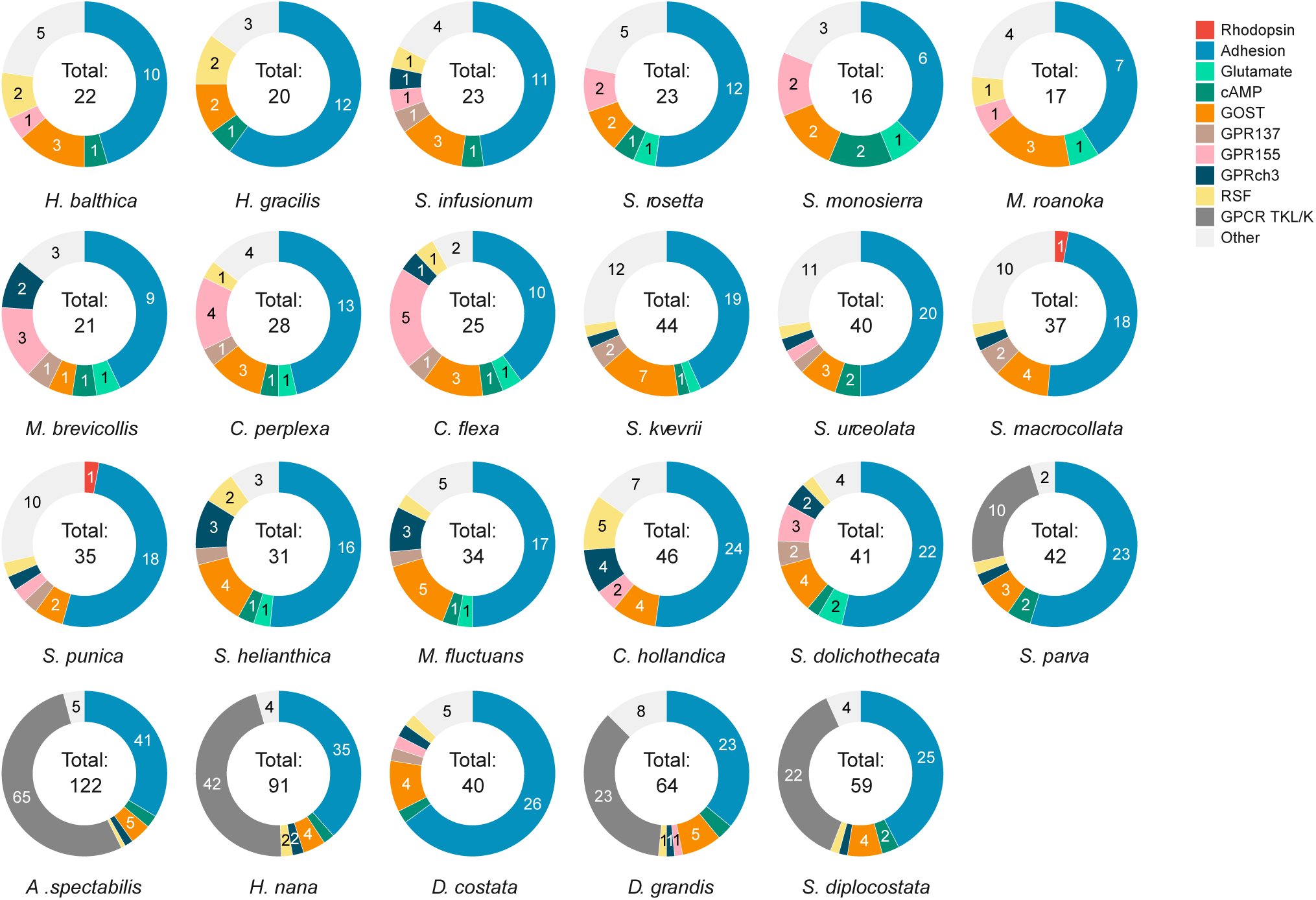
aGPCRs dominate the GPCRome of most choanoflagellates analyzed. Donut charts show the number and proportion of genes in the main GPCR families – Rhodopsin, Adhesion, Glutamate, cAMP, GOST (including TMEM87, GPR107/108, TMEM145, GPR180, and Hi-GOLD), GPR137, GPR155, GPRch3, RSF, GPCR TKL/K and Other GPCRs – for each of the 23 choanoflagellate species analyzed. Adhesion receptors represent, on average, 47% of the GPCRs encoded in these choanoflagellate species. The presence of a large repertoire of GPCR TKL/Ks in acanthoecids contributes, in part, to the larger size of their GPCRomes. The total number of GPCRs detected is shown in the center.

**Figure S5:**
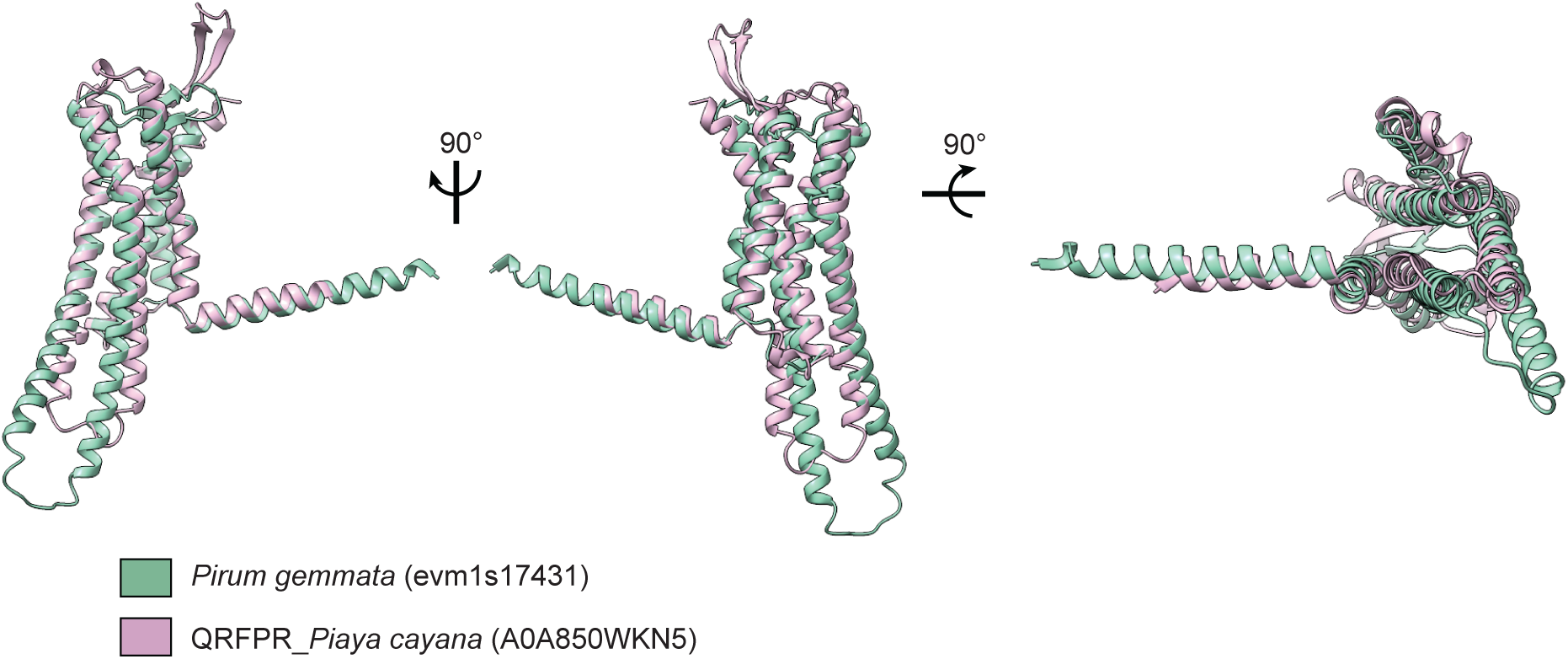
Ichthyosporean Rhodopsin shares structural similarities with metazoan peptide receptor Rhodopsin. Predicted structural homology of model *Pirum_gemmata*_evm1s17431 (green) with Foldseek top hit (E-value: 4.92e^-9^) Pyroglutamylated RFamide peptide receptor (QRFPR)_Piaya_cayana (AF-A0A850WKN5; pink). A cytoplasmic helix 8 (H8), found in most metazoan Rhodopsins, is also predicted in the ichthyosporean rhodopsin. Low confidence regions (>70 pLDDT) were removed for clarity in both models. View of the superimposed models from the plane of the membrane (left and center) and the top (right).

**Figure S6:**
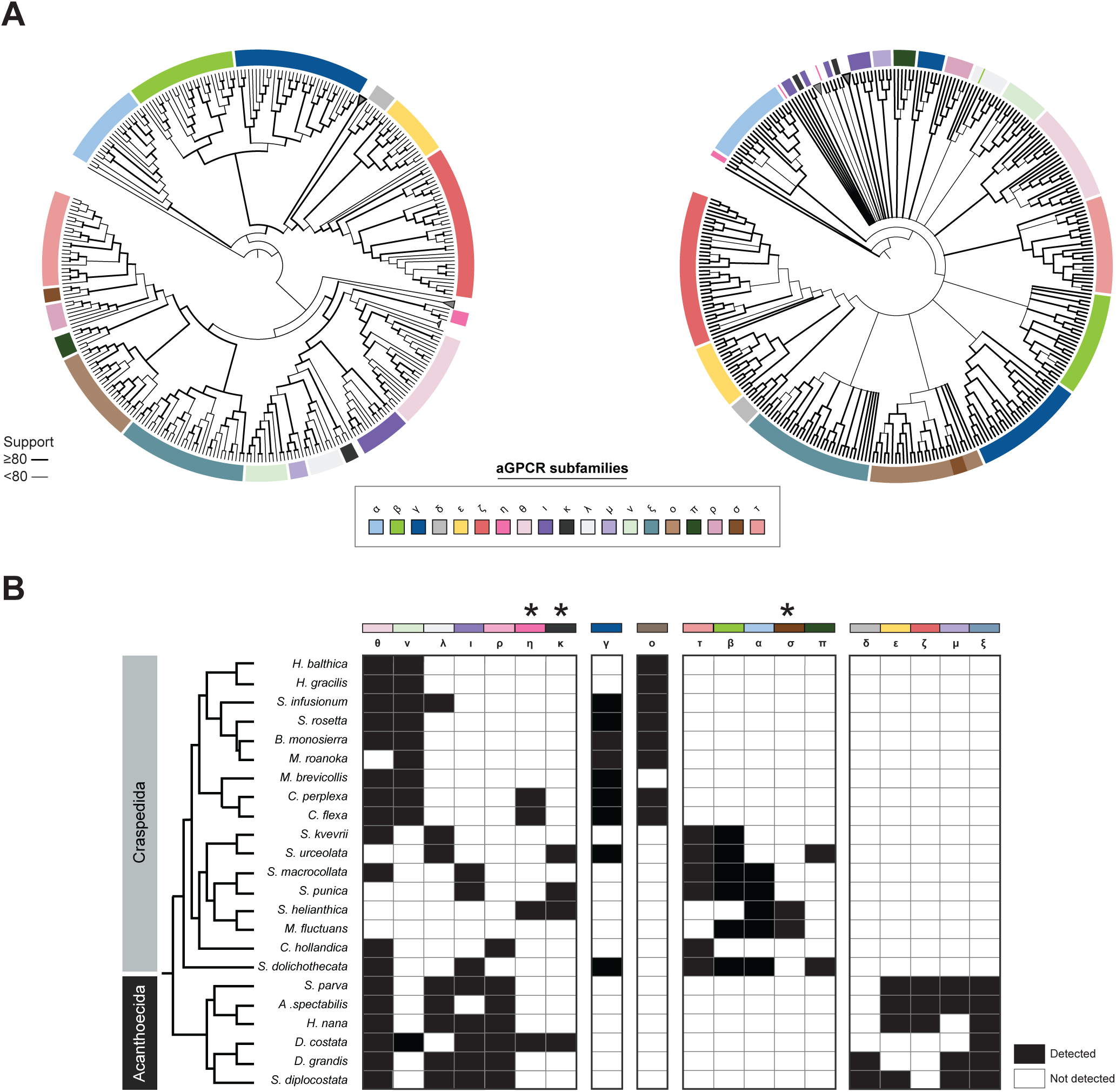
Diversity of aGPCR subfamilies in choanoflagellates. **(A)** Phylogenetic analyses of the 7TMs of choanoflagellate aGPCRs reveal the presence of at least 19 subfamilies, labeled α-τ. Maximum-likelihood inference (left tree) and Bayesian inference (right tree) include full 7TM sequences of 229 choanoflagellate aGPCRs along with a 7TM sequence of a ciliate aGPCR (Stentor_coeruleus_OMJ80129.1_SteCoe_19670), used as an outgroup to root the trees. The 7TM sequences were aligned with MAFFT V7.463 using the E-INS-I algorithm, and phylogenies were built using either IQ-TREE or MrBayes v3.2.7a (see Methods). The two phylogenies cross-verified the grouping of most aGPCRs into the same subfamilies (depicted with the same color code in the two trees; see legend). The evolutionary relatedness between most aGPCR subfamilies could not be determined with the Bayesian approach (polytomy). The width of branches indicates scales with UFboot support (left tree) or Bootstrap support (right tree) for the ancestral node. Branch lengths do not scale with evolutionary distance in this rendering. Clades poorly supported (< 70% Bootstrap support) or composed of sequences from a single choanoflagellate species were collapsed in this rendering (grey triangles). All the sequences used to build the trees are listed in Supplementary File 15. The fully annotated version of the two phylogenies, including bootstrap values, branch lengths, and all species names, are found in Supplementary File 16. **(B)** aGPCR subfamilies differ in their phylogenetic distribution across the choanoflagellate diversity. Subfamilies θ, ν, λ, ι, ρ, η, and κ are detected in diverse craspedid and acanthoecid choanoflagellates – therefore they were probably present in stem choanoflagellates. Notably, the subfamily θ is the most widespread of all choanoflagellate aGPCR subfamilies and is conserved in filastereans and metazoans (Ansel et al. 2024). In contrast, subfamilies γ, ο, τ, β, α, σ, and π are restricted to craspedids. While subfamily III is broadly conserved across the diversity of craspedids, subfamilies ο, τ, β, α, σ, and ξ are confined to distinct subsets of craspedids. Finally, subfamilies δ, ε, ζ, η, and ξ were only observed in acanthoecids and, therefore, might be acanthoecid-specific. Asterisks (**\***) indicate aGPCR subfamilies that were recovered in only one of the two phylogenetic reconstructions (see (**A**)).

**Figure S7:**
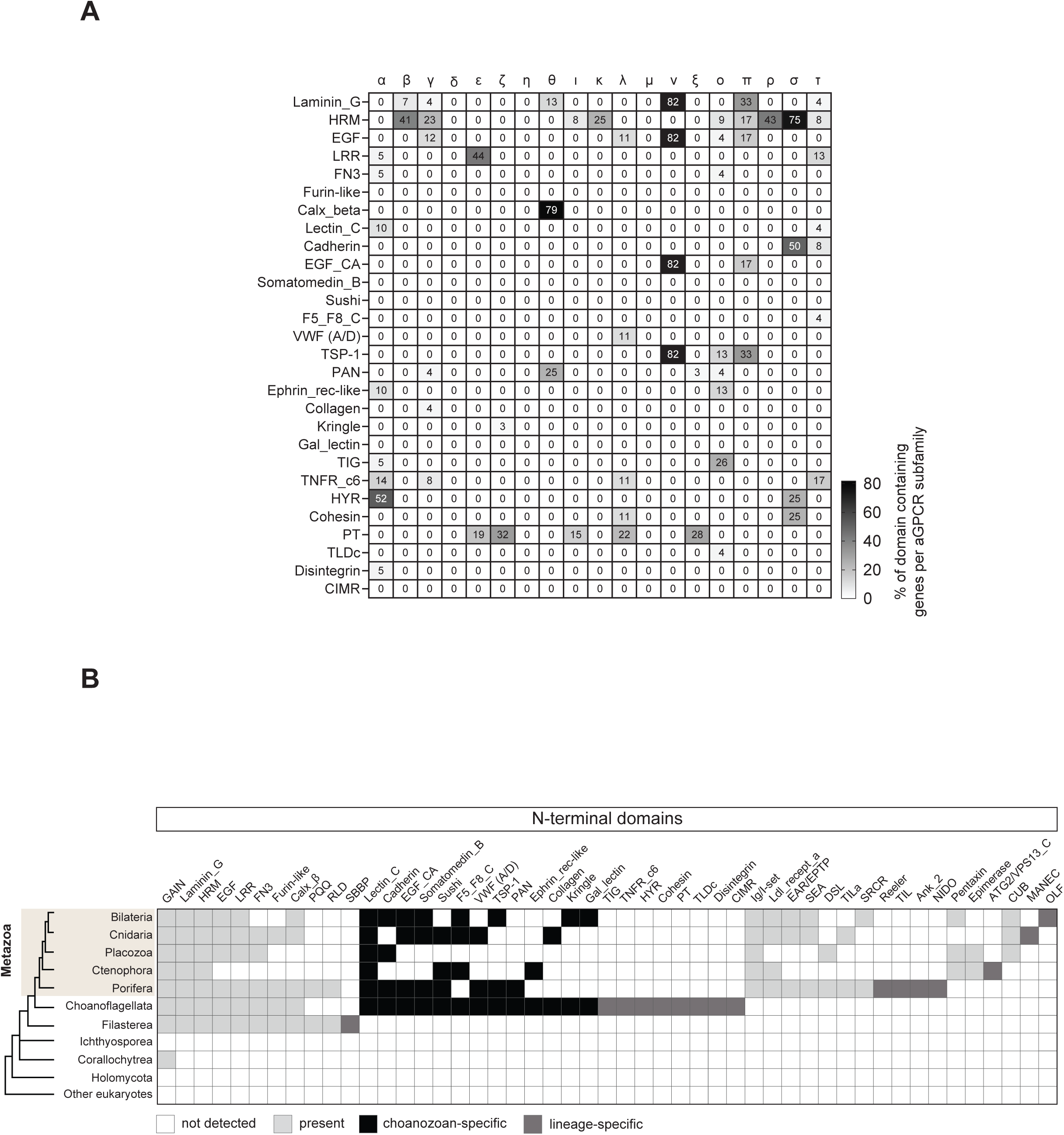
Diversity and evolution of aGPCR extracellular protein domains in choanoflagellates and other opisthokonts. **(A)** Subfamilies of choanoflagellate aGPCRs exhibit distinct N-terminal protein domain signatures. The matrix summarizes the systematic identification of the protein domains (rows) present in the extracellular region of the members of the 19 choanoflagellate aGPCRs subfamilies previously established (columns) (**Fig. S6**). Percentages represent the number of domain-containing genes per aGPCR subfamily. For example, 52% of aGPCRs from the subfamily α contain a Hyalin Repeat domain (HYR) in their N-termini while the domain is mostly absent or underrepresented in the other aGPCR subfamilies. The GAIN domain is not shown in this matrix. All the aGPCR NTFs analyzed are found in Supplementary File 19. **(B)** Phylogenetic distribution of protein domains in the N-termini of aGPCRs in opisthokonts. While non-holozoan aGPCRs do not possess conserved extracellular protein domains, most holozoan lineages, with the possible exception of ichthyosporeans, encode additional domains in their N-termini. Clear recruitment of ECDs is observed in the NTF of filasterean, choanoflagellate, and metazoan aGPCRs, with domains being conserved either in most of these lineages (light grey), in a subset of these lineages (e.g black for choanozoan-specific domains), or being lineage-specific (dark grey).

**Figure S8:**
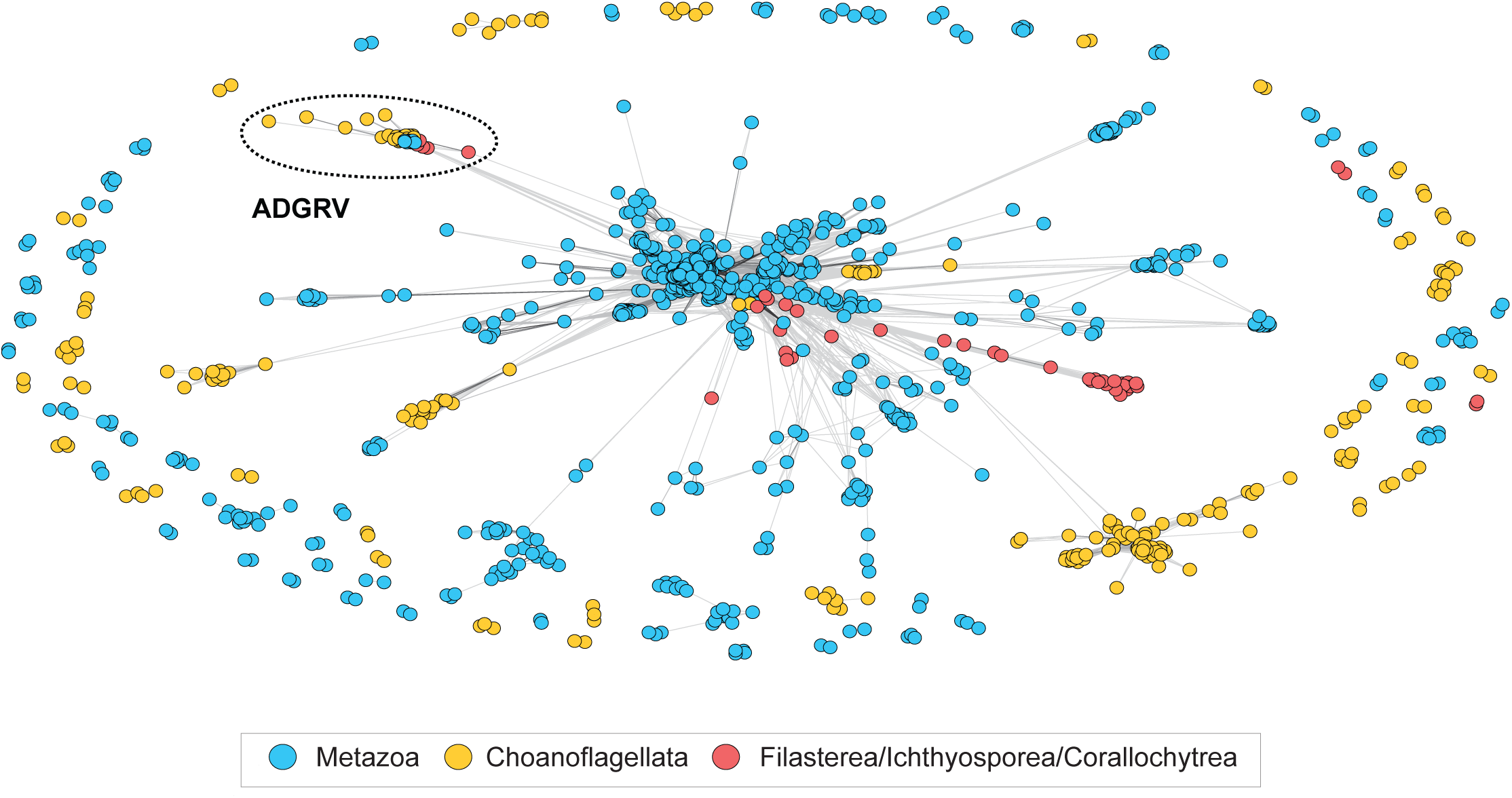
Independent diversification of most aGPCR NTFs in metazoans and CRMs. Pairwise similarity-based clustering analysis of the N-terminal fragment (NTF) of 1074 metazoan (blue), 337 choanoflagellate (orange), and 81 filasterean/ichthyosporean aGPCRs. The NTF of most metazoan aGPCRs cluster separately from choanoflagellate or other holozoan NTFs. A noticeable exception is observed for metazoan NTFs from the ADGRV family, which group with choanoflagellate and filasterean NTF sequences (delineated with a dotted oval). The choanoflagellate NTFs that cluster with metazoan ADGRV NTFs belong to the aGPCR subfamily θ, which we found to be orthologous to metazoan ADGRV based on their 7TM domain (**Fig. 5A**). Edges correspond to BLAST connections of P value <1e-20. All the NTF sequences used in the clustering analysis are available in Supplementary File 19.

**Figure S9:**
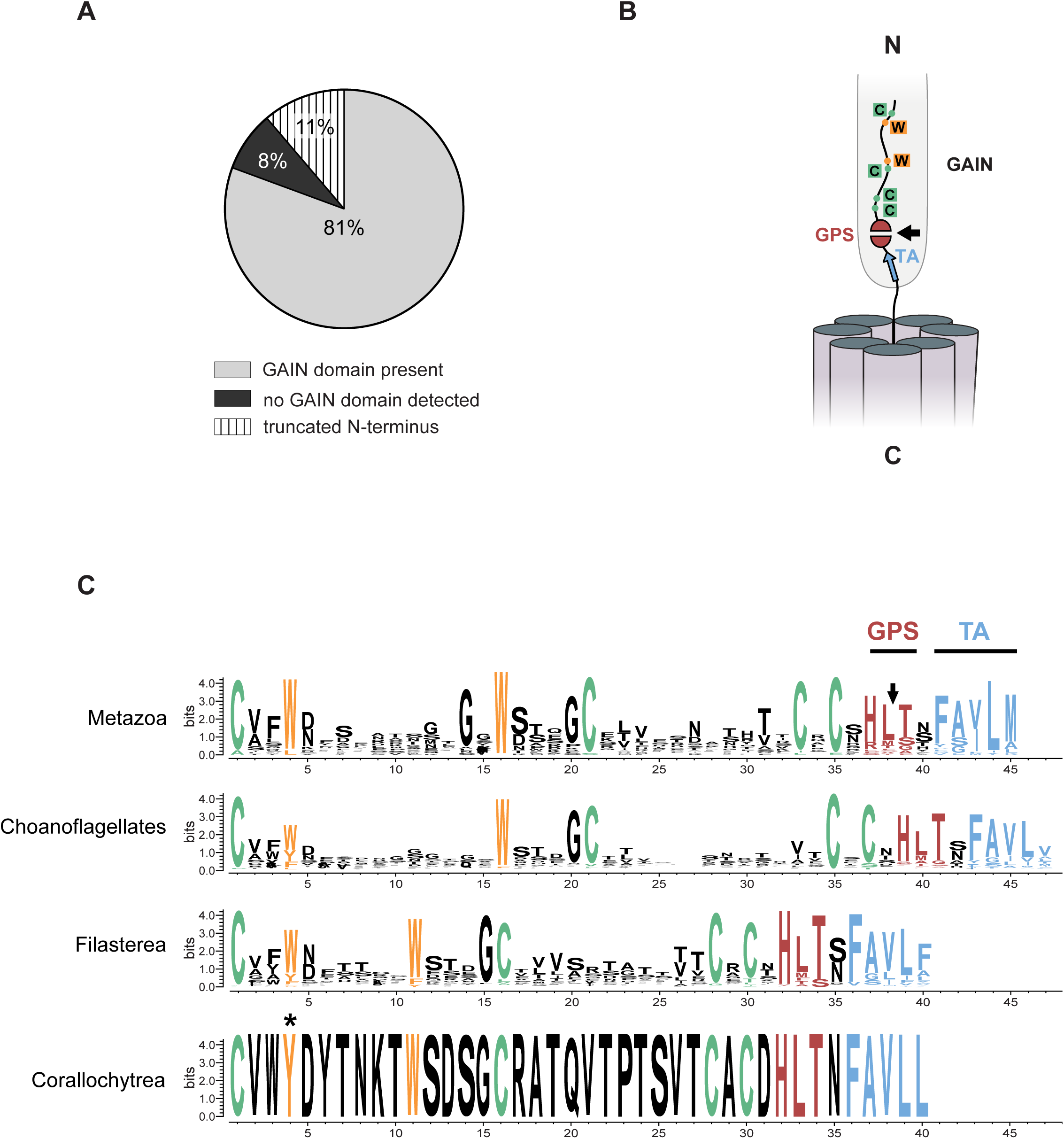
Conservation of the GAIN domain in metazoans, choanoflagellates, filastereans, and corallochytreans. **(A)** Pie chart illustrating the percentage of choanoflagellate aGPCRs exhibiting a GAIN domain (light grey, 81%), lacking a GAIN domain (dark grey, 8%), or presenting an incomplete N-terminus precluding us from testing for the presence of a GAIN domain (stripes, 11%). While a majority of choanoflagellate aGPCRs identified in our analysis possess a GAIN domain, we detected a few sequences with no apparent conservation of the fold (see Supplementary File 20). **(B)** Schematic of the 7TM-proximal GAIN domain, which extends into the extramembrane milieu. Highlighted are key tryptophans (orange Ws) and cysteines (green Cs) flanking the proteolytic GPS motif (red, cleavage site indicated with an arrow) and the hydrophobic tethered agonist element (TA, blue). **(C)** Sequence logos of the metazoan, choanoflagellate, filasterean, and corallochytrean C-terminal GAIN domains. The two canonical cysteines and tryptophans, important for the proper folding of the GAIN domain (Araç et al. 2012), are conserved in choanoflagellates, filastereans, and corallochytreans. Similarly, the autoproteolytic GPS motif and most of the TA consensus sequence, both required for the activation of aGPCRs, are conserved in all these clades (Prömel et al. 2012b; Barros-Álvarez et al. 2022). An alignment of the complete repertoire of aGPCRs (30 sequences) from *Mus musculus* was used to generate the metazoan logo; 301 aGPCR sequences were used for the choanoflagellate logo; and 89 aGPCR sequences were used for the filasterean logo. Because we recovered only one aGPCR sequence with a GAIN domain in corallochytreans, their logo has no real statistical value and is provided as a qualitative point of comparison. All the sequences used to create the logos are found in Supplementary File 21.

**Figure S10:**
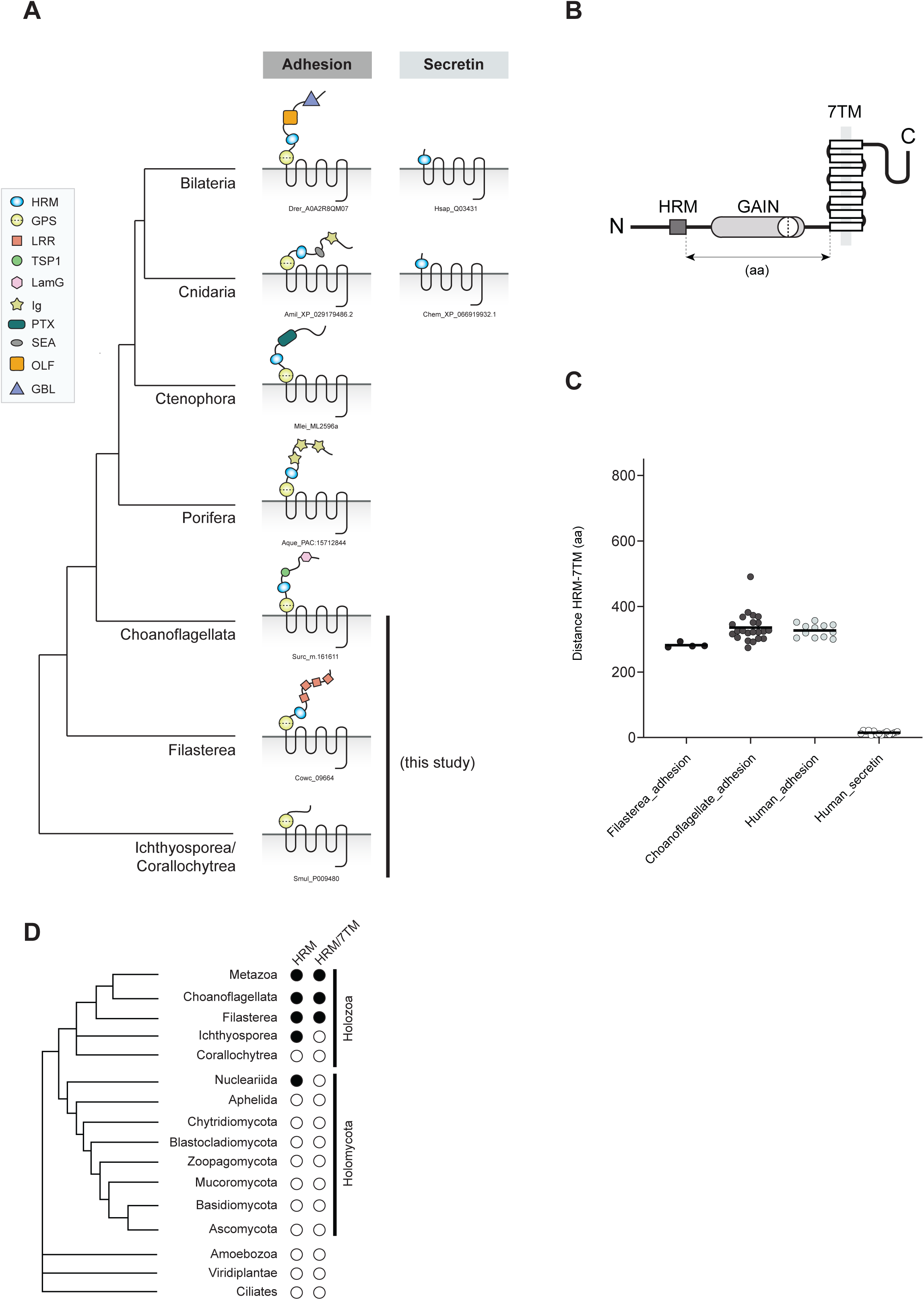
Pre-metazoan origin of the HRM domain and HRM-GAIN-7TM module. **(A)** Evolution of HRM-containing aGPCRs and Secretin GPCRs in Holozoa. Some aGPCRs and all Secretin GPCRs possess a Hormone Receptor Motif (HRM) domain (blue oval) in their extracellular region; the HRM domain is always found in combination with a GAIN domain (yellow circle) in aGPCRs while the GAIN domain is absent from the Secretin receptors. Secretin GPCRs likely evolved from aGPCRs (Nordström et al. 2009b; Scholz et al. 2019) and are only found in Cnidarians (this study) and in Bilaterians (Cardoso et al. 2024) (right). In contrast, HRM-containing aGPCRs are more ancient and were detected in metazoans, choanoflagellates, and filastereans (left). **(B)** Representative illustration of the extracellular region of HRM-containing aGPCRs. The GAIN domain (light grey) sits on the top of the 7TM. The HRM domain (dark grey rectangle) is always positioned N-terminal to the GAIN domain. The distance separating the C-terminal end of the HRM domain from the start of the 7TM (HRM-7TM distance) is depicted with a double-headed arrow and is assessed in panel (C). Diverse additional protein domains (ECDs), distal to the HRM/GAIN/7TM module, are often found in HRM-containing aGPCRs and are not depicted here. **(C)** Conserved HRM-7TM distance in filasterean, choanoflagellate, and human HRM-containing aGPCRs. We measured an average HRM-7TM distance of 282 aa, 335 aa, and 326 aa in HRM-containing filasterean, choanoflagellate, and human aGPCRs, respectively. In contrast, human Secretin GPCRs show an average HRM-7TM distance of 14 aa due to these receptors’ absence of the GAIN domain. Four filasterean, 22 choanoflagellate, 12 human HRM-containing aGPCRs, and 15 human Secretin GPCRs were assessed in this analysis. **(D)** Phylogenetic distribution of HRM and HRM/7TM module across eukaryotes. HRM is found in Holozoa and in the nucleariid *Fonticula alba*, one of the closest relatives of Fungi (Galindo et al. 2019). In contrast, the association of an HRM domain with 7TM (HRM/7TM) is only observed in holozoans – metazoans, choanoflagellates, and filastereans. The five HRM-containing proteins identified in *Fonticula alba* are provided in Supplementary File 22.

**Figure S11:**
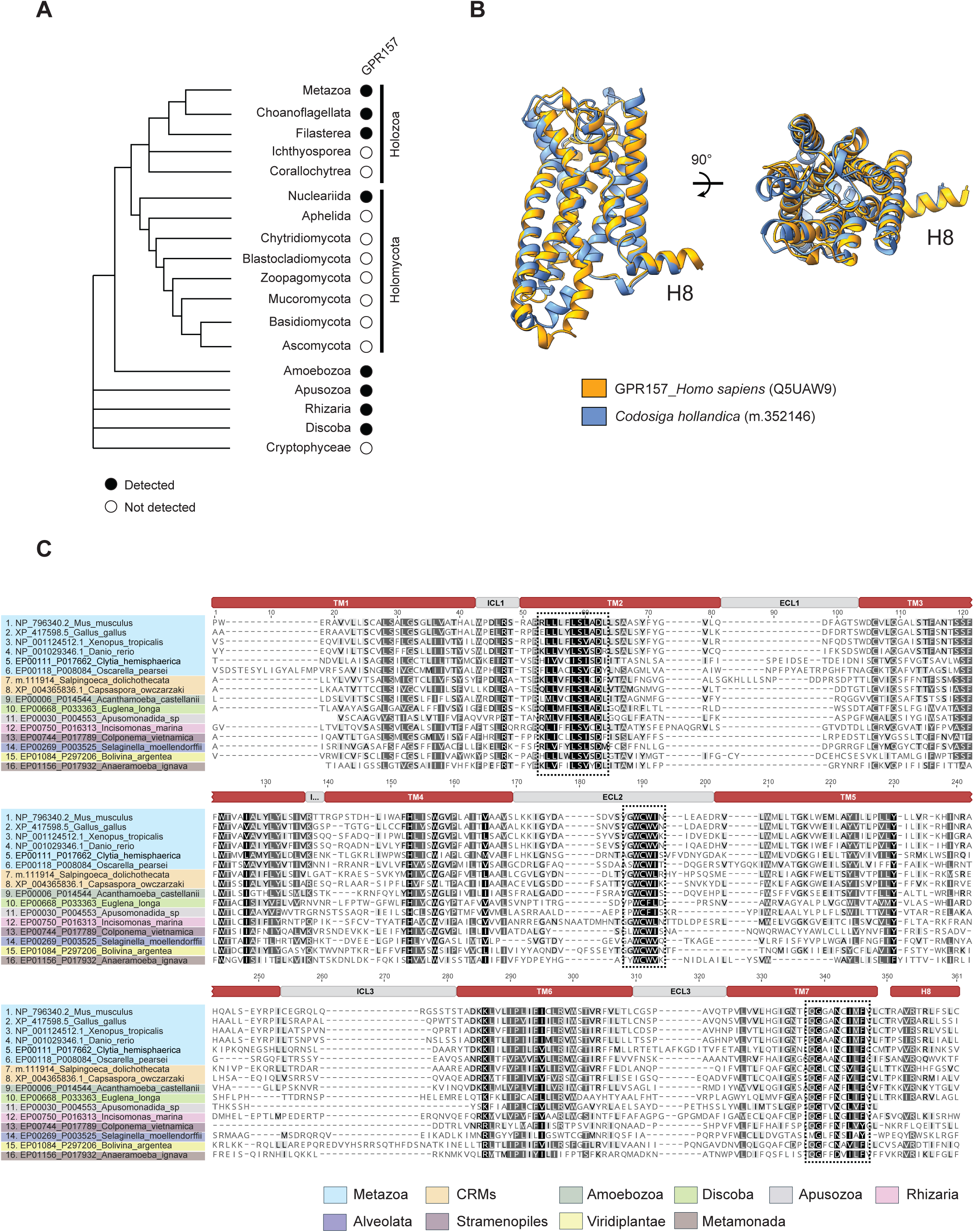
GPR157 is an ancient GPCR family conserved in eukaryotes. **(A)** Phylogenetic distribution of GPR157 across eukaryotes. GPR157 likely forms an ancient GPCR family in eukaryotes with homologs detected in Holozoa, Holomycota (only present in nucleariids), Amoebozoa, Apusozoa, Rhizaria, Discoba, and Cryptophyceae. Homologs were searched by BLASTing murine GPR157 (Q8C206) sequence against the entire dataset (993 species) of the EUKPROT v3 BLAST server (E-value: 1e^-5^) (https://evocellbio.com/eukprot/; (Richter et al. 2022)). We defined *bona fide* GPR157 hits as those with at least 70% query coverage and at least 30% sequence identity. All EUKPROT blast hit sequences are listed in Supplementary File 22. **(B)** Predicted structural homology of the 7TM region of *H.sapiens*_GPR157_Q5UAW9 (orange) and *C.hollandica*_m.352146 (blue) receptors. An additional helix 8 (H8) is predicted in both metazoan and choanoflagellate GPR157. All structural models shown here have a confidence score >70 pLDDT. View of the superimposed models from the plane of the membrane (left) and the top (right). **(C)** Multi-alignment of the 7TM/H8 domain of GPR157 from various eukaryotes reveals motif conservation, including a “LLxxLSL/V” motif in transmembrane helix 2 (TM2) and a “WCWI/V” motif in the extracellular loop 2 (ECL2) (dotted boxes).

**Figure S12:**
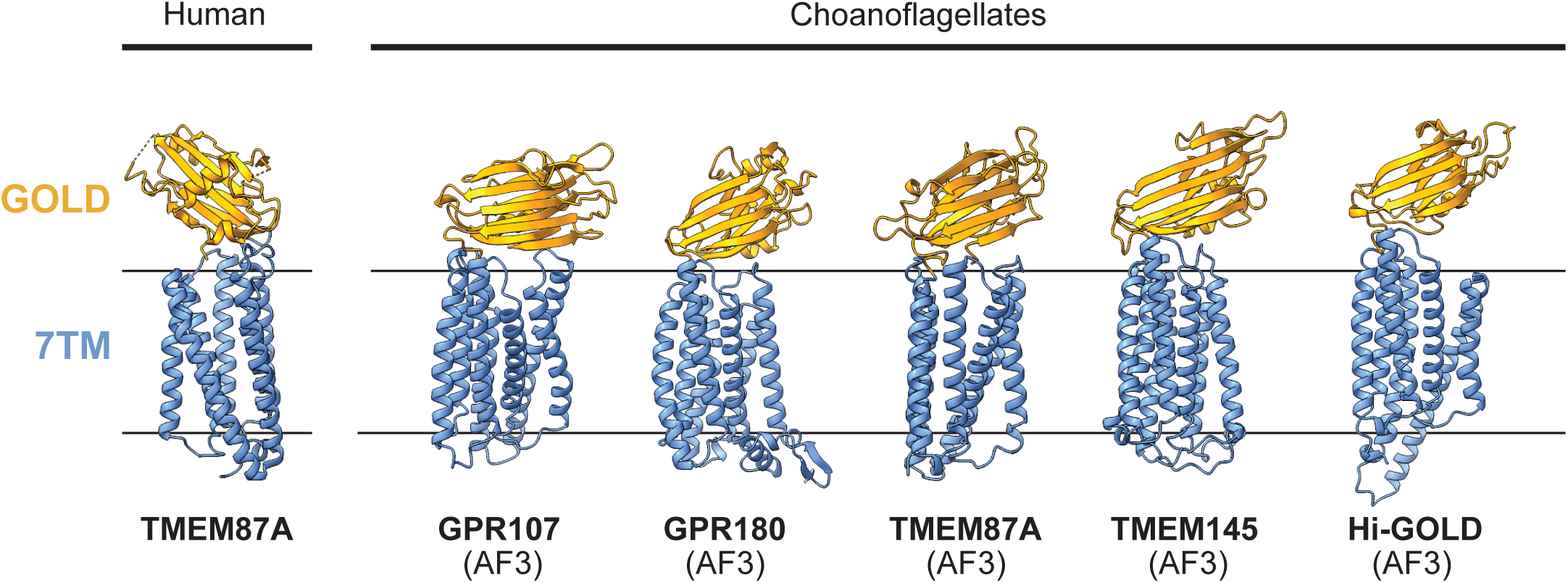
Predicted structural similarities among metazoan and choanoflagellate GOST family receptors. Experimentally determined human TMEM87A structure (PDB:8CTJ; (Hoel et al. 2022)) and AlphaFold3 (AF3) predicted structures of representative proteins from the five GOST subfamilies detected in choanoflagellates (GPR107/108, GPR180, TMEM87A, TMEM145, and the newly identified Hi-GOLD) shown as cartoons, viewed from the plane of the membrane. Low confidence (<70 pLDDT) regions of predicted structures have been removed. Both metazoan and choanoflagellate GOST GPCRs exhibit a GOLD domain (orange) on top of a 7TM domain (blue).

**Figure S13:**
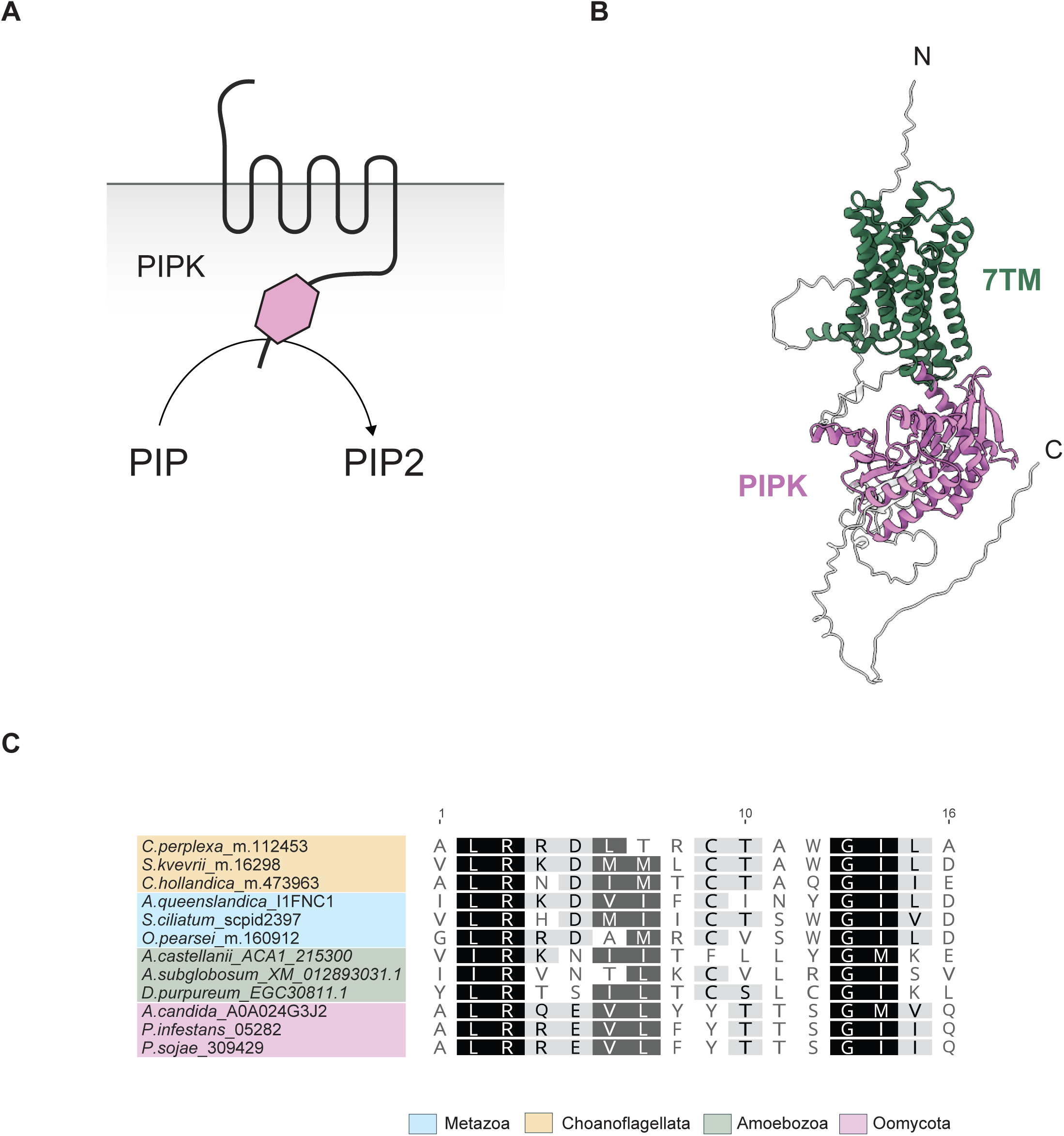
Features of choanoflagellate GPCR PIPKs. **(A)** A schematic depiction of a GPCR PIPK. GPCR PIPKs present a canonical 7TM domain combined with a phosphatidylinositol phosphate kinase (PIPK; pink hexagon) at the C-terminus (Van Den Hoogen et al. 2018; van den Hoogen and Govers 2018). While experimental evidence is lacking, the PIPK domain of GPCR PIPK receptors is likely to signal via the production of phospholipid-based second messengers (e.g PIP2). **(B)** Alphafold-predicted structure of a choanoflagellate GPCR PIPK (*S.urceolata*_m_147488) with the 7TM and PIPK domains depicted in green and pink respectively. Regions with a low prediction score (<70 pLDDT) are depicted in white. **(C)** Multiple sequence alignment showing the conservation in choanoflagellates of the diagnostic GPCR PIPK motif “LR(x)_9_GI” in the linker region separating the 7TM domain from the PIPK domain.

**Figure S14:**
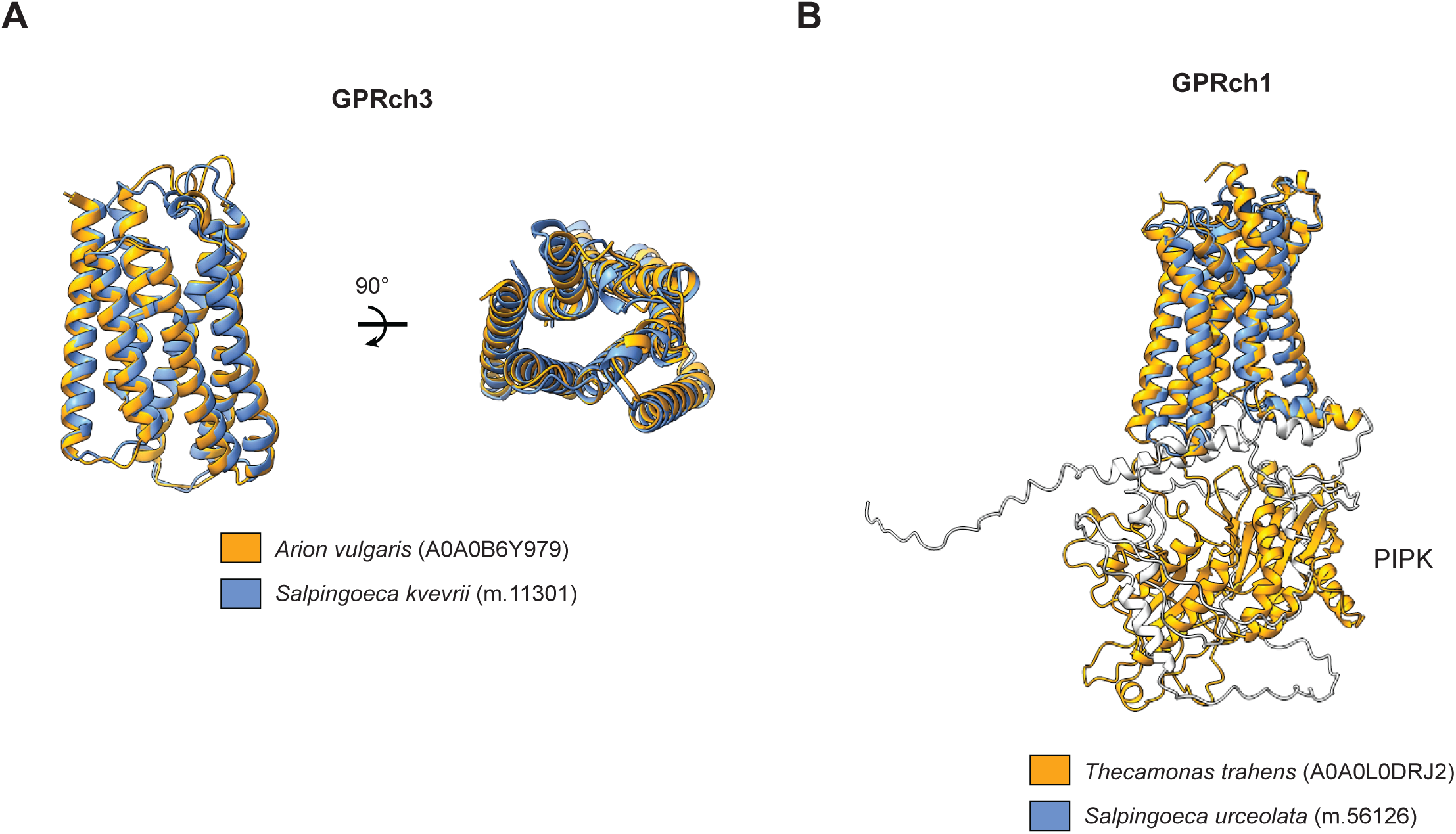
Structural features of GPRch3 and GPRch1 GPCRs. **(A)** Predicted structural homology of the 7TM domain from choanoflagellate GPRch3 GPCR *S.kvevrii*_m.11301 (orange) and its top structural metazoan hit *A.vulgaris*_A0A0B6Y979 (blue) (E-value: 4.64e^-13^). View of the superimposed models from the plane of the membrane (left) and the top (right). All structural models shown here have a confidence score >70 pLDDT. **(B)** Superimposed structures of full-length choanoflagellate GPRch1 GPCR *S.urceolata*_m.56126 (blue) with its top structural hit *T.trahens*_A0A0L0DRJ2 (orange) (E-value: 5.92e^-10^). While ciliates, along with other non-choanozoan eukaryotes that encode GPRch1 GPCRs, possess an additional PIPK domain in the C-terminal region of the GPCR, no GPRch1 GPCRs appeared to have this additional cytosolic enzymatic domain in choanoflagellates. Regions with a low prediction score (<70 pLDDT) are colored in white.

**Figure S15:**
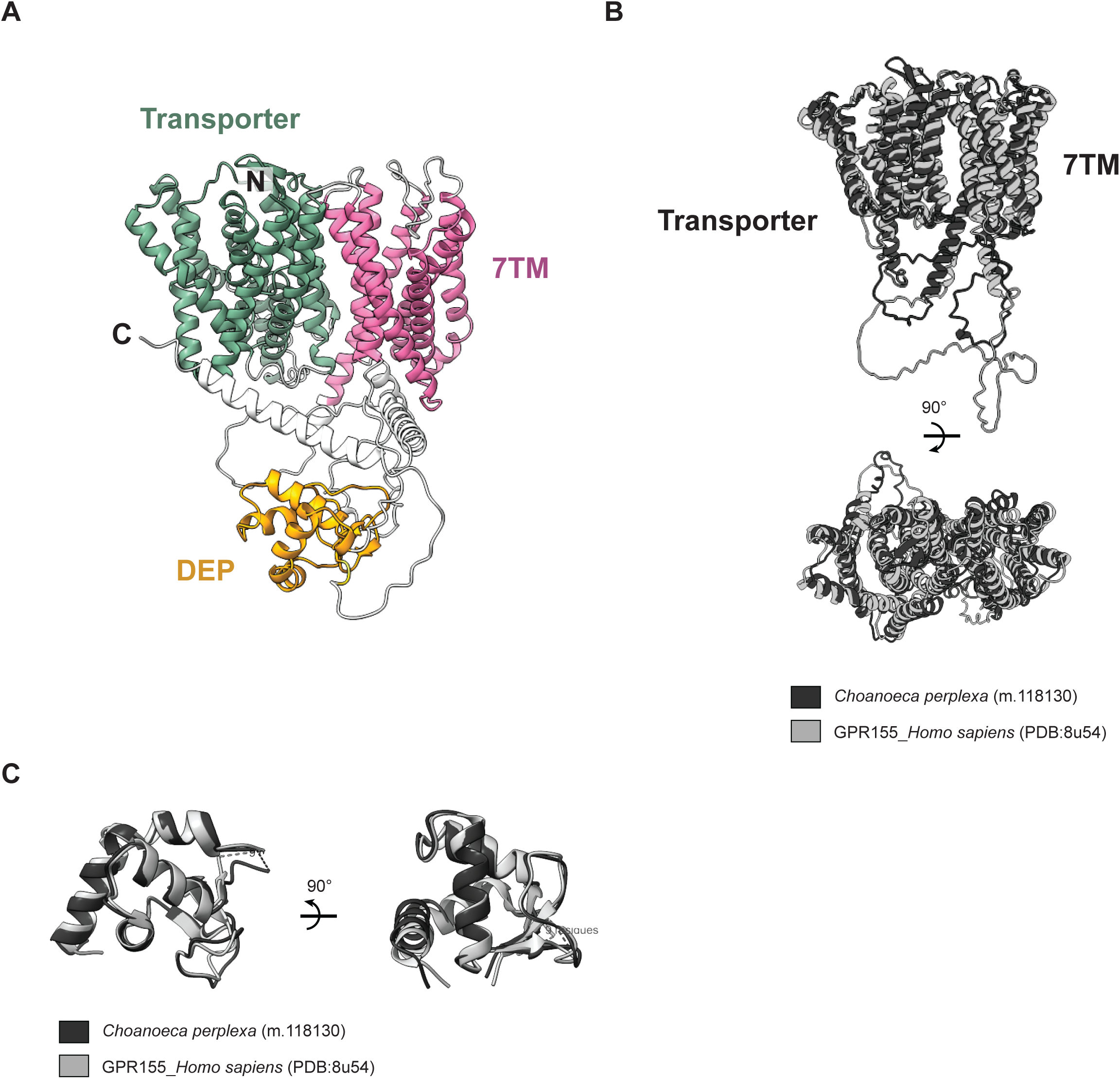
Conservation of metazoan GPR155 in choanoflagellates. **(A)** Predicted structure of a full-length choanoflagellate GPR155 receptor (*C. perplexa*_m.118130). Similar to metazoan GPR155 (Shin et al. 2022; Bayly-Jones et al. 2024)), choanoflagellate GPR155 possesses a 7TM domain (pink) fused to a transporter domain (green), and an additional Dishevelled, EGL-10 and Pleckstrin (DEP) domain (orange) in the C-terminal region. Regions with a low prediction score (<70 pLDDT) are depicted in white. **(B)** Structural homology of the Transporter/7TM module from the alphafold-predicted choanoflagellate GPR155 structure (*C. perplexa*_m.118130; dark grey) and the experimentally solved human GPR155 structure (PDB: 8u54; light grey). View of the superimposed models from the plane of the membrane (top) and the top (bottom). **(C)** Superimposed structures of the DEP domain from choanoflagellate (*C. perplexa*_m.118130; dark grey) and human (PDB: 8u54; light grey) GPR155 GPCRs. The choanoflagellate DEP model has a confidence score >70 pLDDT.

**Figure S16:**
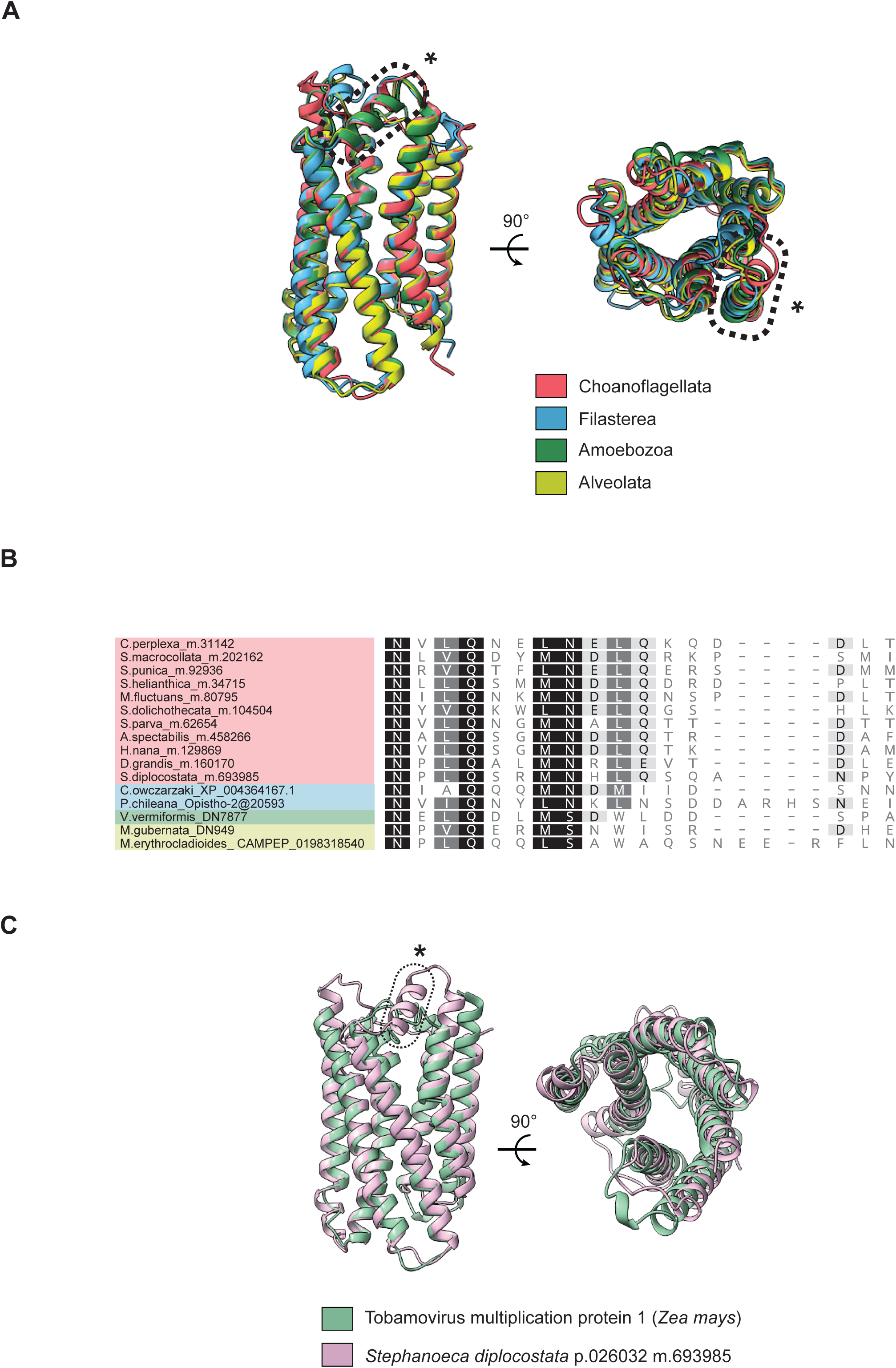
Structural features of the newly established RSF GPCR family. **(A)** Predicted structural homology of the RSF 7TM domain from *S. diplocostata*_m.693985 (red), *C. owczarzaki*_P006019 (blue), *M.gubernata*_dn949 (yellow), and *V. vermiformis*_dn7877 (green). A conserved short helix between TM6 and TM7 is predicted in all the structures depicted (dotted box). View of the superimposed models from the plane of the membrane (left) and the top (right). All structural models shown here have a confidence score >70 pLDDT. **(B)** A multi-alignment of sequences corresponding to the short extra helix depicted in (A), revealed a NxLQxxMN” motif conserved across eukaryotes. **(C)** Mild structural similarities are shared between predicted *S. diplocostata*_m693985 (pink) and top structural Foldseek hit *Z. mays*_TOM1_B7ZYE1 (green) (E-value:1.66e^-5^). The short extra helix (dotted box) is not observed in TOM1 GPCRs. View of the superimposed models from the plane of the membrane (left) and the top (right).

**Figure S17:**
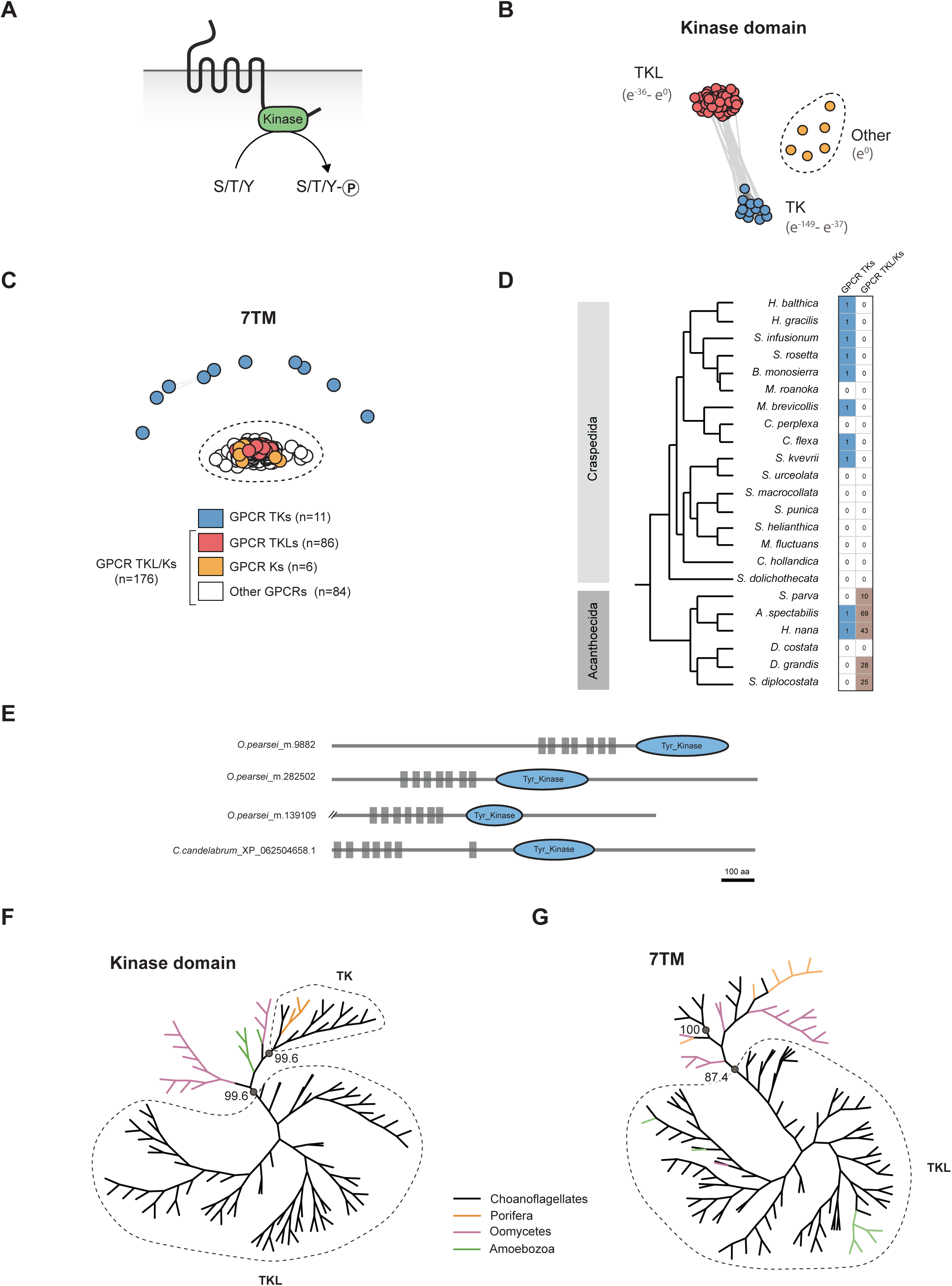
**Features and evolution of choanoflagellate GPCR TK/TKL/Ks**. **(A)** A schematic depiction of a GPCR Kinase. GPCR Kinases present a canonical 7TM domain combined with a kinase domain (green oval) in their C-terminus (Van Den Hoogen et al. 2018; van den Hoogen and Govers 2018). The kinase domain possibly phosphorylates the GPCR itself or other proteins (van den Hoogen and Govers 2018). **(B)** All-against-all pairwise comparison of the catalytic domain of choanoflagellate GPCR Kinases reveals distinct kinase domain populations with 11 tyrosine kinase (TK) domains grouping together (blue) and sharing sequence similarities with a second cluster encompassing 86 tyrosine kinase-like (TKL); 6 additional domains (orange) do not share sequence similarities with any other kinase domains in this dataset (Other) (Supplementary Files 25 and 26). Each kinase domain (circle) was blasted against a local version of KinBase, the curated protein kinase dataset from www.kinase.com (Manning et al. 2002; Bradham et al. 2006; Goldberg et al. 2006)), for identification (Supplementary File 26). The E-values depicted summarize the lowest and highest e-values associated with top blast hits obtained for members of each cluster. While choanoflagellate GPCR TKs systematically recovered metazoan tyrosine kinases as their top BLAST hits, choanoflagellate GPCR TKL preferentially matched with amoebozoan tyrosine kinase-like proteins. No BLAST hits were recovered for the 6 isolated kinase domains. Connecting lines correspond to pairwise BLAST scores of p-value <1e^-20^. **(C)** Distinct families of GPCR Kinases are found in choanoflagellates. All-against-all pairwise comparison of the 7TM domain of choanoflagellate GPCR Kinases revealed that they cluster into different families, with the 86 GPCR TKL receptors (red) previously identified in **(B)** grouping with the 6 GPCR K (orange), and 84 additional choanoflagellate GPCRs that do not possess a C-terminal kinase domain (white), forming a cluster of 176 GPCRs in total (dotted line; Fig. 1). On the opposite, the 11 GPCR TKs (blue) previously identified in **(B)** grouped independently from the GPCR TKL/Ks and were either single GPCRs or grouped with other GPCR TKs. Connecting lines correspond to pairwise BLAST scores of p-value <1e^-6^. The 7TM sequences analyzed and the output of the clustering analysis are found in Supplementary Files 28 and 29. **(D)** Phylogenetic distribution of GPCR TKs (blue) and GPCR TKL/Ks (brown) within the choanoflagellate phylogeny. While GPCR TKs are found in a large range of choanoflagellate species, including both craspedids and acanthoecids, GPCR TKL/Ks appear to be restricted to acanthoecids (Fig. 2). **(E)** Schematic representation of selected sponge GPCR TKs recovered in our analysis by blasting the kinase domain of choanoflagellate GPCR kinases against the EUKPROT database (Richter et al. 2022).GPCRs are depicted at scale. Scale bar = 100 aa. **(F and G)** Phylogenetic analyses of the kinase **(F)** and 7TM domains **(G)** from diverse GPCR TKL/Ks and GPCR TKs. **(F)** The TKL domains of choanoflagellate GPCR TKLs form a well-supported monophyletic clade to the exclusion of the kinase domains of other GPCR kinases (Bootstrap support 99.6%). The TK domains of choanoflagellate and sponge GPCR-TKs form a well-supported clade (Bootstrap support 99.6%). **(G)** The 7TM domains of GPCR TKL/Ks cluster with amoebozoan GPCR TKL/Ks (Bootstrap support 87.4%), to the exclusion of 7TMs from other GPCR kinases, including GPCR TKs. All the extracted Kinase and 7TM sequences used to build the phylogenies in Fig. S17F and G, as well as the resulting trees are listed in Supplementary files 30, 31, 32, and 33.

**Figure S18:**
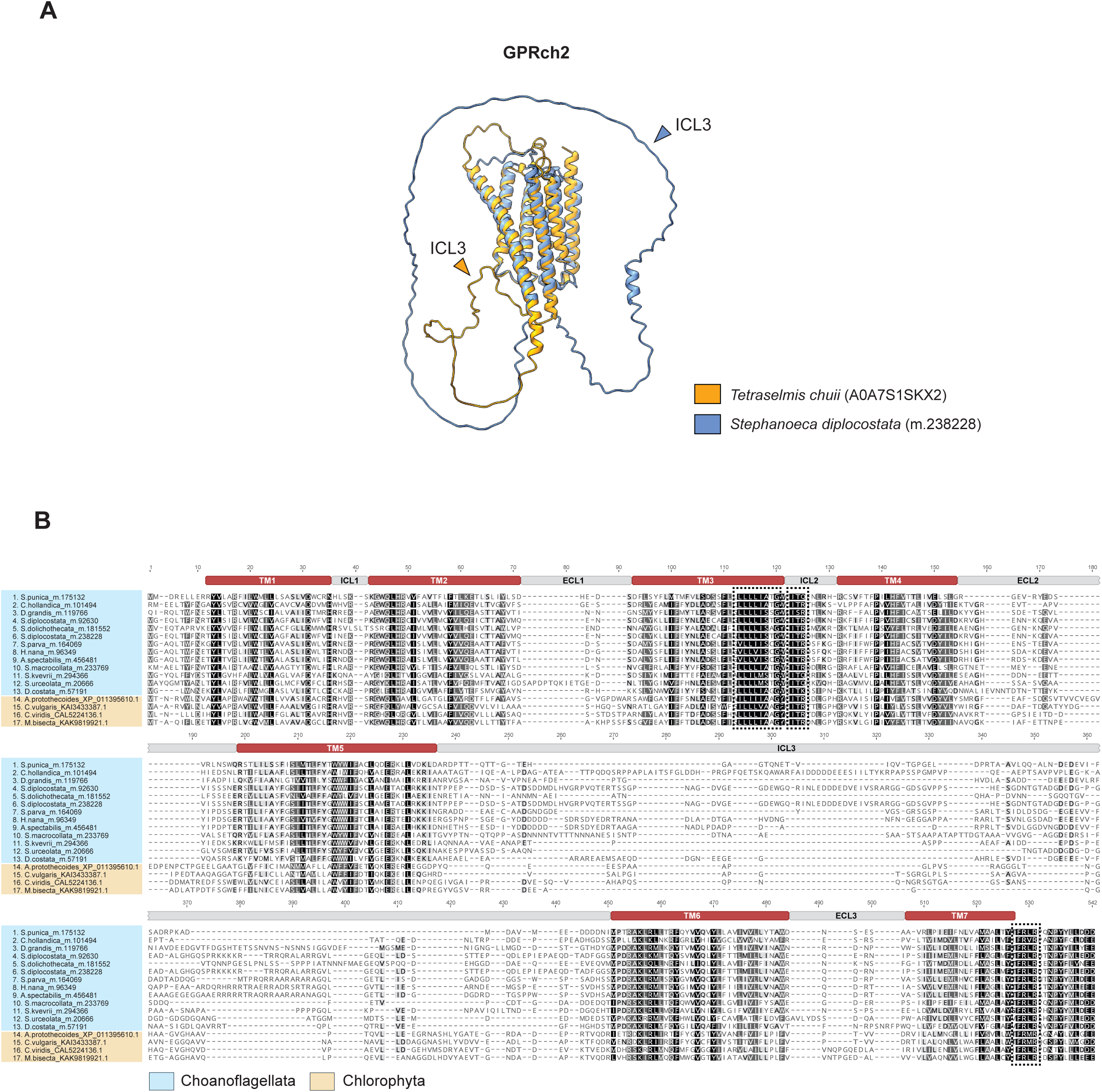
Structural features of GPRch2 GPCRs. **(A)** Predicted structural homology of the 7TM domain from choanoflagellate GPRch2 GPCR *S.diplocostata*_m.238228 (blue) and its top structural chlorophyte hit *T.chuii*_A0A7S1SKX2 (orange) (E-value: 6.83e^-12^). A large intracellular loop 3 (ICL3) between TM5 and TM6 (indicated with an arrowhead) is observed in choanoflagellate and, to a lesser extent, in chlorophyte GPRch2 GPCRs. ICL3 is unstructured with a low confidence score found in both models (<70 pLDDT). **(B)** Multiple sequence alignment of the 7TM region of GPRch2 GPCRs from choanoflagellates and chlorophytes revealed a conserved “LLLLIS/AxG” motif in TM3, an “ITR” motif in ICL2, and a “FRLRxxNPY” motif in the C-terminal region (dotted boxes).

**Figure S19:**
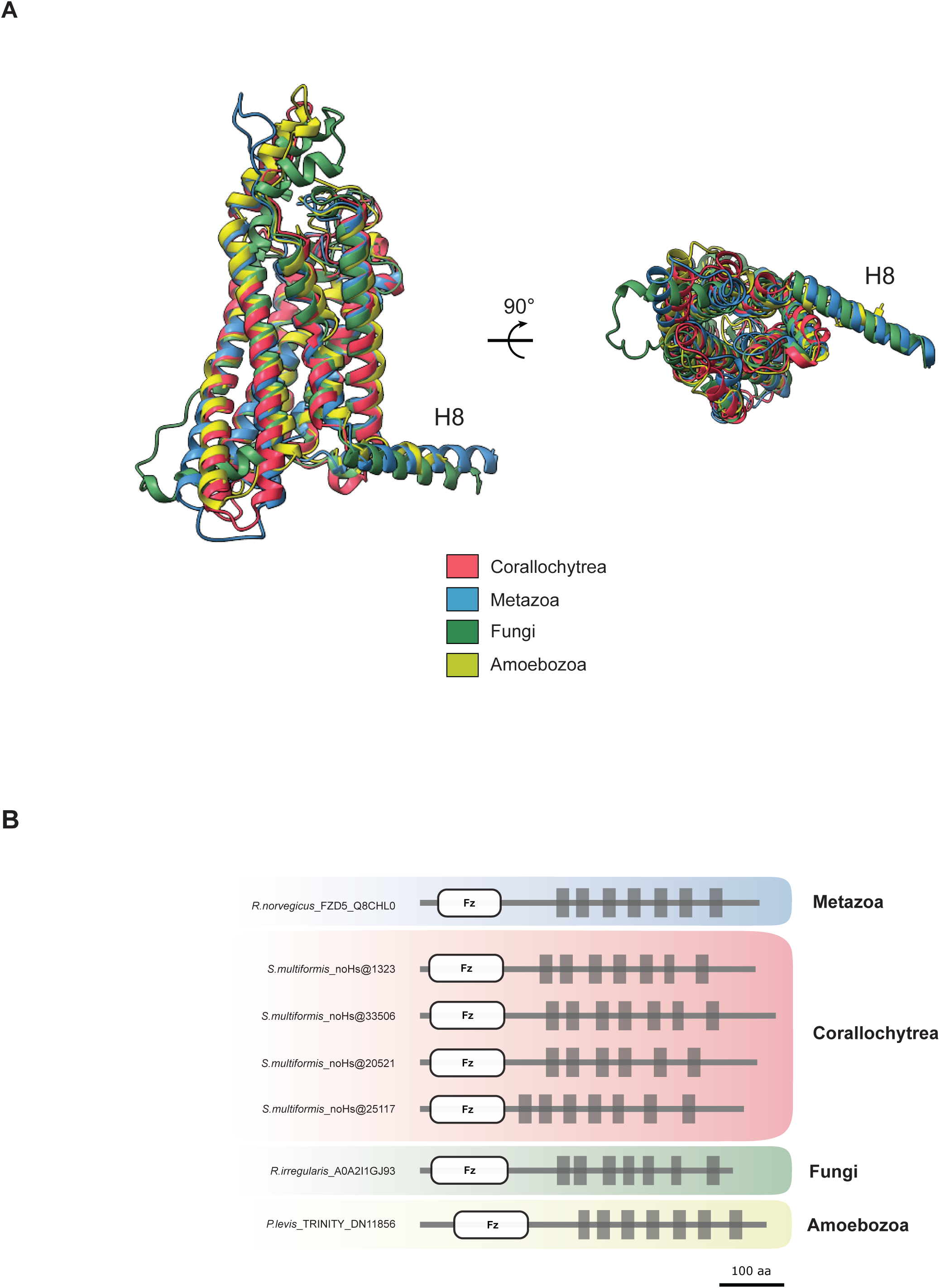
Metazoan Frizzled/Smoothened homologs detected in corallochytreans, fungi, and amoebozoans. **(A)** Predicted structural similarities of the 7TM region of *S.multiformis*_p007117_colp12_trinity150504 (red), *R.norvegicus*_Q8CHL0 (blue), *R.irregularis*_a0a2i1gj93 (green), and *P.levis*_p001581_dn11856 (yellow) Frizzled/Smoothened receptors. An additional helix 8 (H8) C-terminal to the 7TM domain is also depicted. All predicted regions shown here have a confidence score >70 pLDDT. View of the superimposed models from the plane of the membrane (left) and the top (right). **(B)** Schematic representation of Frizzled/Smoothened GPCRs found in metazoans, corallochytreans, fungi, and amoebozoans. The structural homology extends beyond their 7TMs as a canonical Frizzled domain (Fz) is detected in the N-termini of all these receptors. The Frizzled/Smoothened homologs detected in *S.multiformis* are listed in Supplementary File 34. GPCRs are depicted at scale. Scale bar = 100 aa.

